## Supporting Information for "Profiling Cytosolic Drug Delivery in Mammalian Cells: A Generalizable Assay for Intracellular Accumulation"

### TABLE OF CONTENTS

#### SUPPLEMENTARY FIGURES


| | | |
|---|---|---|
| **Figure S1** | Fluorescent microscopy images showing Wild type HeLa and HaloTag HeLa (HT HeLa) cells in presence and absence of TMRcl. | S4 |
| **Figure S2** | SDS-PAGE analysis of cells treated with TMRcl, including a no-TMRcl control. | S5 |
| **Figure S3** | Concentration dependent response of DBCOcl treatment | S6 |
| **Figure S4** | Concentration dependent competition with DBCOcl using CAPA | S7 |
| **Figure S5** | Time dependent response of DBCOcl treatment | S8 |
| **Figure S6** | Investigation of optimum concentration scan of TMRaz | S9 |
| **Figure S7** | Investigation of optimum incubation time with TMRaz | S10 |
| **Figure S8** | Chemical Structure of the library of Polyarginines | S11 |
| **Figure S9** | Membrane permeabilization assay using 40ug/mL of digitonin | S12 |
| **Figure S10** | Stereochemistry effect on polyarginine accumulation in complete media | S13 |
| **Figure S11** | Apparent accumulation of the *N*-alkylation library with varying degree of backbone *N*-alkylation | S14 |
| **Figure S12** | Apparent accumulation of the *N*-alkylation library with varying positions of backbone *N*-alkylation | S15 |


#### BIOLOGICAL METHODS


| | |
|---|---|
| Reagents | S16 |
| Cell culture | S16 |
| One-step assay with Tamra chloroalkane (TMRcl). | S16 |
| CHloroalkane Azide Membrane Penetration (CHAMP) assay | S17 |
| Confocal fluorescence Microscopy | S17 |
| Membrane permeabilization assay | S17 |
| +36GFP amino acid sequence | S18 |
| Isolation and purification of +36GFP and Wild Type GFP | S18 |


#### SYNTHESIS AND CHARACTERIZATION OF PEPTIDES, SMALL MOLECULES, AND ANTIBIOTICS


| | |
|---|---|
| Synthesis and characterization of DBCOcl | S21 |
| Synthesis of Polyarginine peptides (R5az, R7az, R9az, R11az) | S30 |

| | |
|---|---|
| Synthesis of the macrocyclization peptide library | S32 |
| Synthesis of the *N*-methylation and *N*-alkylation peptide libraries | S34 |
| Synthesis and characterization of 1p and 1n | S43 |
| Synthesis and characterization of 2p and 2n | S53 |
| Synthesis and characterization of 3p and 3n | S57 |
| Synthesis and characterization of 4a-4e | S79 |
| Synthesis and characterization of 5a-5e | S108 |
| Synthesis and characterization of **vanAZ1** | S112 |
| Synthesis and characterization of **rifAZ1** | S115 |
| Synthesis and characterization of **triAZ1** | S117 |
| Synthesis and characterization of **moxAZ1** | S123 |
| Synthesis and characterization of **novAZ1** | S128 |
| Synthesis and characterization of **purAZ1** | S133 |
| Synthesis and characterization of **zolAZ1** | S137 |
| Synthesis and characterization of **zolAZ2** | S139 |
| Synthesis and characterization of **zolAZ3** | S141 |
| Synthesis and characterization of **metAZ1** | S143 |
| Synthesis and characterization of **sulAZ1** | S147 |
| 404 Library of azido-tagged molecules (Enamine/Sigma) | S150 |
| **REFERENCES** | S151 |

### SUPPLEMENTARY FIGURES

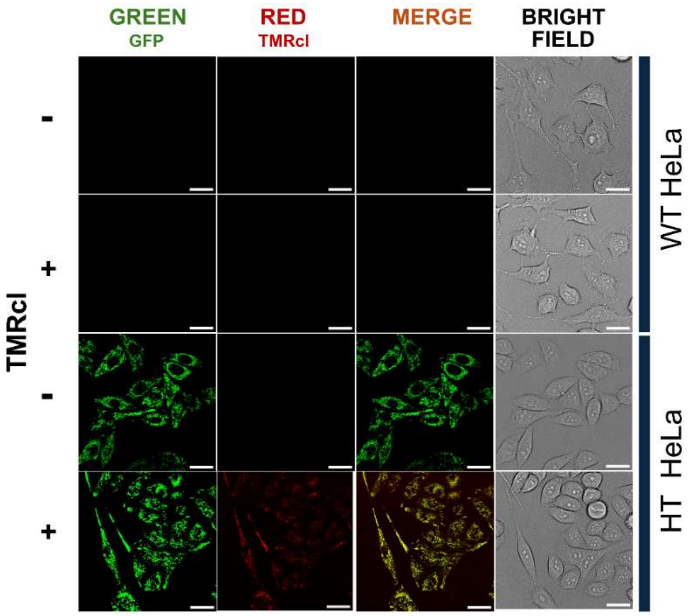


**Figure S1**. Fluorescent microscopy images showing “pulse-chase” experiments with cells pulsed with DBCOcl and chased with TMRcl. WT HeLa and HT HeLa cells not treated with DBCOcl were used as negative control. The scale bar represents 25 μM.

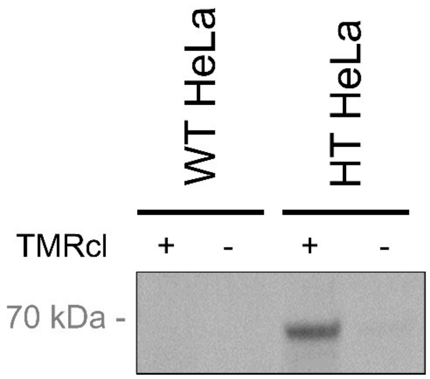


**Figure S2**. SDS-PAGE analysis of cells treated with TMRcl, including a no-TMRcl control. WT HeLa cells were used as a negative control.

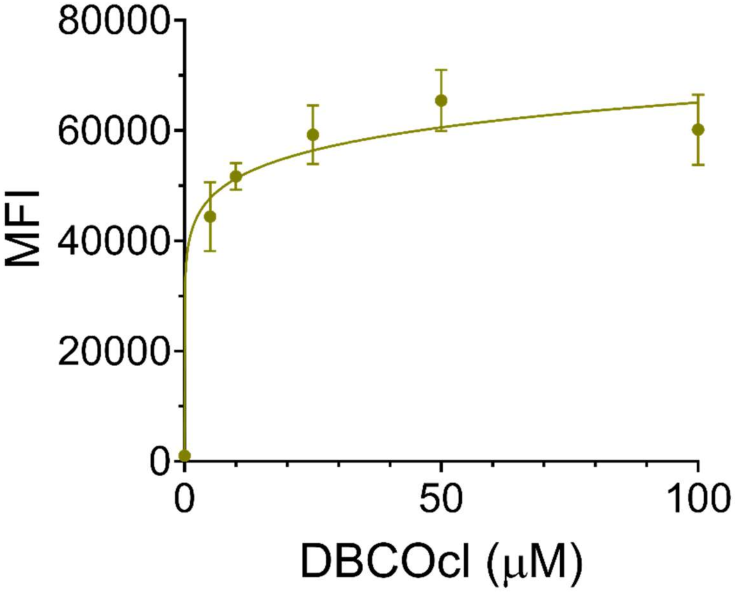


**Figure S3.** Concentration scan of DBCOcl. HT HeLa cells are treated with different concentrations of DBCOcl followed by treatment with 50 μM TMRaz. Data are represented as mean ± SD (n= 3).

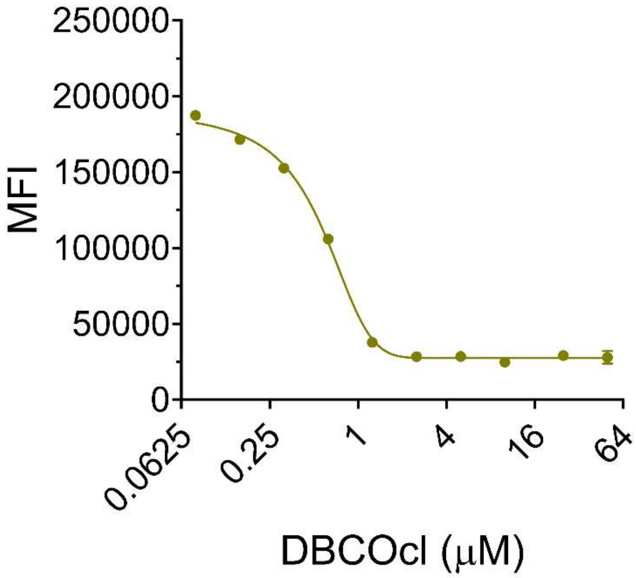


**Figure S4.** Concentration scan of DBCOcl using the CAPA assay. HT HeLa cells were pulsed with different concentrations of DBCOcl and chased with TMRcl at 50 μM. Data are represented as mean ± SD (n= 3).

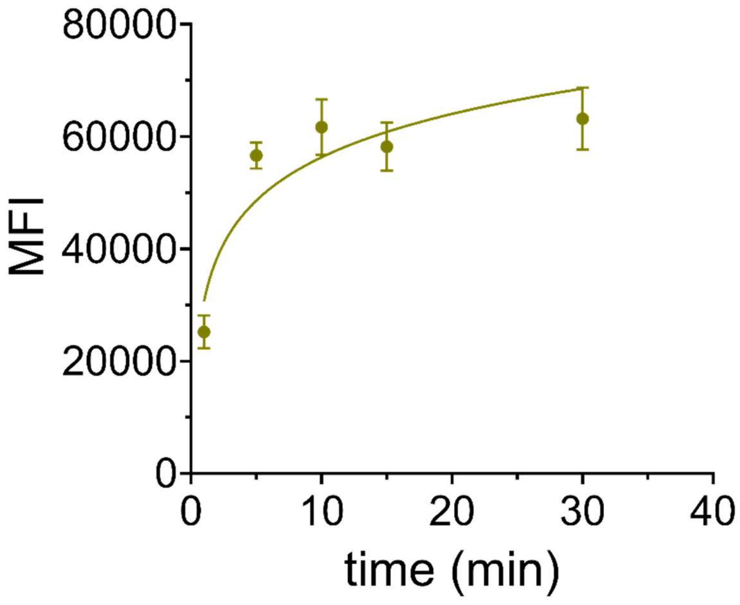


**Figure S5**. Kinetics of DBCOcl covalent attachment to HaloTag in HT HeLa cells at five different time points (0, 1, 5, 10, 15, 30 minutes). Data are represented as mean ± SD (n= 3).

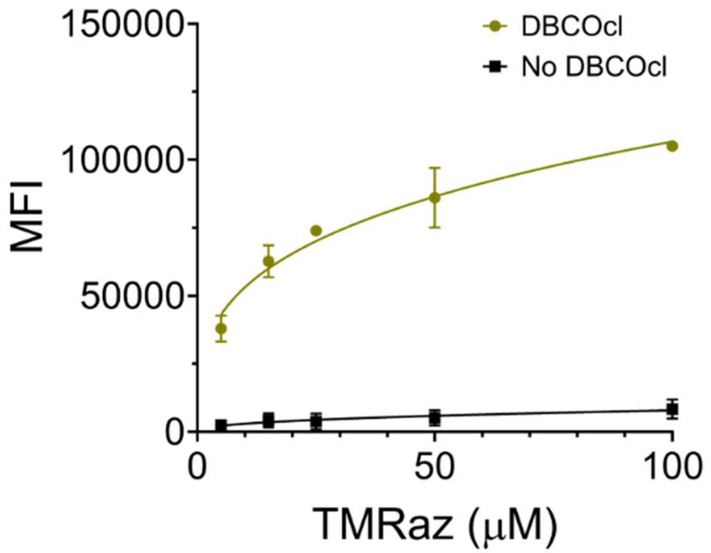


**Figure S6**. Investigation of optimum incubation time with TMRaz in HT HeLa cells at five different time points (0, 1, 5, 10, 15, 30 minutes). Data are represented as mean ± SD (n= 3).

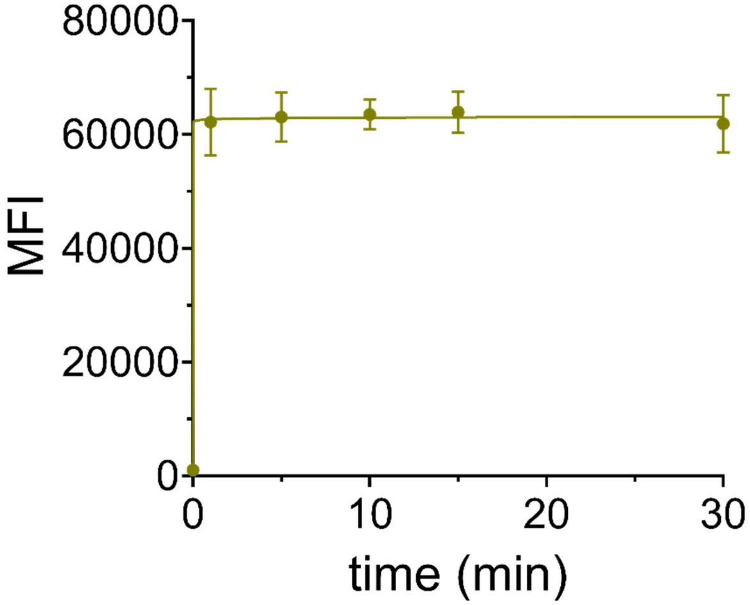


**Figure S7**. Investigation of optimum concentration scan with TMRaz in HT cells at five different concentrations (5,15, 25, 50, 100 μM). Data are represented as mean ± SD (n= 3).

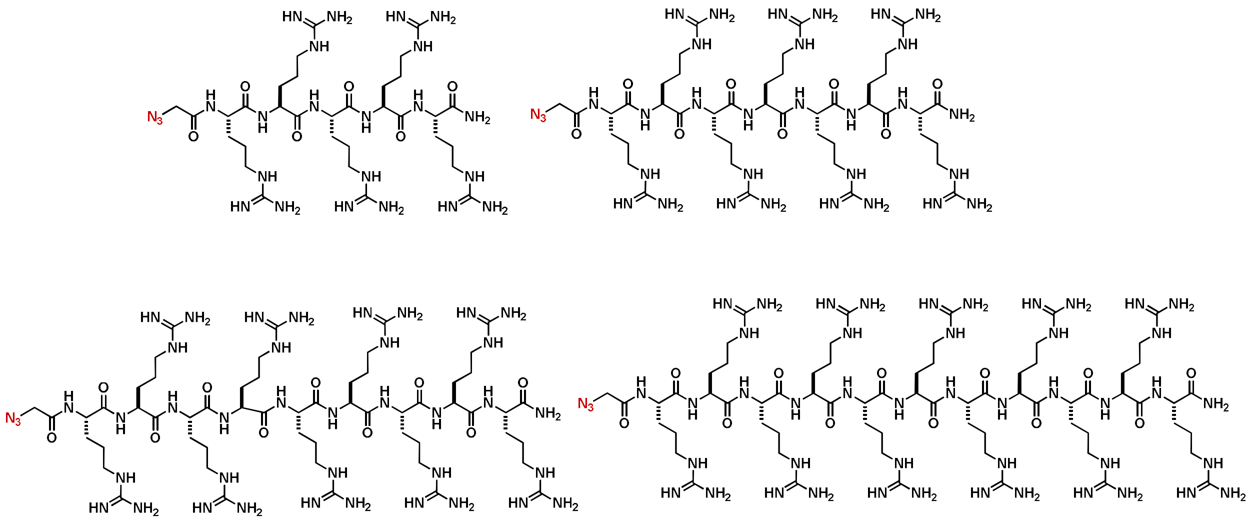

**Figure S8:** Chemical Structures of azide tagged polyarginines. R5az, R7az, R9az, and R11az (from left to right)

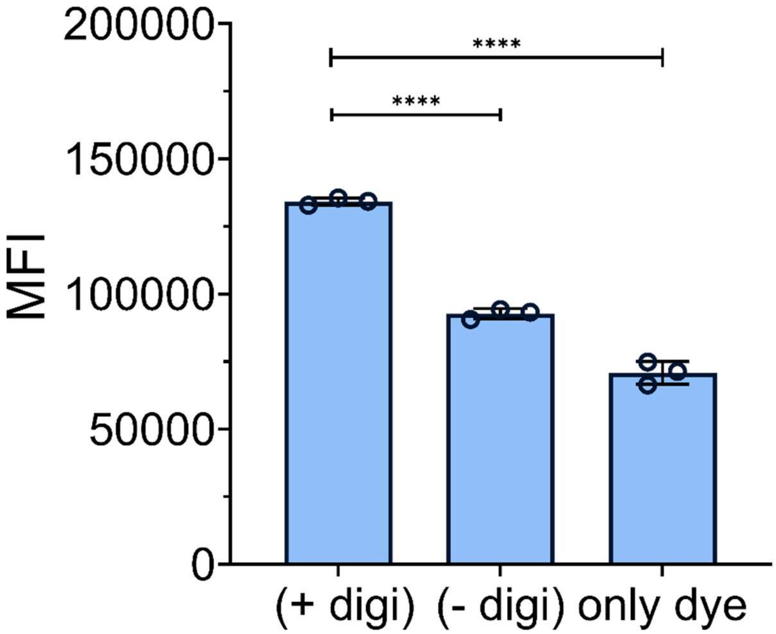


**Figure S9.** Membrane permeabilization assay using 40 ug/mL of digitonin. DBCOcl-treated cells were incubated with digitonin along with the membrane impermeable dye, Sulfo-cyanine 5 for 1 h. Cells treated with dye only were used to show the background signal. Data are represented as mean ± SD (n= 3).

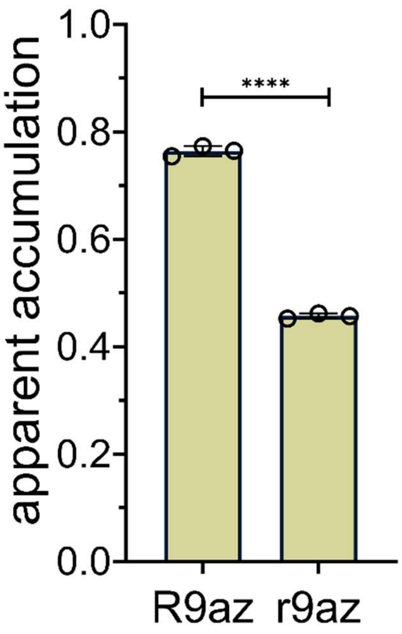


**Figure S10.** Stereochemistry effects on polyarginine accumulation in complete media. Data are represented as mean ± SD (n= 3).

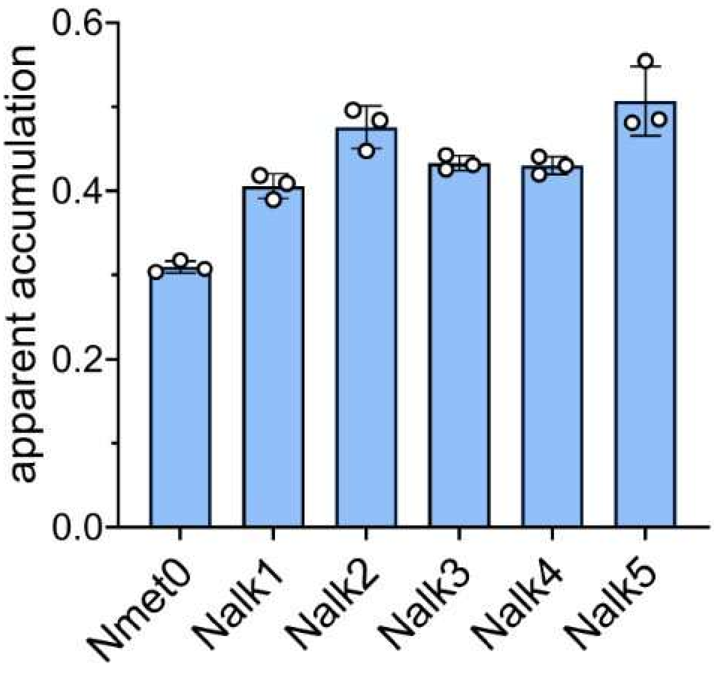


**Figure S11.** The apparent accumulation of the *N*-alkylation library with varying degrees of backbone *N*-alkylation. Data are represented as mean ± SD (n= 3).

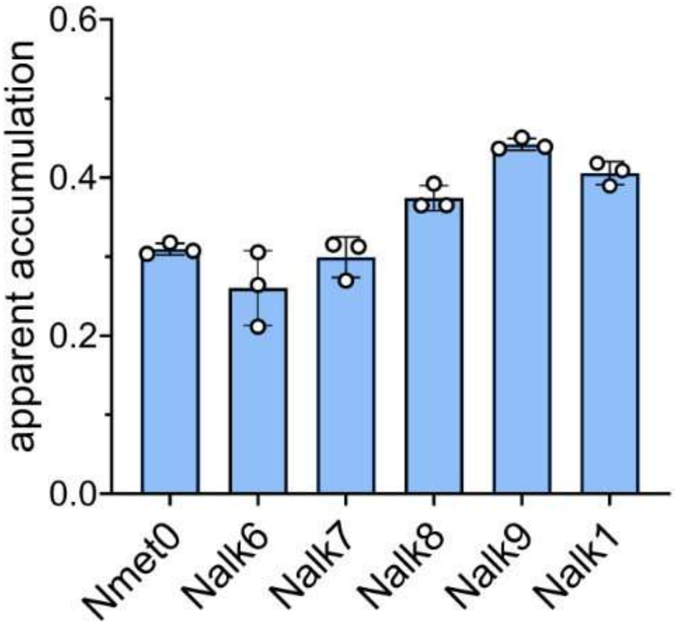


**Figure S12.** Apparent accumulation of the *N*-alkylation library with varying positions of backbone *N*-alkylation. Data are represented as mean ± SD (n= 3).

#### BIOLOGICAL METHODS

##### Reagents

HeLa cells expressing a GFP-HaloTag fusion protein (HaloTag HeLa: HT HeLa) were generously provided by the Chenoweth Lab at the University of Pennsylvania. Dulbecco's Modified Eagle Medium (DMEM), Fetal Bovine Serum (FBS), penicillin-streptomycin, puromycin, and digitonin were obtained from Sigma-Aldrich. A library of 380 azide-containing small molecules was purchased from Enamine (catalog # AZD-380-X-100). Hoechst 33342 and HEPES were acquired from Thermo Fisher Scientific. The plasmid encoding pET-6xHis-GFP (+36) was obtained from Addgene (plasmid #199165). DBCO-NHS ester was purchased from Click Chemistry Tools.

##### Cell Culture

HeLa Halo cells were cultured in Dulbecco's Modified Eagle Medium (DMEM) supplemented with 10% fetal bovine serum (FBS), 1% penicillin-streptomycin, and 1 μg/mL puromycin. Cells were maintained at 37°C with 5% $CO_2$. WT HeLa cells were cultured under identical conditions, except without puromycin selection. HeLa Halo cells were passaged using trypsin upon reaching 80–90% confluency and resuspended in DMEM. Cells were seeded in 96-well plates at a density of 50,000 cells per well, incubated for 24 hours at 37°C with 5% $CO_2$, and subsequently treated with compounds at the required concentration and time point.

##### One-step assay with dye (TMRcl/TMRaz)

HT HeLa cells were seeded into 96-well plates one day before the assay, reaching 80–90% confluency. Following overnight incubation at 37°C, cells were washed three times with PBS and treated with 10 μM DBCOcl for 15 minutes which is followed by chased with 50 μM of a dye (TMRcl/TMRaz) for 15 minutes. Then, the cells were washed three times using PBS, trypsinized for 5 min, and fixed with 4% formaldehyde in PBS for 30 min statically at room temperature. The cells were then subjected to analysis by flow cytometry.

**Chloroalkane HaloTag Azide-based Membrane Penetration (CHAMP) Assay**

HT HeLa cells were seeded into 96-well plates one day before the assay, reaching 80–90% confluency. Following overnight incubation at 37°C, cells were washed three times with PBS and treated with 10 μM DBCOcl for 15 minutes. This was followed by a pulse with azide-tagged compounds at 50 μM concentration, then a chase step with 50 μM TMRaz for 15 minutes. Cells were washed three times with PBS between each step. After staining, cells were washed again, trypsinized, fixed with 4% formaldehyde, and analyzed using flow cytometry on the Attune NxT Acoustic Focusing Cytometer (Invitrogen).

**Confocal Fluorescence Microscopy**

Confocal images were acquired using a Leica SP5X laser scanning microscope and a Leica STELLARIS 8 confocal/FLIM/tauSTED system equipped with tunable white light lasers and a 37-2 digital temperature controller. Image acquisition was performed using the LAS-AF software, and Fiji (ImageJ) was used for further processing. HeLa Halo and unmodified HeLa cells were trypsinized upon reaching 80–90% confluency, resuspended in DMEM, and seeded at a density of 50,000 cells on a 10-mm plate. After 24 hours of incubation at 37°C with 5% $CO_2$, cells were stained with Hoechst 33342 (Thermo Fisher, H3570) following the manufacturer's instructions. Cells were subsequently washed twice with PBS, rinsed three times with pre-warmed DPBS, and incubated in 1.5 mL DMEM containing 25 mM HEPES (Thermo Fisher, 21-063-029) for imaging.

**Screening of Azide-Modified Compound Library**

The azide-modified compound library was screened at 50 μM for 1 hour using the CHAMP assay as described above. Fluorescence data were corrected for background contributions by subtracting signals from blank controls and normalized to the maximum fluorescence signal.

**Membrane Permeabilization Assay**

Digitonin (Sigma, Part No. D141-100MG) was used for the membrane permeabilization assay. For this assay, 40 μg/mL digitonin was co-incubated with peptides for 24 hours following a 15-minute pulse with 10 μM DBCOcl. This was followed by a chase step with

50 μM TMRaz for 15 minutes. Cells were then trypsinized, fixed with 4% formaldehyde, and analyzed using flow cytometry.

**+36GFP amino acid sequence**

MGHHHHHHGGASKGERLFRGKVPILVELKGDVNGHKFSVRGKGKGDATRGKLTLKFI
CTTGKLPVPWPTLVTTLTYGVQCFSRYPKHMKRHDFFKSAMPKGYVQERTISFKKDGK
YKTRAEVKFEGRTLVNRIKLKGRDFKEKGNILGHKLRYNFNSHKVYITADKRKNGIKAKF
KIRHNVKDGSVQLADHYQQNTPIGRGPVLLPRNHYLSTRSKLSKDPKEKRDHMVLLEF
VTAAGIKHGRDERYK

**Expression and Purification of +36GFP and Wild-Type GFP**

Plasmids encoding pET-6xHis-GFP (+36) were transformed into *Escherichia coli* BL21 cells and inoculated into 1 L of Lysogeny Broth (LB) supplemented with 100 μg/mL ampicillin. Cultures were grown at 37°C with shaking at 250 rpm until reaching an OD600 of 0.5. Protein expression was induced by the addition of 1 mM IPTG, followed by incubation at 37°C for 3 hours and then overnight at 24°C. Cells were harvested by centrifugation at 4000 × g for 15 minutes at 4°C. All subsequent purification steps were performed on ice. Cell pellets were lysed by sonication (Fisher Scientific FB505) for 25 minutes in cycles of 3 minutes on and 5 minutes off. Cell debris was removed by centrifugation at 4000 × g for 20 minutes at 4°C, followed by an additional centrifugation at 23,000 × g for 20 minutes to separate soluble and insoluble fractions. Supercharged proteins were purified under native conditions using Ni-NTA agarose chromatography. The wash buffer contained 50 mM phosphate buffer, 1 M NaCl, and 35 mM imidazole (pH 8.0), while the elution buffer contained 50 mM phosphate buffer, 1 M NaCl, and 250 mM imidazole (pH 8.0). Purified proteins were concentrated using an Ultra-15 centrifugal filter unit (Millipore, UFC901024) and dialyzed in 50 mM phosphate-buffered saline (pH 8.4). The azide tag was introduced using azidoacetic NHS ester (Click Chemistry Tools, Az103-100).

#### SYNTHESIS AND CHARACTERIZATION OF PEPTIDES, SMALL MOLECULES, AND ANTIBIOTICS

##### Synthesis of DBCOcl

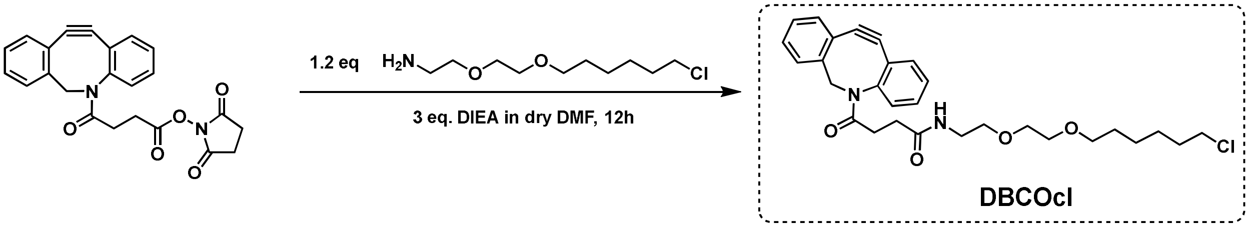


Following a published protocol, 41 mg of DBCO-NHS (0.1 mmol, BroadPharm) in dry DMF was mixed with 32 mg of 2-(2-((6-Chlorohexyl)oxy)ethoxy)ethanamine hydrochloride (0.12 mmol, AmBeed) into which was added 43 µL of DIPEA (0.24mmol).[1] The reaction mixture was stirred at room temperature for 1hr following which, DMF was removed by rotary evaporation of a DMF/toluene mixture. The crude residue was acidified and purified by RP-HPLC (Mobile phase A: 90% v/v water initial to 0% v/v water with 0.1% TFA; mobile phase B: 10% v/v MeCN -100% v/v MeCN with 0.1% over a 70min gradient) to afford 35mg of an oily product (67% yield) which was found to be DBCOcl, the analytical data was in agreement with assigned structure. Calc.: $C_{29}H_{36}ClN_2O_4$ $[M+H]^+$ 511.2358; Obs. 511.2.

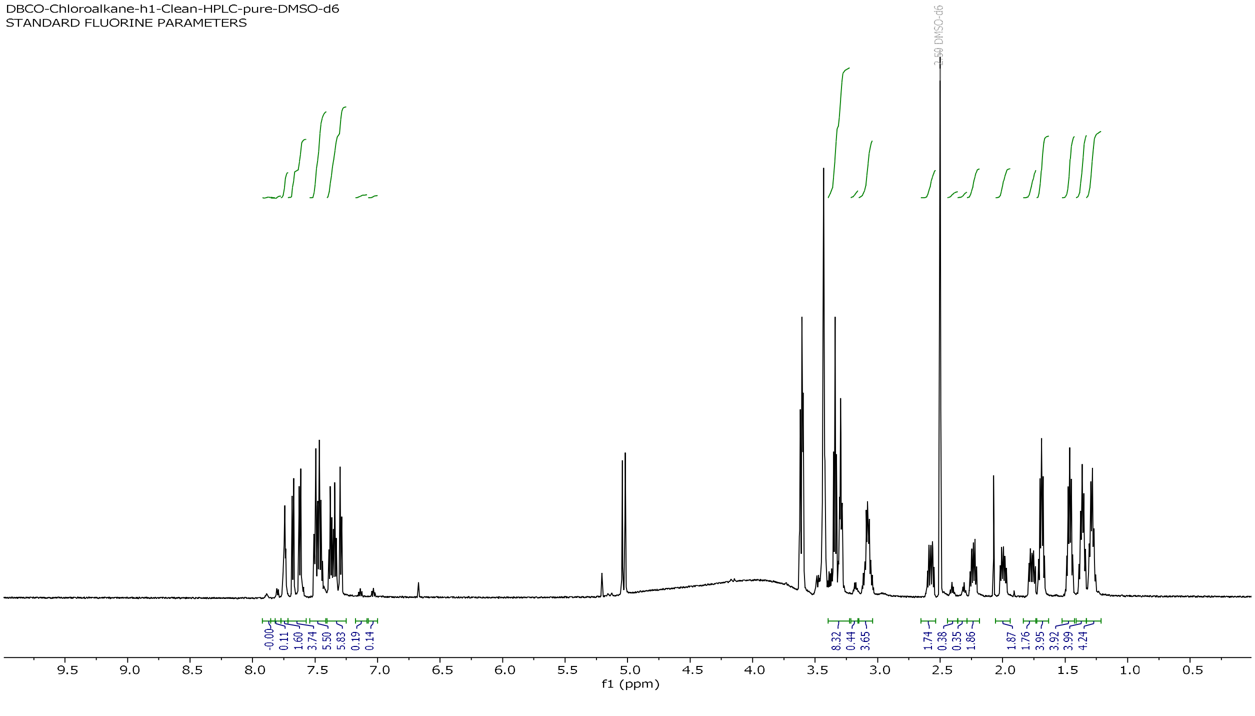

[1]NMR spectrum of DBCOcl.

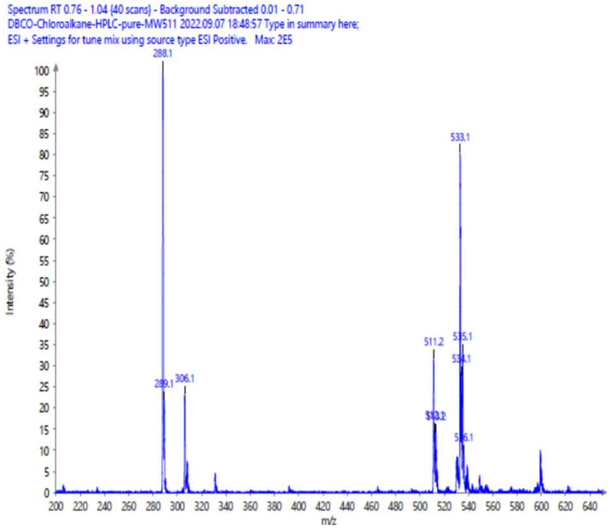

ESI-MS of DBCOcl.

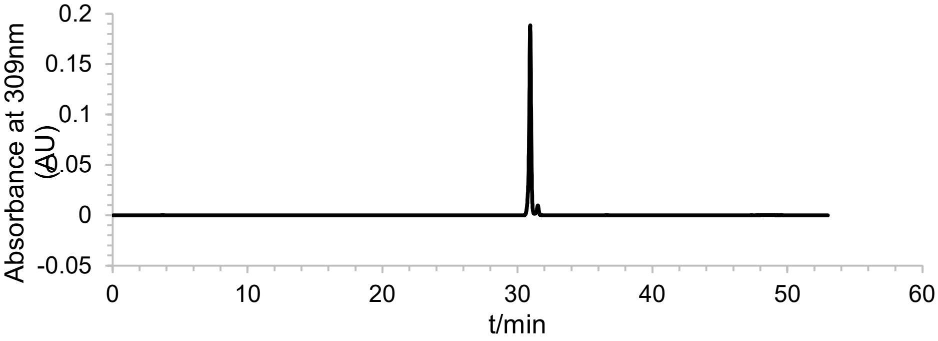


DBCOcl purity check analytical RP-HPLC detection at 309 nm.

#### Synthesis of Polyarginine Peptides

Polyarginine peptides were synthesized using an automated peptide synthesizer via standard solid-phase chemistry. To a 15 mL peptide synthesis vessel, an appropriate amount of resin was added. Rink amide resin (0.48 mmol/g loading capacity) was used to ensure C-terminal amidation upon cleavage. It was followed by addition of 20% piperidine in *N, N*-Dimethylformamide (DMF, 15 mL), and then washed (3X) with *N, N*-Dimethylformamide. After the last wash, 4 equiv. of amino acid was added along with 4 equiv. of ethyl cyanohydroxyiminoacetate (Oxyma) and 4 equiv. of *N, N'*-Diisopropylcarbodiimide (DIC). The remainder of the amino acids were coupled in the same manner. After coupling with the last amino acid, the azide coupling steo was done outside the automated peptide synthesizer. Peptides were cleaved from the resin using a TFA/TIPS/$H_2O$ mixture (95:2.5:2.5, v/v/v) (unless otherwise mentioned) shaking at room temperature for 2 h. The solution was filtered and precipitated by the addition of cold diethyl ether to yield crude peptide. Peptides were purified via reverse-phase high-performance liquid chromatography (RP-HPLC) equipped with Waters 1525 with 2489 UV/Visible Detector on a Phenomenex Luna 10 µm C8(2) 100 Å (250 x 21.2 mm) column using gradient elution with $H_2O$/MeOH with 0.1% TFA at 10 mL/min. The HPLC fractions of the desired purified compounds were first concentrated under reduced pressure using a rotary evaporator. The final concentrated aqueous solutions were lyophilized to dryness using Labconco Freezone 4.5 L (-84 $^oC$) lyophilizer. and verified to be >97% pure by RP-HPLC and MALDI-TOF mass spectrometry. Peptide identities were confirmed via high resolution electrospray ionization mass spectrometry (HRMS, ESI/MS). Analyses were obtained on an Agilent 6545B Q-TOF LC/MS equipped with 1260 infinity II LC system with auto sampler. Samples were dissolved in $CH_3CN$ and eluted with a $CH_3CN/H_2O$ solution containing 0.1% formic acid.

#### Characterization of Test Molecules

Peptides were analyzed for purity using a Phenomenex Luna 5 µm C8(2) column via RP-HPLC with gradient elution in $H_2O$/MeOH containing 0.1% trifluoroacetic acid (TFA) at a flow rate of 1 mL/min. Peptide identities were confirmed via high-resolution electrospray ionization mass spectrometry (HRMS, ESI/MS) using an Agilent 6545B Q-TOF LC/MS system equipped with a 1260 Infinity II LC system. Samples were dissolved in acetonitrile and eluted with a $CH_3CN/H_2O$ solution containing 0.1% formic acid. Characterization data are provided in supplementary information.

**Analytical HPLC Chromatogram of R5az**

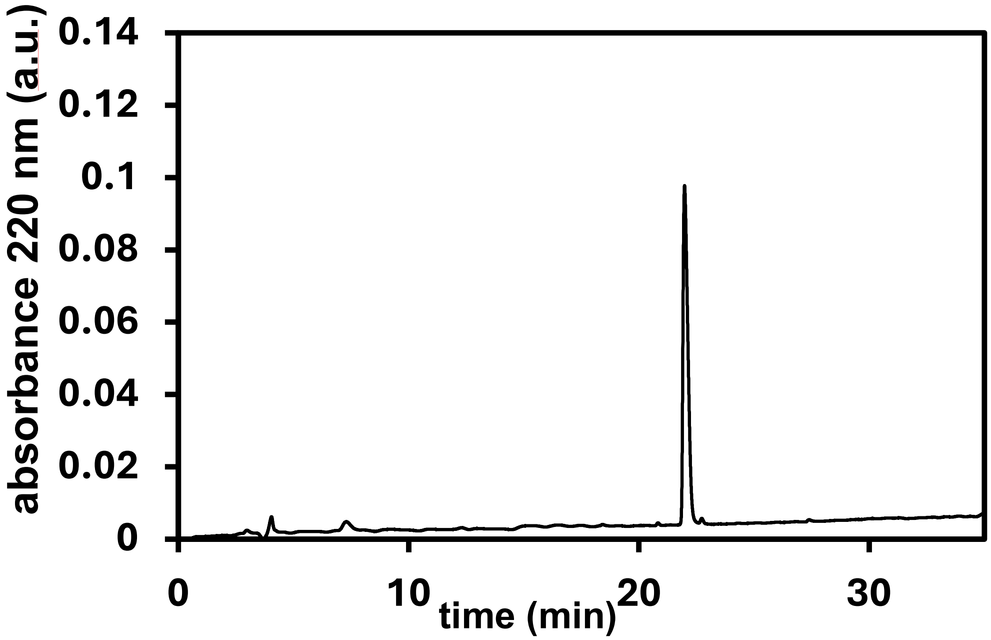

QTOF High Resolution Mass Spectrum for R5az (m/z 880.5441for $[M+H]^+$ found 881.5522)

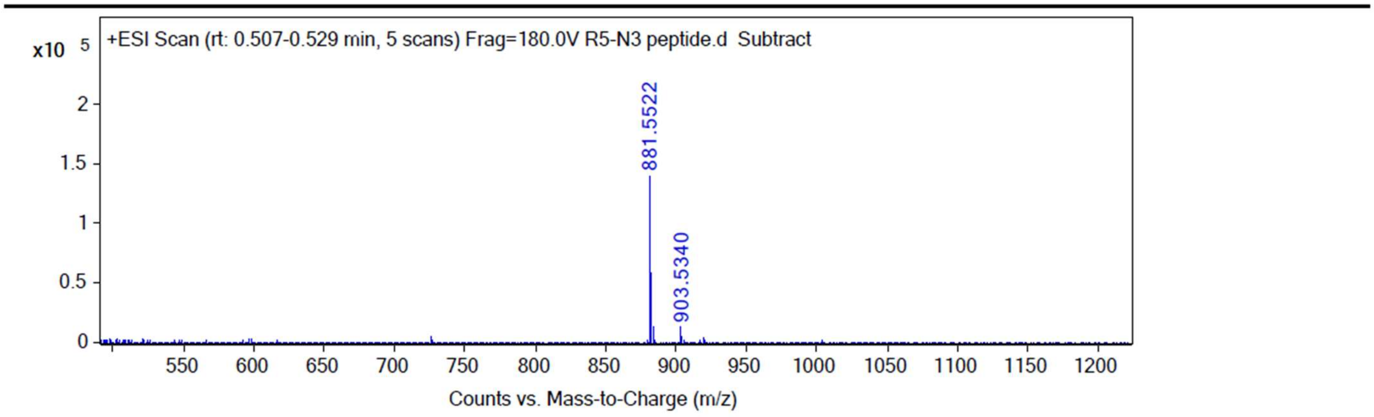

Analytical HPLC Chromatogram of R7az

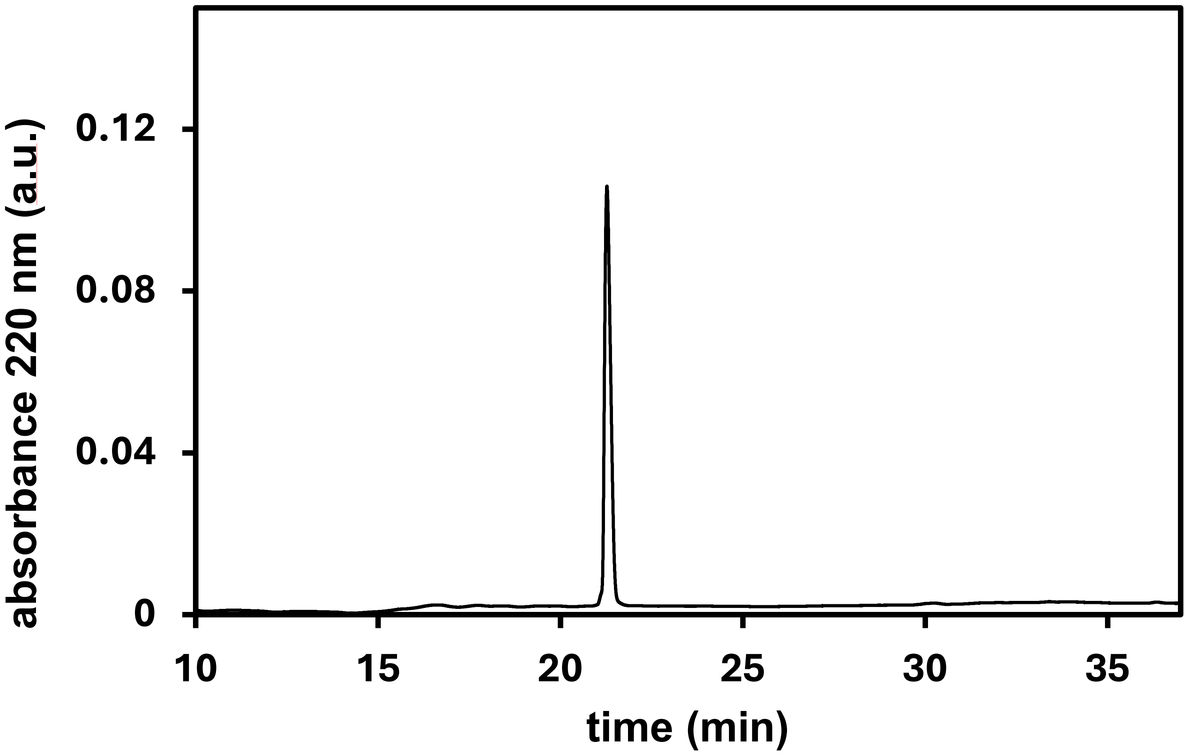

QTOF High Resolution Mass Spectrum for R7az (m/z 1192.7463) for $[M+H]^+$ found 1193.7542)

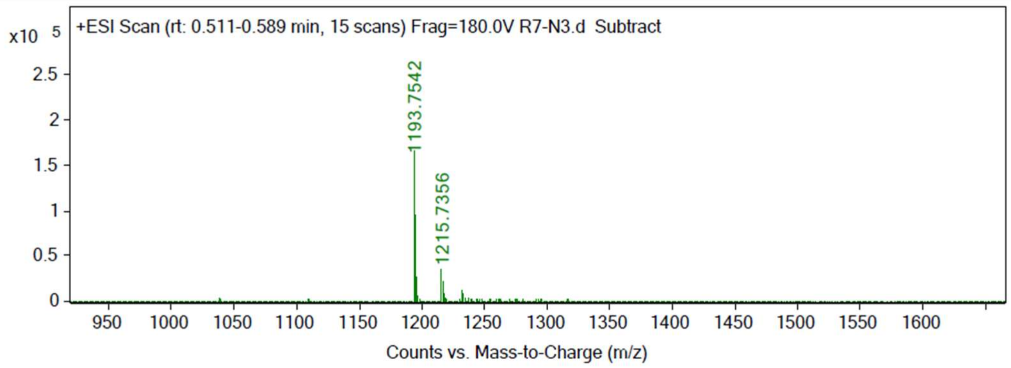

Analytical HPLC Chromatogram of R9az

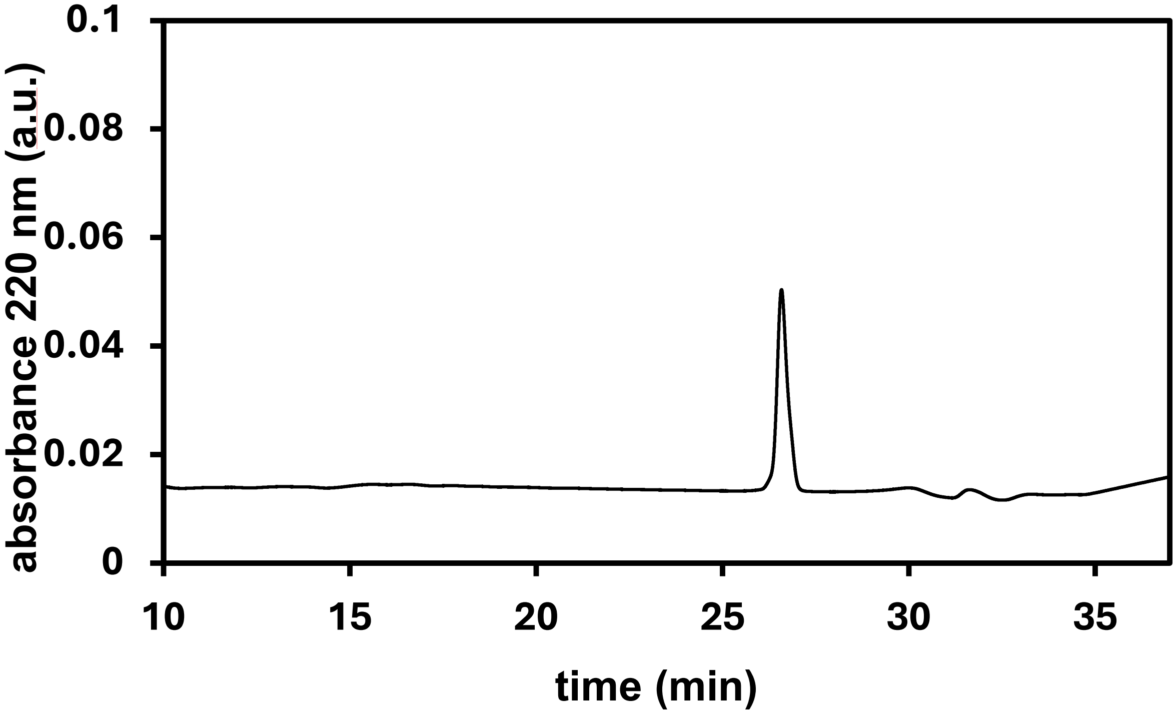

QTOF High Resolution Mass Spectrum for R9az (m/z .1504.9485 for $[M+H]^+$ found 1505.9566)

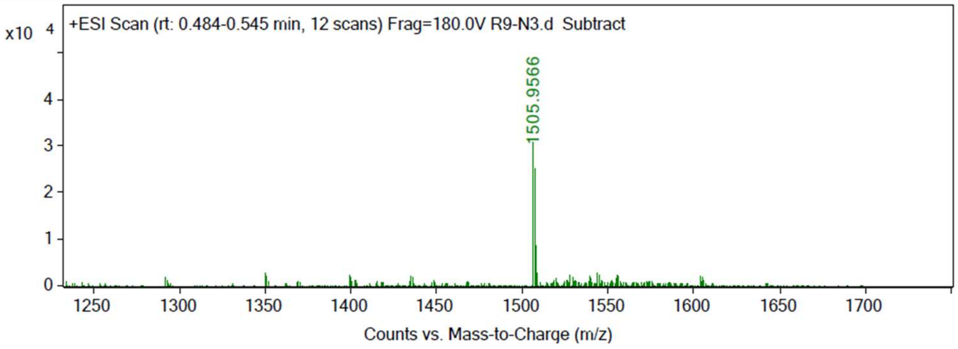

Analytical HPLC Chromatogram of R11az

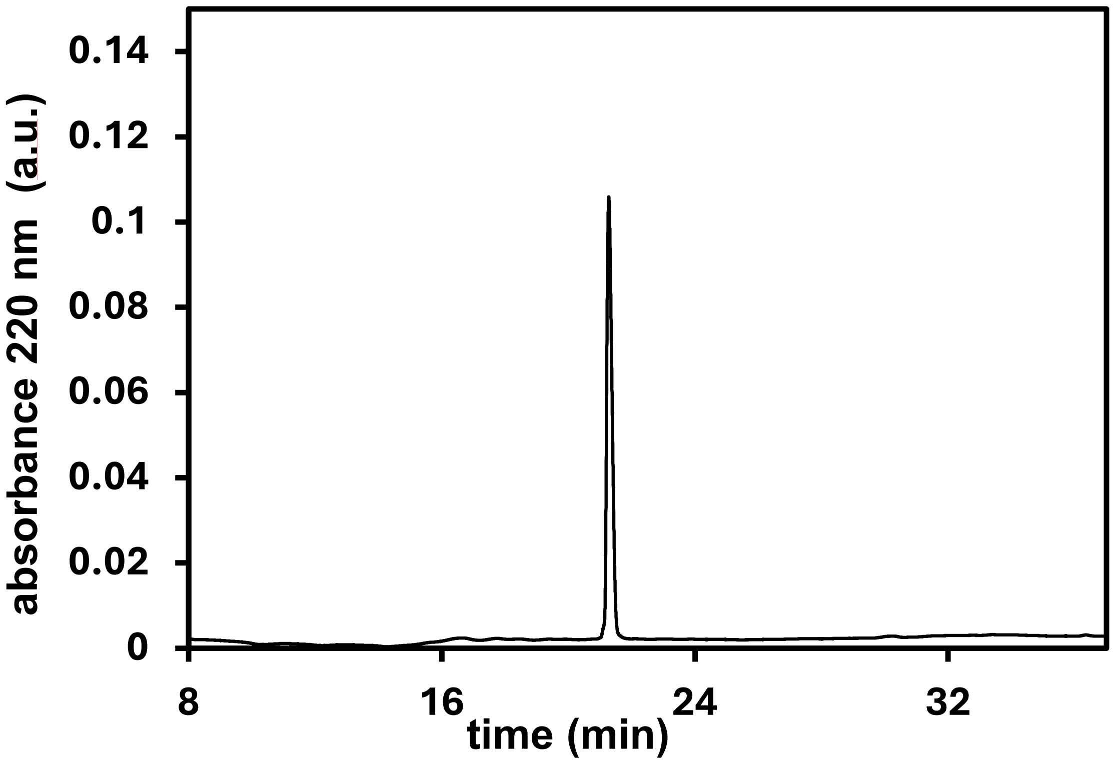

QTOF High Resolution Mass Spectrum for R11az (m/z 1817.1501 for $[M+H]^+$ found 188 .5522)

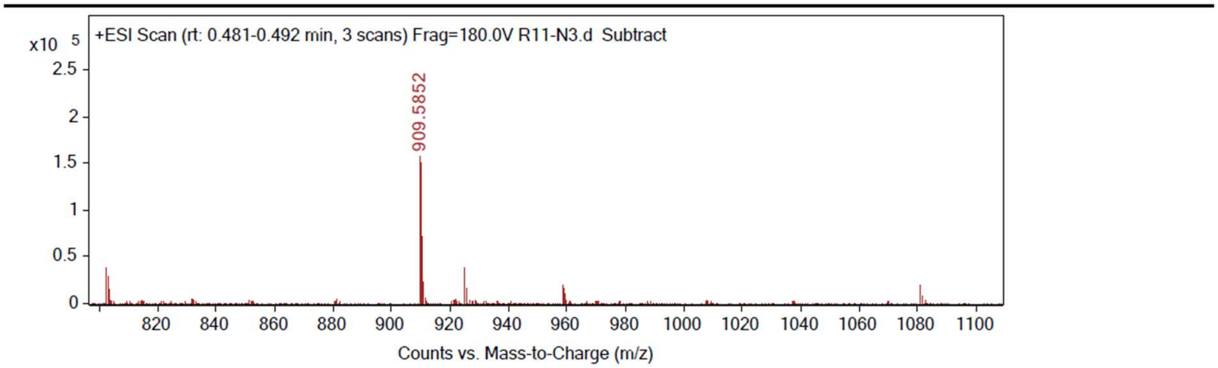

**Synthesis of the macrocyclization peptide libraries**

To prepare peptides in this library, Rink-amide resin (0.48 mmol/g of loading capacity) was used such that the *C*-terminus would be amidated upon cleavage. For all linear analogs within the library, the *N*-terminus was acetylated by combining the peptide with a mixture of acetic anhydride, DIEA, and DMF (5:8.5:86.5, v/v) for 1 hour at room temperature. For the cyclization chemistry of the **Cyc0-4** and **Lar1-5** series, 10 equiv. of chloroacetic chloride and 15 equiv. of DIEA were dissolved in anhydrous DCM and coupled to the *N*-terminally deprotected peptide by shaking it for 2 hours at room temperature. Cleavage from resin was performed as described above. For the linear analogs, purification was then subsequently carried out as described above. For the macrocyclic peptides **Cyc0-4** and **Lar1-5**, the cleaved, crude peptide scaffolds were then dissolved in a mixture of $H_2O/CH_3CN$ (1:1, v/v) to an approximate concentration of ~1.5 mM. This was followed by the addition of 0.5 M $NH_4HCO_3$ (pH 8.5) to a final concentration of 50 mM to form the thioether bonds for macrocyclization. The cyclization reaction mixture was then stirred overnight at room temperature (~16 h). Cyclization was monitored using matrix-assisted laser desorption ionization time-of-flight (MALDI-TOF) mass spectroscopy (Shimadzu 8020). For the macrocyclization of peptide Dit1, the crude peptide scaffold was dissolved in a mixture of $H_2O/CH_3CN$ (1:1, v/v) to an approximate concentration of ~1.5 mM. This was followed by the addition of 0.5 M $NH_4HCO_3$ (pH 8.5) to a final concentration of 50 mM to form the disulfide bond required for macrocyclization. The cyclization reaction mixture was then stirred overnight at room temperature (~16 h). Cyclization was assessed by monitoring the presence of free thiols using Ellman's reagent as per the standard protocol provided by the manufacturer. For the macrocyclization **Dit2**, the crude peptide scaffold was dissolved in a mixture of $H_2O/CH_3CN$ (1:1, v/v) to an approximate concentration of ~1.5 mM. Further, the bis-electrophilic cyclization linker 1,4-Bis(bromomethyl)benzene was also dissolved into this mixture to an approximate concentration of ~1.5 mM (1 equiv.). Thioether bond formation between the **Dit2** peptide scaffold and the linker was brought about by the addition of 0.5 M $NH_4HCO_3$ (pH 8.5) to a final concentration of 50 mM. The cyclization reaction mixture was then stirred overnight at room temperature (~16 h). Cyclization was monitored using matrix-assisted laser desorption ionization time-of-flight (MALDI-TOF) mass spectroscopy (Shimadzu 8020). For all the above-mentioned macrocyclic peptides, upon confirmation of macrocyclization, the samples were then concentrated under reduced pressure using a rotary evaporator. The concentrated aqueous solutions were then lyophilized to dryness using a Labconco Freezone 4.5 L (-84 $^{o}$C) lyophilizer. Purification was subsequently carried out as described above. All peptides in this library were characterized using UV-Vis absorbance of the phenylalanine residue in their sequence at 257 nm (ε = 195 $cm^{-1}M^{-1}$). For **Dit2**, the extinction coefficient at 257 nm of 1,4-Bis(bromomethyl)benzene was also determined separately using a calibration curve and accounted for during concentration determination of **Dit2**.

##### Synthesis of the *N*-methylation and *N*-alkylation peptide libraries

To prepare peptides in this library, Rink-amide resin (0.48 mmol/g of loading capacity) was used such that the *C*-terminus would be amidated upon cleavage. Commercially available *N*-methyl amino acids were used for the *N*-methyl analogs in the peptide library. 2-azido-acetic acid was coupled on the *N*-terminus of each peptide on resin. For the peptoid library, the *N*-alkyl substituted glycine residues in the peptoids were added as per reported protocols. Briefly, a 25 mL peptide synthesis vessel containing 100 mg of Rink amide resin (0.048 mmol), conjugated with or without peptide residues, was added 20% piperidine in DMF (15 mL), with shaking at room temperature for 30 min. The resin was then washed with $CH_3OH$/DCM. After the last wash, to the resin was added 1 mL 0.6 M 2-bromo acetic acid with 200 µL of 50% DIC in DMF. The resin was shaken at room temperature for 20 min, then washed with DMF 5 times. 1 mL 1.5 M primary amine building blocks were then added to the resin and the resin was shaken at room temperature for 1-2 h. The rest of the amino acids are coupled in the same manner according to the sequence of the peptide. As before, 2-azido-acetic acid was coupled on the *N*-terminus of each peptide/peptoids on resin. Cleavage and purification were carried out as described before. All peptides in this library were characterized using UV-Vis absorbance of the phenylalanine residue in their sequence at 257 nm (ε = 195 $cm^{-1}M^{-1}$).

#### General methods for synthesis of peptide probes

All peptides were built by solid-phase peptide synthesis. Amidated C-terminus peptides were built starting from Fmoc-aminomethyl phenoxy resin (Rink amide resin) procured from Sigma-Aldrich (St. Louis, MO, USA); COOH terminated peptides were built on either chloro-trityl resin (Sigma-Aldrich) or Fmoc-amino acid preloaded on Wang resin procured from p2bioSystems (Louisville, KY 40223). Crude peptides were purified by preparative reverse phase HPLC purification instruments equipped with Waters 1525 pumps and 2489 UV/Visible Detector on a Phenomenex Luna 10 μm C8(2) 100 Å (250 x 21.2 mm) or C18 columns using a 5 to 100% linear gradient of methanol (MeOH) in $H_2O$ or acetonitrile (MeCN) in water each containing 0.1% TFA at 10 mL/min. The HPLC fractions of the desired compounds were first concentrated under reduced pressure using a rotary evaporator. The concentrated aqueous solutions were then frozen and lyophilized with Labconco Freezone 4.5L lyophilizer (-84$^{o}$C). The purity of the samples was ascertained by analytical HPLC using a Phenomenex Luna 5 μm C8(2) or C18 columns of 100 Å (250 x 4.6 mm) on the same instrument; using gradient elution in $H_2O$/MeCN or $H_2O$ /MeOH with 0.01% TFA in each solvent at 1 mL/min prior to use in biological assays. Electrospray ionization (ESI)-based mass analysis was performed on Advion Expression® CMS mass spectrometer using standard parameters for intermediates. A low fragmentation and low energy setup were used to analyze fragmentation-sensitive compounds. Reaction progress and purification of products were analyzed with Additional LCMS analysis, which were performed on Shimadzu Prominence-i LCMS system equipped with LC-2030C 3D liquid chromatograph, autosampler, PDA and LCMS-2020 mass detector. Data analysis was performed with Lab Solutions software. MALDI-TOF mass spectra were obtained for certain high molecular weight compounds on the Shimadzu MALDI-8020 instrument with α-Cyano-4-hydroxycinnamic acid (α-CHCA) matrix. The observed molecular weights for compounds were represented as m/z. High resolution electrospray ionization mass spectrometry (HRMS, ESI/MS) analyses were obtained on an Agilent 6545B Q-TOF LC/MS equipped with 1260 Infinity II LC system with auto sampler and equipped with C18(2) column (Luna 5 μm 100Å 250 x 4.6 mm. Quantitative LCMS analysis was performed on Agilent 6545B Q-TOF as describe above in positive ion mode. For each antibiotics-azide tested, parameters were such as collision energy and fragmentation pattern (formation of defined daughter ions and their intensities were optimized). Prior to actual quantification of recovered antibiotics-azides from cell lysates, standard curve at lowest possible concentration were obtained. The LCMS/MS data was processed with Mass Hunter software, and subsequent quantification was done with Q-TOF MS quantitative software. UV spectroscopic analysis was performed on Genesys 50 (Thermo Scientific) UV-Visible Spectrophotometer. $^{1}$H and $^{13}$C-NMR spectra for final compounds and intermediates were acquired on a Varian 600MHz spectrophotometer. $^{1}$H and $^{13}$C-NMR spectra for final compounds and intermediates were acquired on a Varian 600MHz spectrophotometer. All NMR spectra were processed and analyzed using MestreNova software. Residual solvent signals from deuterated solvents were used as an internal standard with reference to tetramethylsilane (TMS) for defining chemical shifts. Chemical shifts are reported in ppm (d) and coupling constants (J) are reported in Hertz [Hz].

Fmoc-Lys(N3)-OH (**2n**) is commercially available and was purchased from aapptec (Louisville, KY 40228) and used without further purification.

**Synthesis of 1p.**

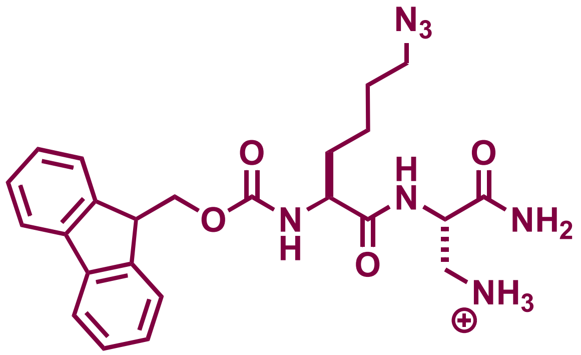


100 mg of Fmoc-aminomethyl)-phenoxy resin (Rink amide resin) of loading capacity 0.48 mmol/g was swollen in DCM after which the Fmoc protecting group was removed using 20% piperidine in DMF for 30 min. The resin was washed using alternating DCM and MeOH and finally rinsed with DMF. Fmoc-Dap (Boc)-OH was coupled to the resin using 4 equivalents of the amino acid and 4 equivalents each of Oxyma and DIC; the coupling reaction was done at room temperature with shaking for 2 h. The Fmoc group was similarly removed using 20% piperidine and following wash steps as described above, Fmoc-Lys($N_3$)-OH was similarly coupled to Dap (Boc) to make Fmoc-Lys($N_3$)-Dap(Boc)-Rink amide resin. The peptide was cleaved off the resin using 95% TFA:2.5% TIPS: 2.5% $H_2O$ and purified by preparative RP-HPLC on a on a Phenomenex Luna 10 µm C8(2) 100 Å (250 x 21.2 mm) using a 20 to 100% linear gradient of MeCN in $H_2O$ each containing 0.1% TFA at 10 mL/min. ESI-MS $[M+H]^+$ m/z calcd. 480.2354 for $C_{24}H_{30}N_7O_4$, obsd. 480.2356. $^1H$ NMR (600 MHz, CD3CN) δ 1.41-1.44 (m,1H), 1.55-1.59 9M, 2H), 1.60-1.67 (m, 4H), 1.78-1.82 (m, 1H), 3.23 (dd, J= 1 and 6 Hz, 1H), 3.29 (t, J= 6Hz, 2H), 3.38 (dd, J= 1 and 6 Hz, 1H), 4.05 (brs, 1H), 4.23 (t, J= 6Hz, 1H), 4.31 (t, J= 6Hz, 1H), 4.35 (dd, J= 2 and 6Hz, 1H), 4.64 (brs, 1H), 6.14 (brs, 1H), 6.57 (brd, J= 6Hz, 1H), 7.07 (brs, 1H), 7.34 (t, J= 6Hz, 2H), 7.42 (t, J= 6Hz, 2H), 7.69 (brt, J= 6Hz, 2H), 7.83 (d, J= 6Hz, 2H), 8.20 (brd, J= 6Hz, 1H).

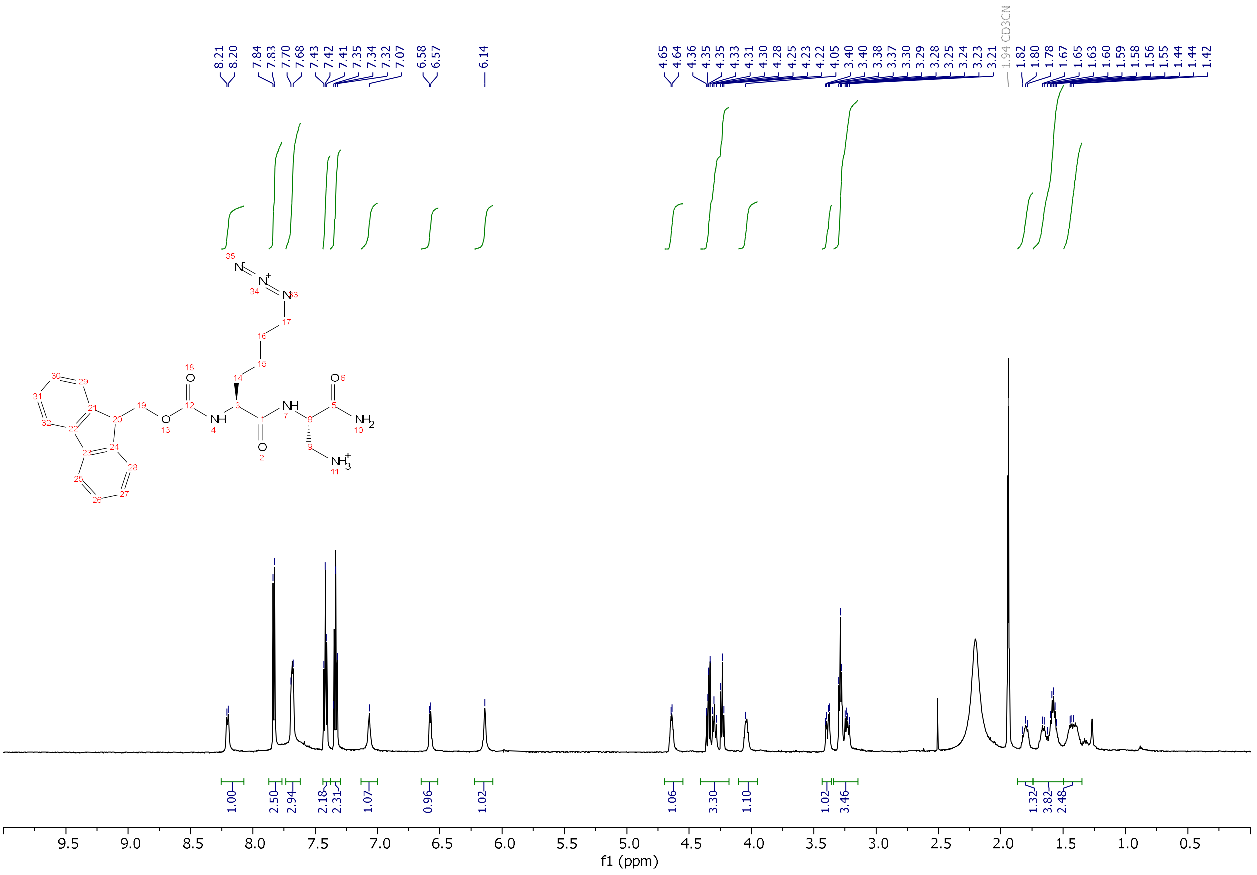

$^1$H-NMR of **1p** in $CD_3CN$.

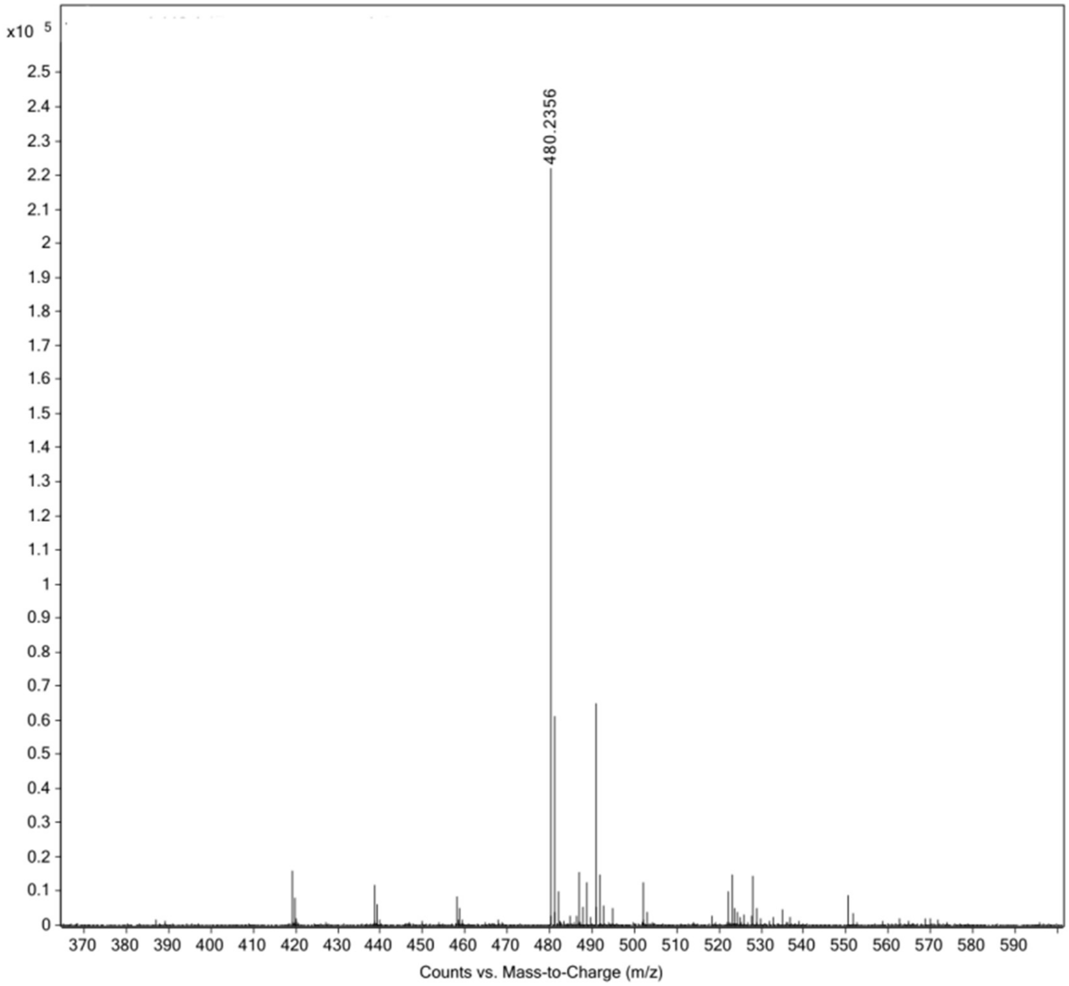


High resolution mass spectrum (ESI) of **1p**.

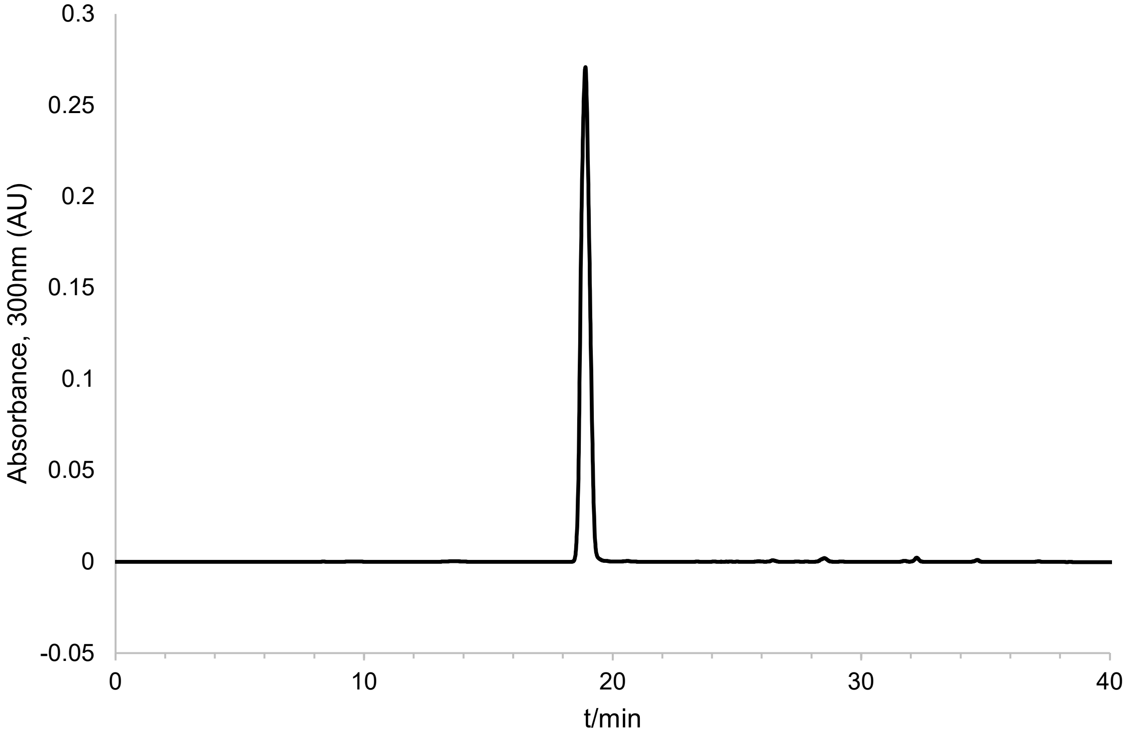


Analytical HPLC chromatogram of **1p**.

**Synthesis of 1n.**

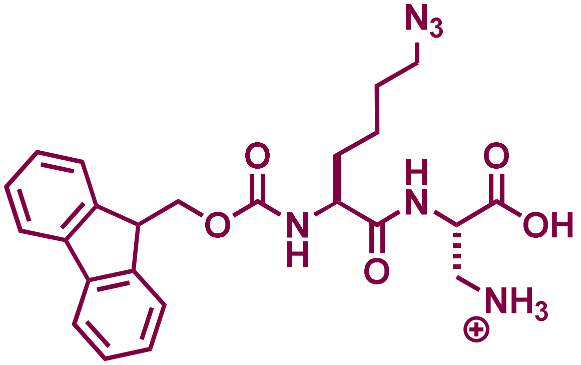


50 mg of 2-chlorotrityl chloride resin (Rink amide resin) of loading capacity 1.5 mmol/g was swollen in dry DCM in a peptide synthesis vessel. Fmoc-Dap (Boc)-OH was dissolved in dry DCM (1eq.) and added to the resin and shaken for 5min after which 4 eq of DIEA was added and incubated for 1 h with shaking at room temperature. The resin was washed 3X with dry DCM, washed 1X with MeOH, and rinsed with DMF. Next, the Fmoc protecting group was removed using 20% piperidine in DMF for 30min. The resin was washed using alternating DCM and MeOH and finally rinsed with DMF. Fmoc-Lys($N_3$)-OH was coupled to the resin using 4 equivalents of the amino acid and 4 equivalents each of Oxyma and DIC; the coupling reaction was done at room temperature with shaking for 2h to make Fmoc-Lys($N_3$)-Dap (Boc)-trityl chloride resin. The peptide was cleaved off the resin using 95% TFA:2.5% TIPS: 2.5% $H_2O$and purified by preparative RP-HPLC on a on a Phenomenex Luna 10 µm C8(2) 100 Å (250 x 21.2 mm) using a 20 to 100% linear gradient of MeCN in $H_2O$ each containing 0.1% TFA at 10 mL/min. ESI-MS $[M+H]^+$ m/z calcd. 481.2194 for $C_{24}H_{29}N_6O_5$, obsd. 481.2197. $^1H$ NMR (600 MHz, CD3CN) δ 1.41-1.44 (m,1H), 1.55-1.59 9M, 2H), 1.60-1.67 (m, 4H), 1.78-1.82 (m, 1H), 3.23 (dd, J= 1 and 6 Hz, 1H), 3.29 (t, J= 6Hz, 2H), 3.38 (dd, J= 1 and 6 Hz, 1H), 4.05 (brs, 1H), 4.23 (t, J= 6Hz, 1H), 4.31 (t, J= 6Hz, 1H), 4.35 (dd, J= 2 and 6Hz, 1H), 4.64 (brs, 1H), 6.14 (brs, 1H), 6.57 (brd, J= 6Hz, 1H), 7.07 (brs, 1H), 7.34 (t, J= 6Hz, 2H), 7.42 (t, J= 6Hz, 2H), 7.69 (brt, J= 6Hz, 2H), 7.83 (d, J= 6Hz, 2H), 8.20 (brd, J= 6Hz, 1H).

$^{13}$C NMR (151 MHz, $CD_3CN$) δ 23.5, 29.0, 31.8, 42.1, 47.9, 50.9, 51.9, 56.2, 118.3, 121.0, 126.2, 126.2, 128.1, 128.8, 142.1, 144.9, 145.0, 157.8, 170.5, 174.6, 183.4.

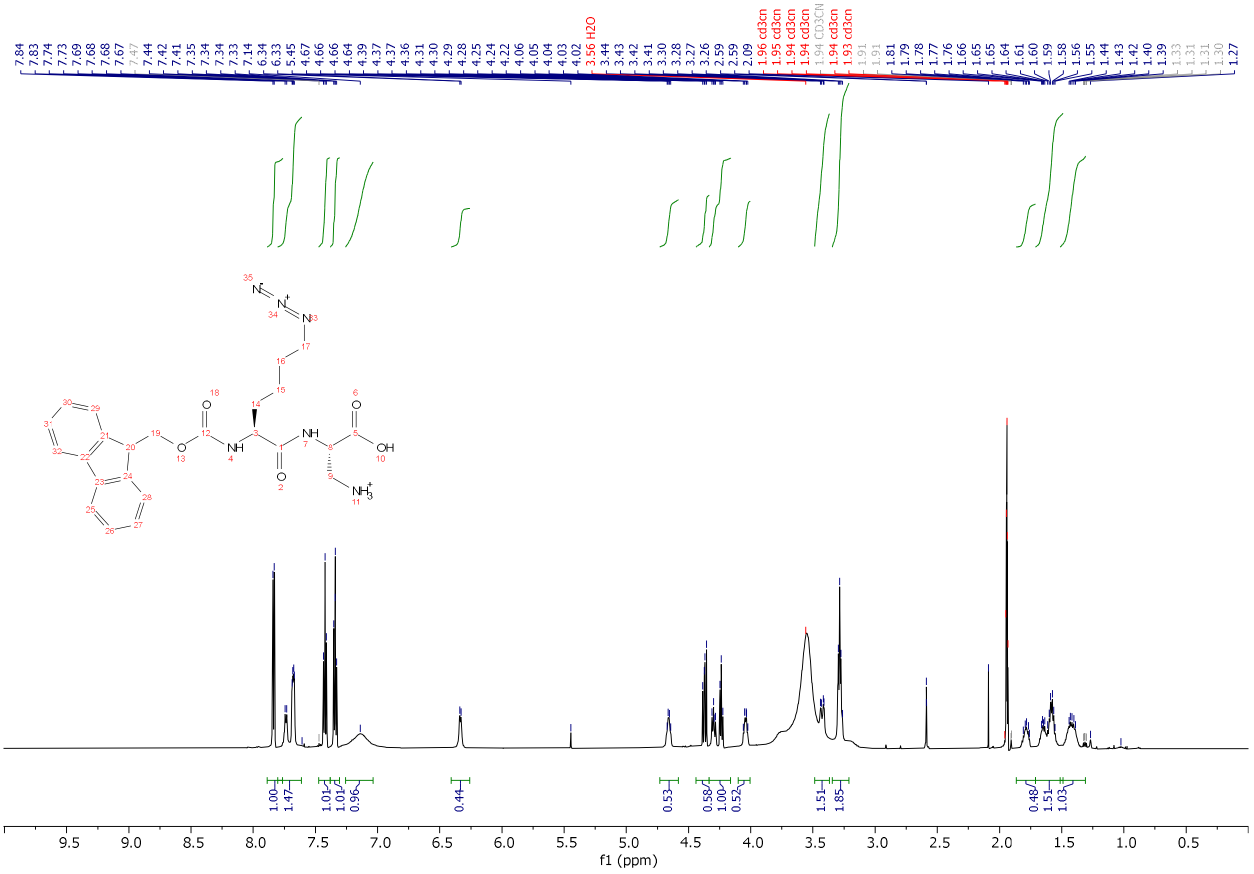

$^1$H-NMR of **1n** in $CD_3CN$.

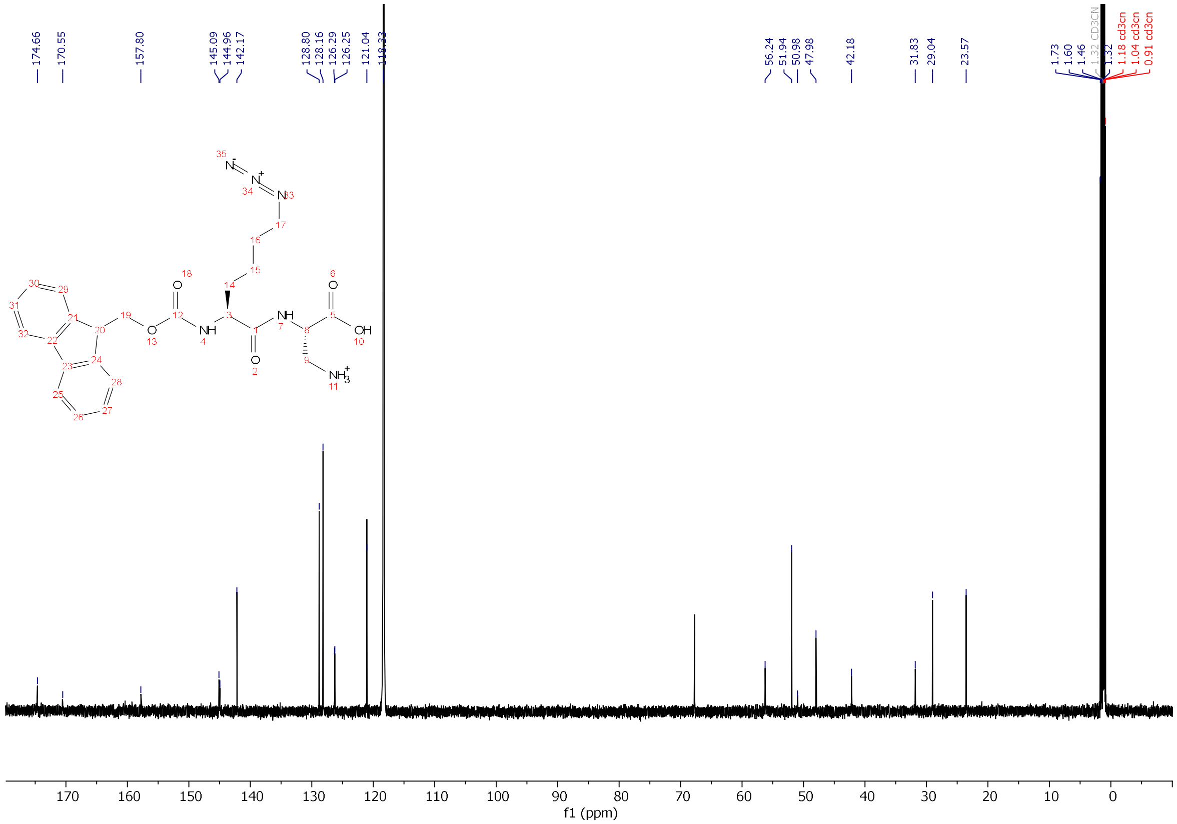

$^{13}$C-NMR of **1n** in $CD_3CN$.

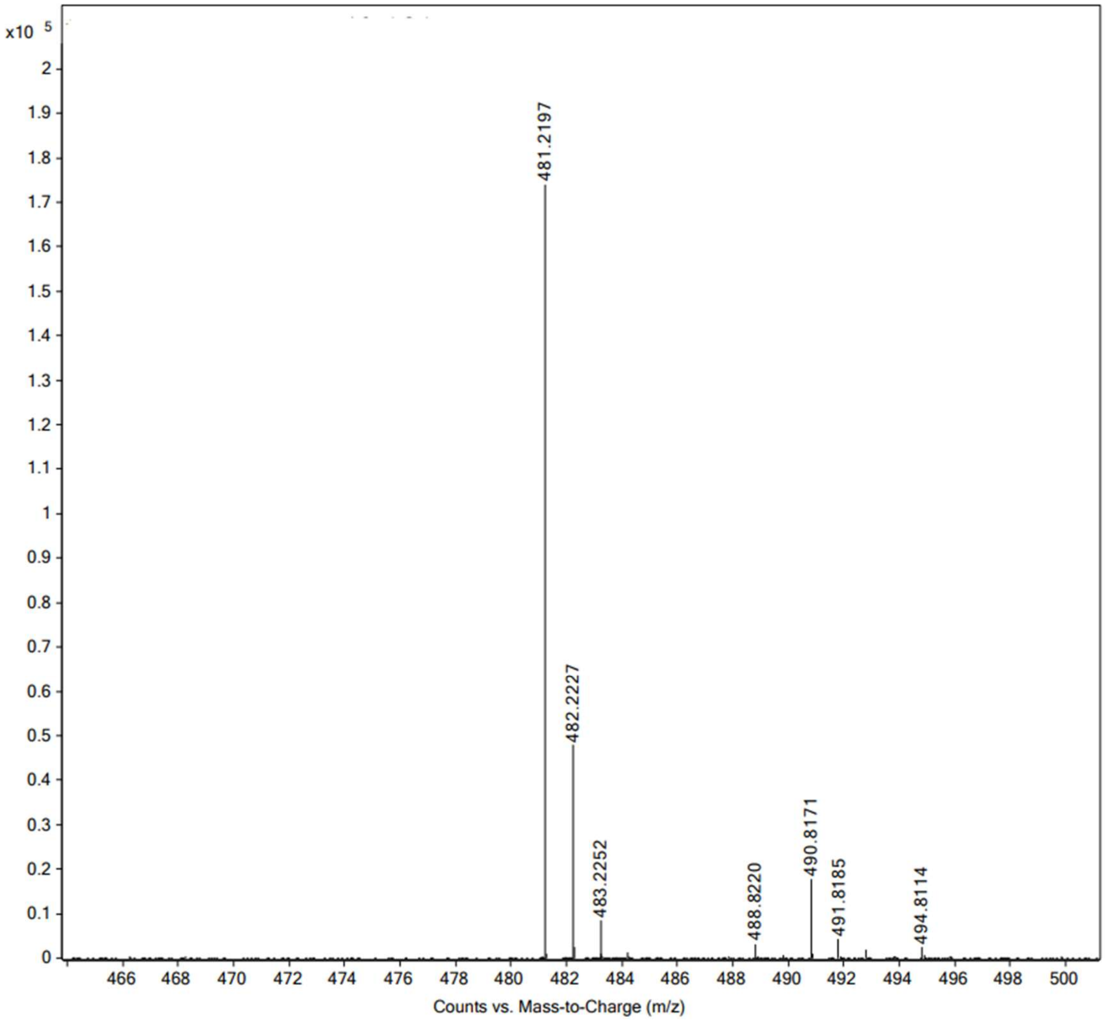


High resolution mass spectrum (ESI) of **1n**.

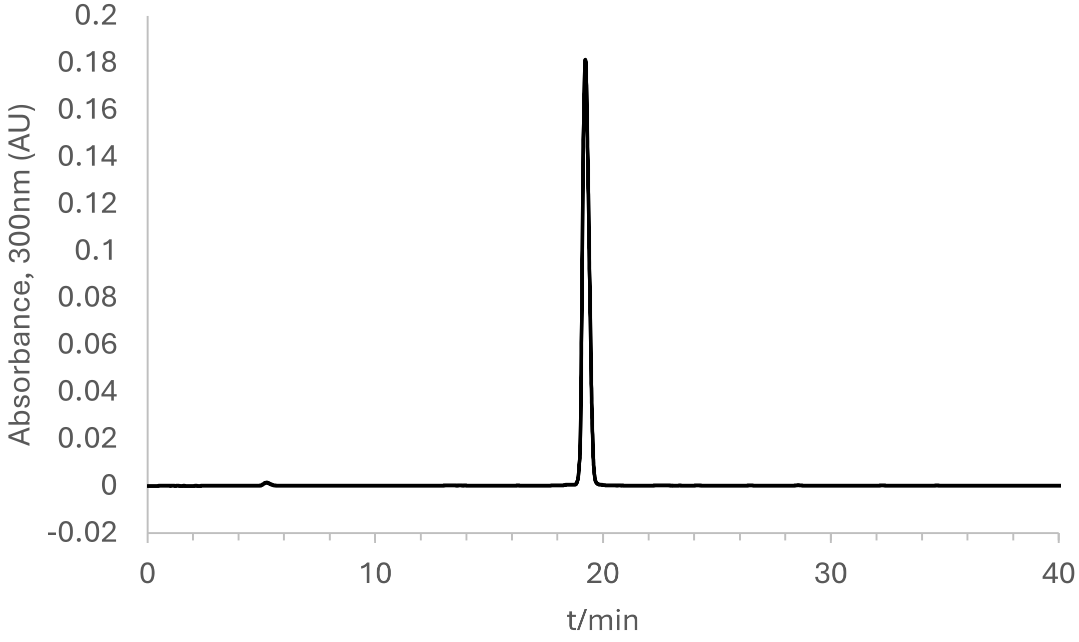


Analytical HPLC chromatogram of **1n**.

**Synthesis of 2p.**

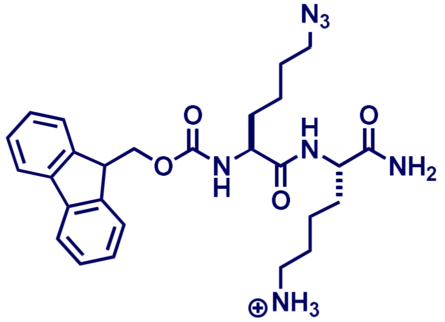


100 mg of Fmoc-(aminomethyl)-phenoxy resin (Rink amide resin) of loading capacity 0.48 mmol/g was swollen in DCM after which the Fmoc protecting group was removed using 20% piperidine in DMF for 30 min. The resin was washed using alternating DCM and MeOH and finally rinsed with DMF. Next, Fmoc-Lys (Boc)-OH was coupled to the resin using 4 equivalents of the amino acid and 4 equivalents each of Oxyma and DIC; the coupling reaction was done at room temperature with shaking for 2h. The Fmoc group was similarly removed using 20% piperidine and following wash steps as described above, Fmoc-Lys($N_3$)-OH was similarly coupled to Lys (Boc) to make Fmoc-Lys($N_3$)-Lys(Boc)-Rink amide resin. The peptide was cleaved off the resin using 95% TFA:2.5% TIPS: 2.5% $H_2O$ and purified by preparative RP-HPLC on a on a Phenomenex Luna 10 µm C8(2) 100 Å (250 x 21.2 mm) using a 20% to 100% linear gradient of MeCN in $H_2O$ each containing 0.1% TFA at 10 mL/min. ESI-MS $[M+H]^+$ m/z calcd. 522.2823 for $C_{27}H_{36}N_7O_4$, obsd. 522.2825 and $[M+Na]^+$ 544.2651. $^1H$ NMR (600 MHz, $CD_3CN$) δ 1.30-1.40 (m, 2H), 1.50-1.65 (m, 3H), 1.72-1.79 (m, 1H), 2.86 (m, 1H), 3.27 (t, J=6 Hz, 1H), 4.01 (dd, J=6, 8 Hz, 1H), 4.21- 4.24 (m, 1H), 4.28 (dd, J= 6Hz, 1H), 4.36 (dd, J= 6Hz, 1H), 7.33 (t, J= 6Hz, 2H), 7.41 (t, J= 6Hz, 2H), 7.66 (brt, J= 6Hz, 2H), 7.82 (d, J= 6Hz, 2H).

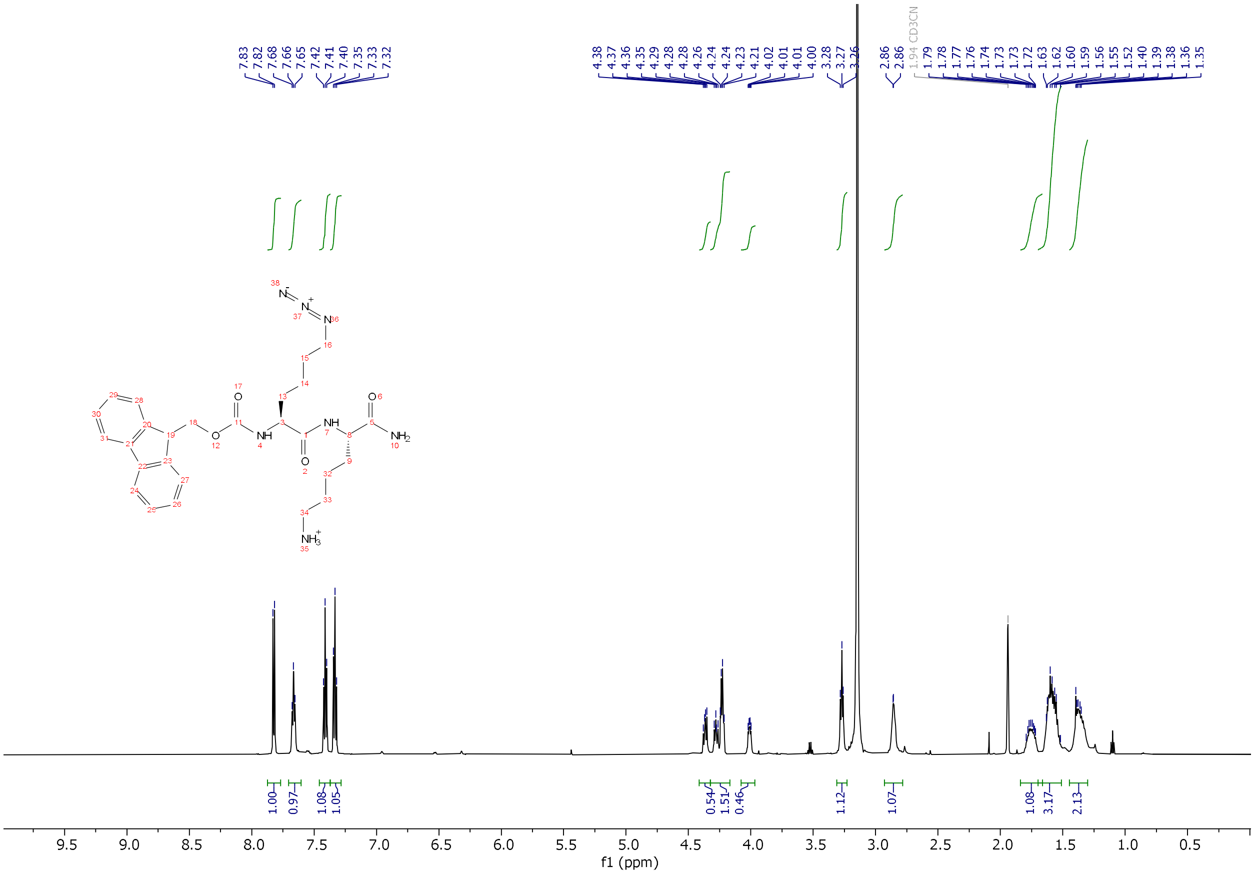

[1]H-NMR of **2p** in $CD_3CN$.

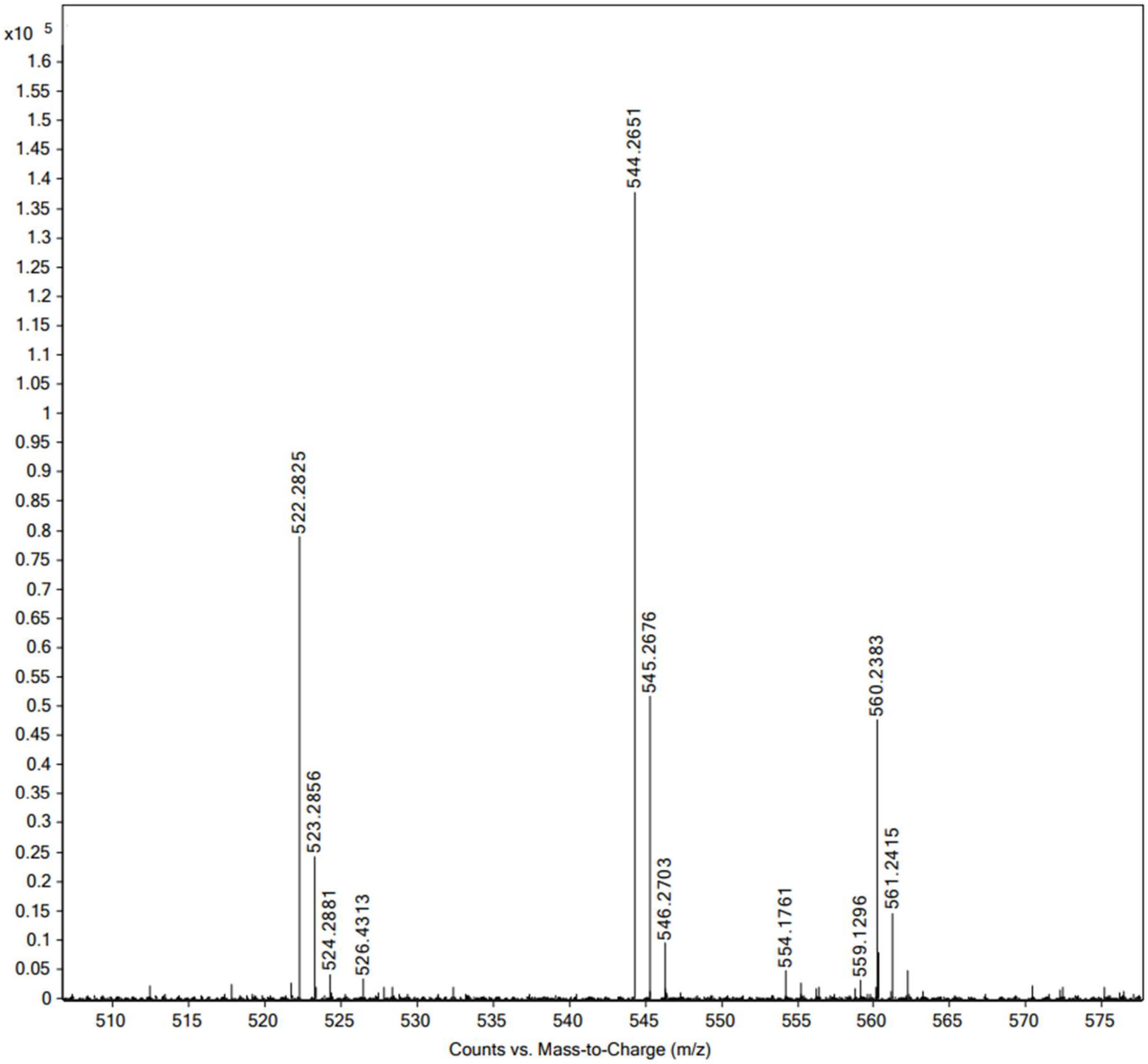


High resolution mass spectrum (ESI) of **2p**.

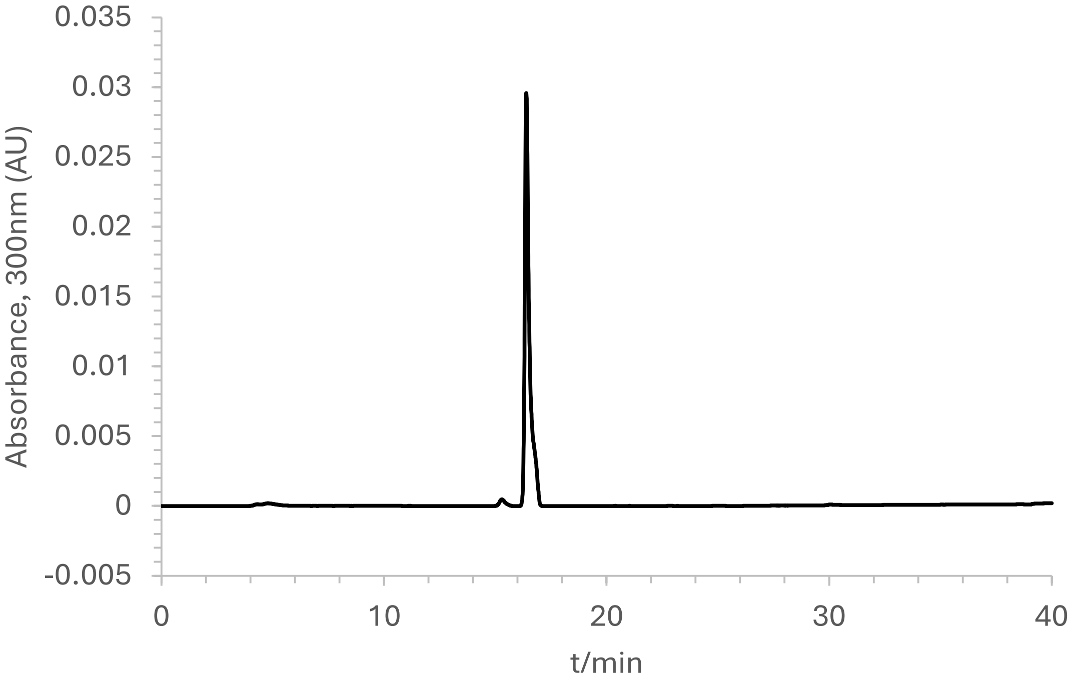


Analytical HPLC chromatogram of **2p**.

**Synthesis of 2n**.

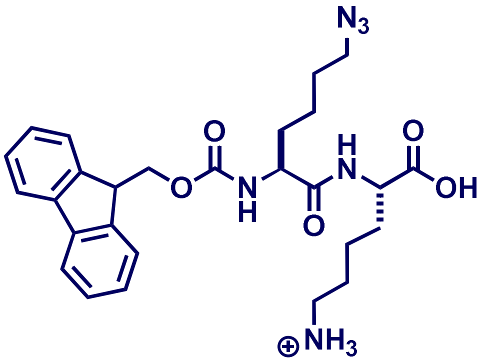


50 mg of 2-chlorotrityl chloride resin (Rink amide resin) of loading capacity 1.5mmol/g was swollen in dry DCM in a peptide synthesis vessel. Fmoc-Lys (Boc)-OH was dissolved in dry DCM (1eq.) and added to the resin and shaken for 5min after which 2eq of DIEA was added and incubated for 1h with shaking at room temperature. The resin was washed 3X with dry DCM, washed 1X with MeOH, and rinsed with DMF. Next, the Fmoc protecting group was removed using 20% piperidine in DMF for 30 min. The resin was washed using alternating DCM and MeOH and finally rinsed with DMF. Next, Fmoc-Lys(Boc)-OH was coupled to the resin using 4 equivalents of the amino acid and 4 equivalents each of Oxyma and DIC; the coupling reaction was done at room temperature with shaking for 2 h. The Fmoc group was similarly removed using 20% piperidine and Following wash steps as described above, Fmoc-Lys($N_3$)-OH was similarly coupled to Lys(Boc) to make Fmoc-Lys($N_3$)-Lys(Boc)-trityl chloride resin. The peptide was cleaved off the resin using 95% TFA:2.5% TIPS: 2.5% $H_2O$ and purified by preparative RP-HPLC on a on a Phenomenex Luna 10 µm C8(2) 100 Å (250 x 21.2 mm) using a 20 to 100% linear gradient of MeCN in $H_2O$ each containing 0.1% TFA at 10 mL/min. ESI-MS $[M+H]^+$ m/z calcd. 523.2663 for $C_{27}H_{35}N_6O_5$, obsd. 523.2669. $^1H$ NMR (600 MHz, $CD_3CN$) δ 1.30-1.40 (m, 2H), 1.50-1.65 (m, 3H), 1.72-1.79 (m, 1H), 2.90 (t, J=6 Hz, 1H), 3.26 (t, J=6 Hz, 1H), 4.09 (dd, J=6, 8 Hz, 1H), 4.21- 4.24 (m, 1H), 4.28 (dd, J= 6Hz, 1H), 4.35 (dd, J= 6Hz, 1H), 6.31 (brs, 0.5H), 7.16 (brs, 1H), 7.33 (t, J= 6Hz, 2H), 7.41 (t, J= 6Hz, 2H), 7.67 (brt, J= 6Hz, 2H), 7.82 (d, J= 6Hz, 2H). $^{13}C$ NMR (151 MHz, $CD_3CN$) δ 22.7, 23.5, 26.9, 29.0,

31.1, 32.4, 40.4, 47.9, 51.9, 53.0, 55.7, 67.4, 120.9, 120.9, 126.2, 126.2, 128.0, 128.1, 128.6, 142.0, 144.9, 145.0, 157.3.

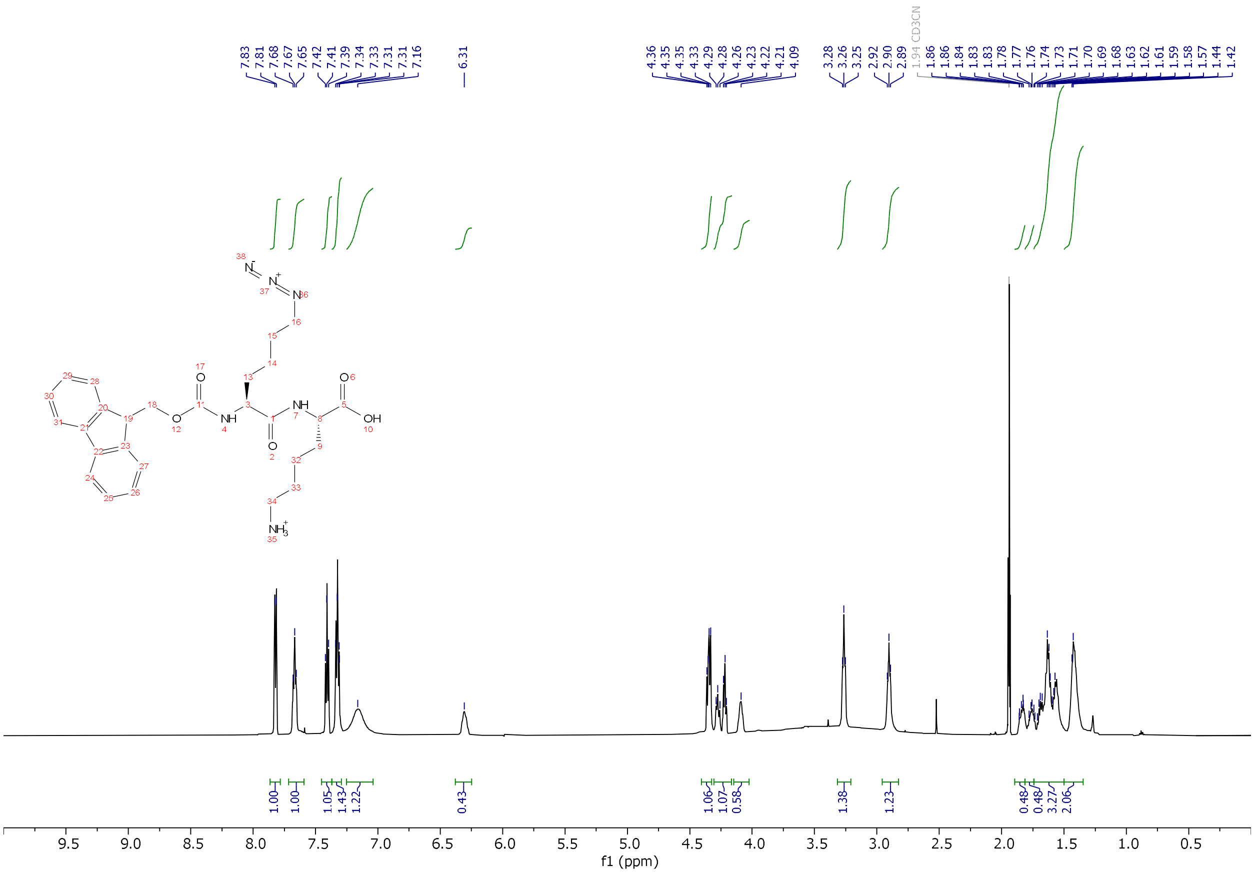

[1]H-NMR of **2n** in $CD_3CN$.

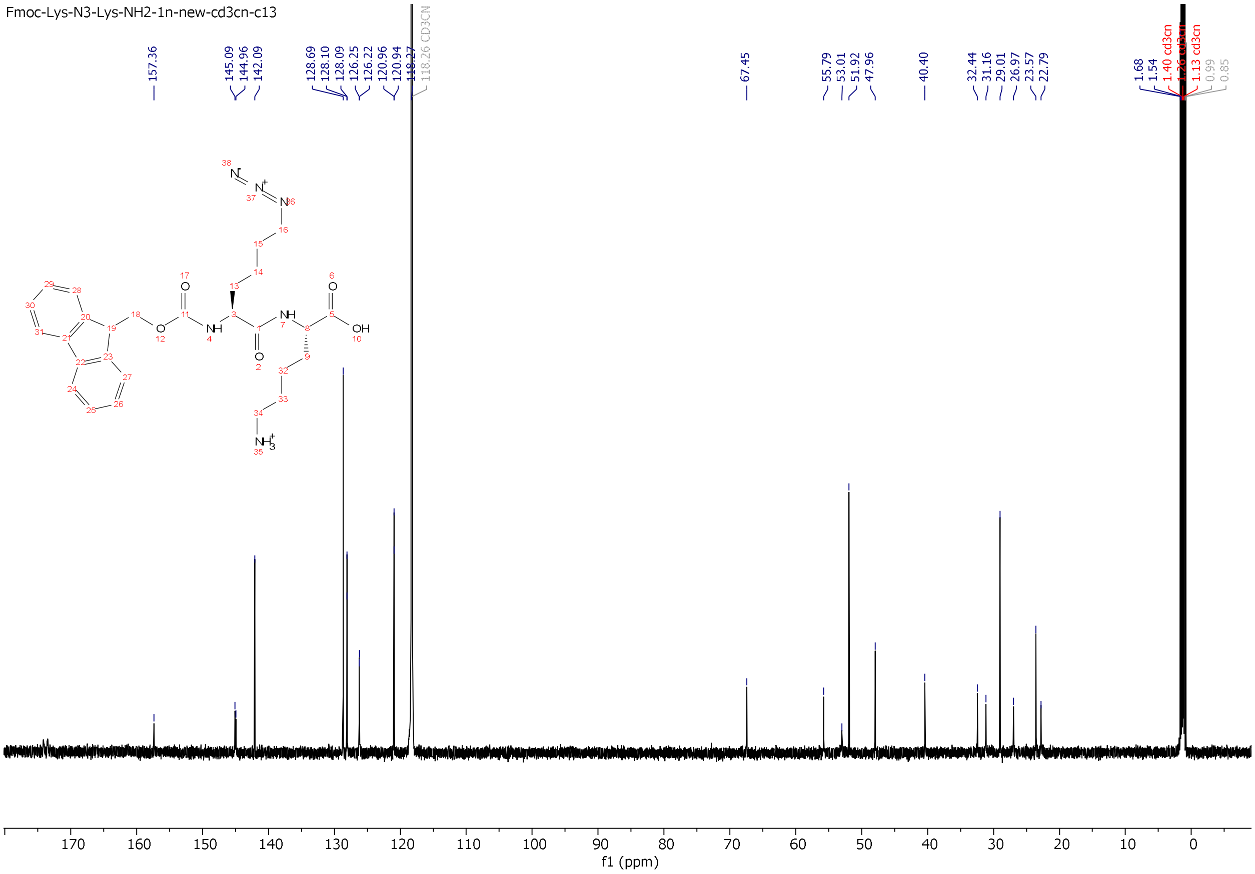


$^{13}$C-NMR of **2n** in $CD_3CN$.

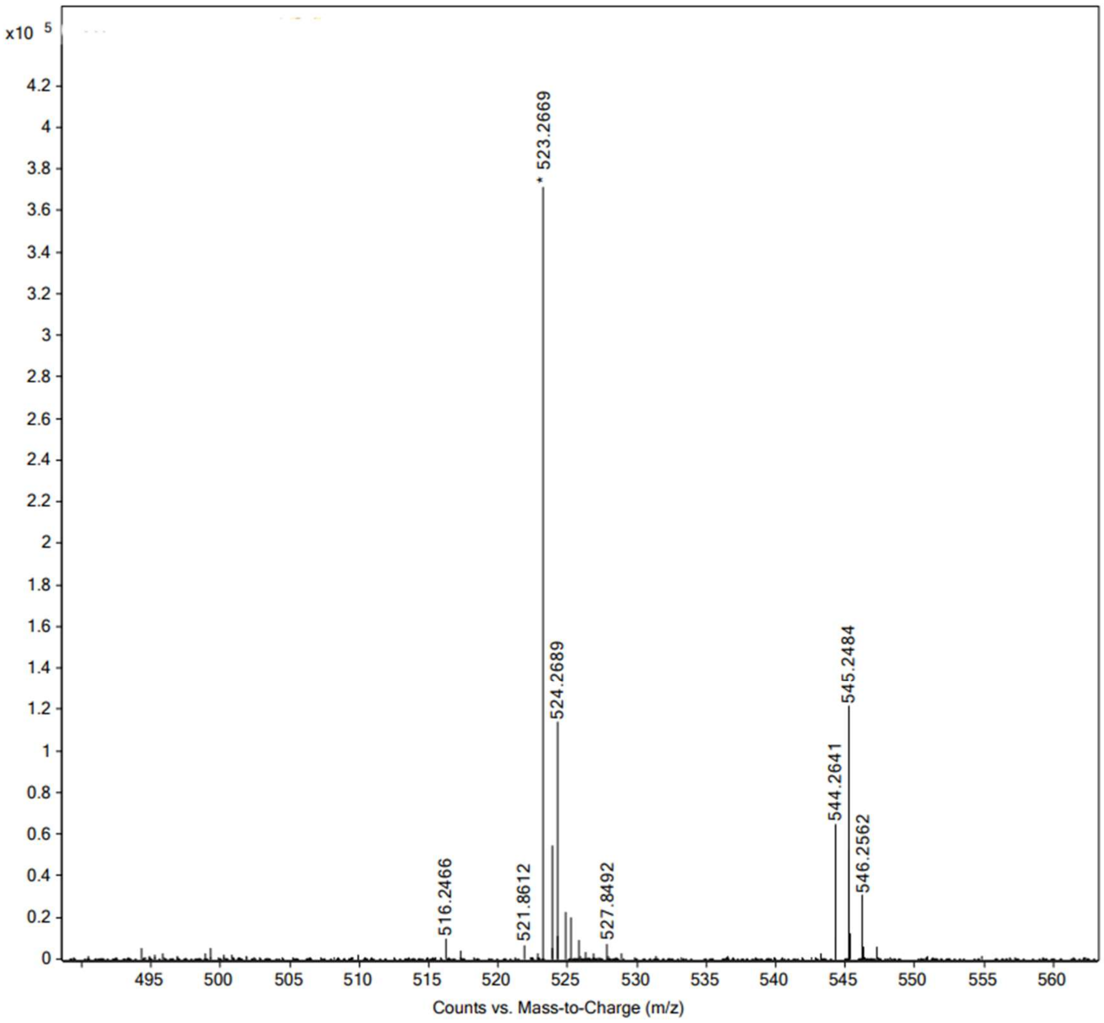


High resolution mass spectrum (ESI) of **2n**.

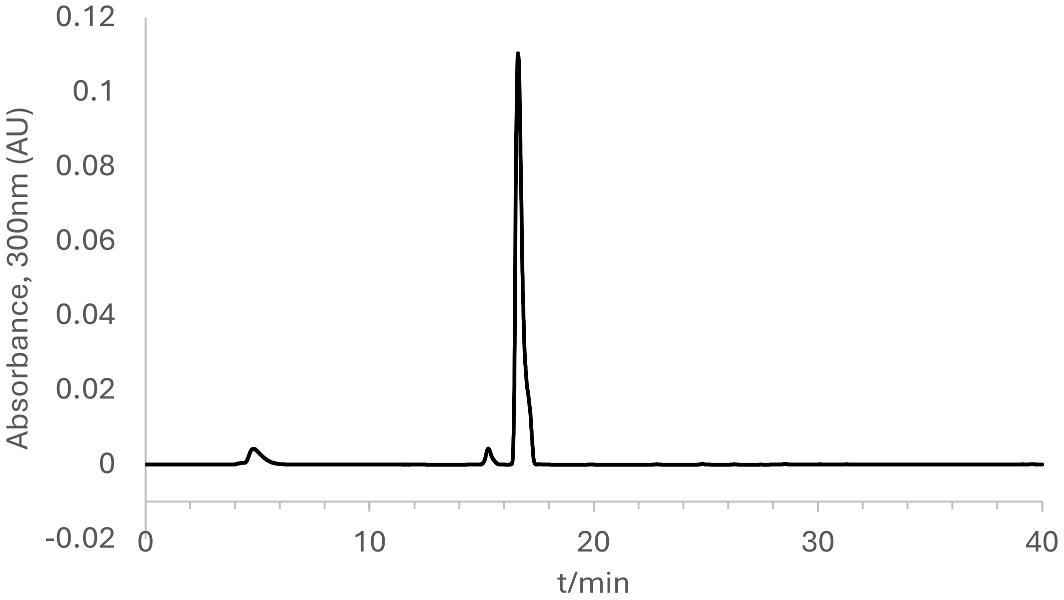


Analytical HPLC chromatogram of **2n**.

**Synthesis of 3p**.

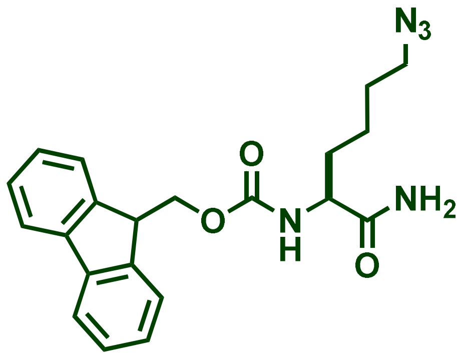

100 mg of Fmoc-aminomethyl)-phenoxy resin (Rink amide resin) of loading capacity 0.48 mmol/g was swollen in DCM after which the Fmoc protecting group was removed using 20% piperidine in DMF for 30min. The resin was washed using alternating DCM and MeOH and finally rinsed with DMF. Fmoc-Lys($N_3$)-OH was coupled to the resin using 4 equivalents of the amino acid and 4 equivalents each of Oxyma and DIC; the coupling reaction was done at room temperature with shaking for 2h. The amidated amino acid was cleaved off the resin using 95% TFA:2.5% TIPS: 2.5% $H_2O$ and purified by preparative RP-HPLC on a on a Phenomenex Luna 10 µm C8(2) 100 Å (250 x 21.2 mm) using a 20 to 100% linear gradient of MeCN in $H_2O$ each containing 0.1% TFA at 10 mL/min. ESI-MS $[M+H]^+$ m/z calcd. 394.1874 for $C_{21}H_{24}N_5O_3$ obsd. 394.1876. $^1$H NMR (600 MHz, $CD_3CN$) δ 1.30-1.40 (m, 1H), 1.50-1.65 (m, 2H), 1.72-1.82 (m, 1H), 2.90 (brs), 3.29 (t, J=6 Hz, 1H), 3.99 (brs, 1H), 4.24 (t, J= 6Hz, 1H), 4.33 (t, J= 6Hz, 1H), 4.38 (dd, J= 6Hz, 1H), 5.78 (brs, 0.5H), 5.91 (brs, 0.5H), 6.33 (brs, 0.5H), 7.16 (brs, 1H), 7.34 (dt, J= 1 and 6Hz, 2H), 7.42 (t, J= 6Hz, 2H), 7.68 (brt, J= 6Hz, 2H), 7.83 (d, J= 6Hz, 2H).

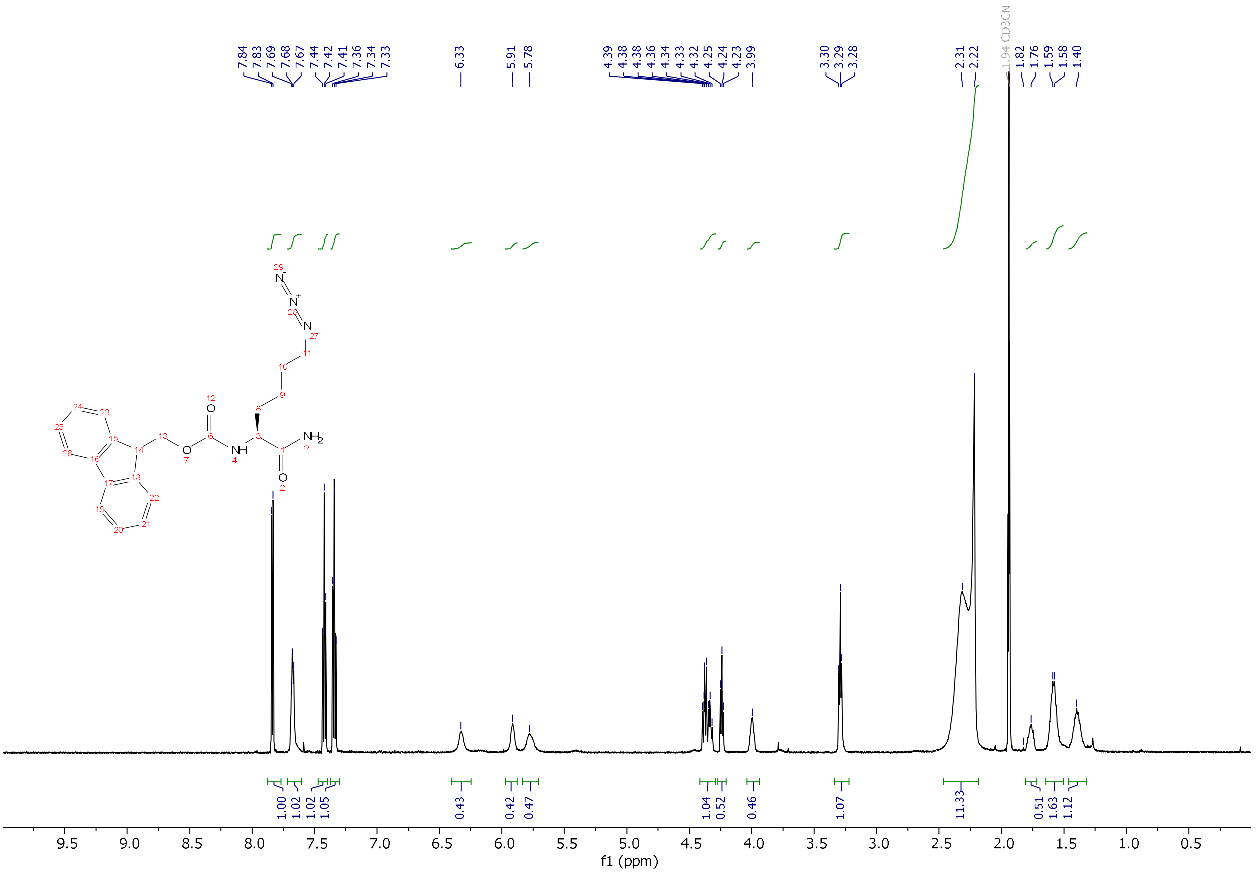

[1]H-NMR of **3p** in $CD_3CN$.

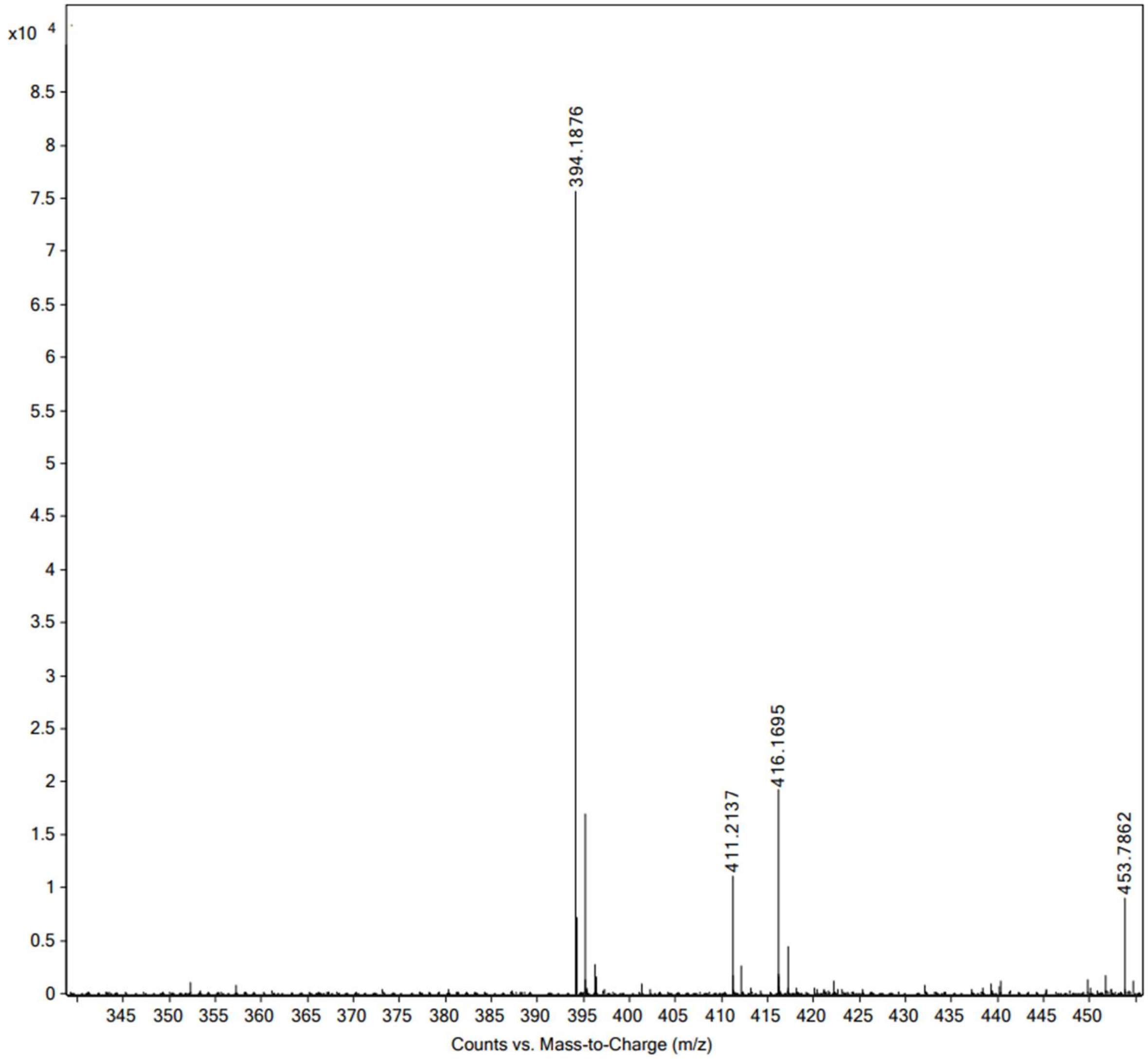


High resolution mass spectrum (ESI) of **3p**.

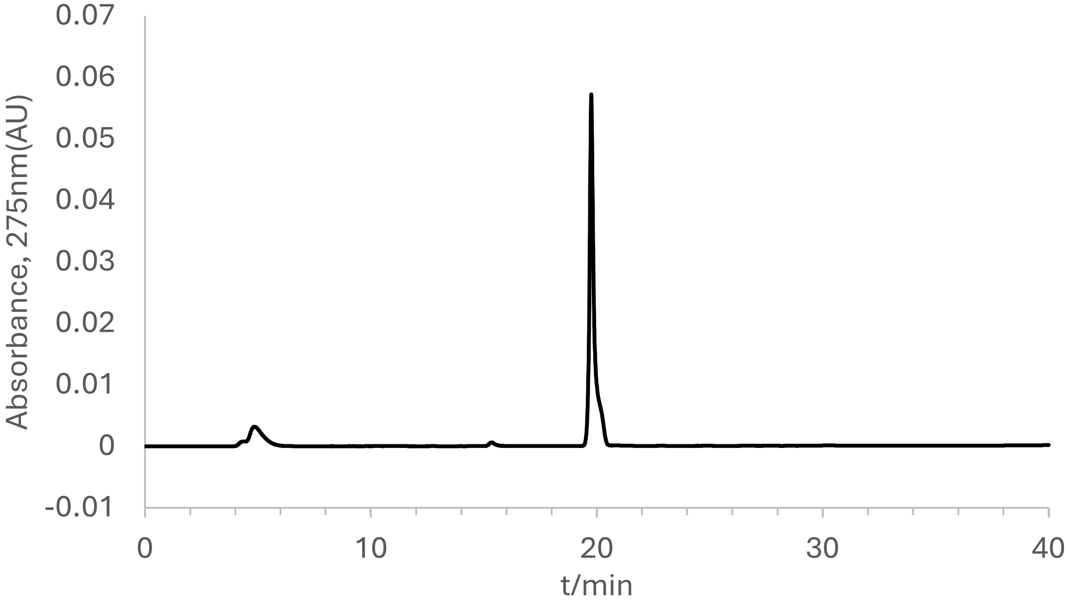


Analytical HPLC chromatogram of **3p**.

**Synthesis of 4a**.

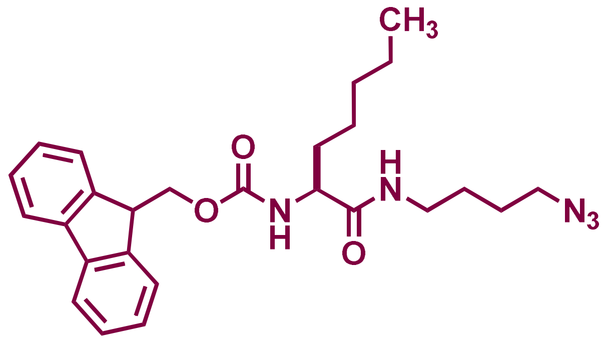


4-azidobutylamine was coupled to Fmoc-Nle-OH using 2 equivalents of the amino acid, 2 eq. of HBTU and 4 eq. DIEA in DMF. The coupling reaction was done at room temperature with agitation for 2 h to make Fmoc-Nle-butyl azide. Excess DMF solvent was removed by rotary evaporation as a DMF-toluene azeotrope until dry powder was obtained. The powder was resuspended in nanopure water to make a white suspension. The suspension was centrifuged at 4000g for 10min to separate the coupling reagents (aqueous layer) from the product (pellet). The pellet was resuspended in 20% HPLC grade acetonitrile in water (0.1% TFA) and injected into preparative RP-HPLC for purification on a Phenomenex Luna 10 µm C8(2) 100 Å (250 x 21.2 mm) using a 20 to 100% linear gradient of MeCN in $H_2O$ each containing 0.1% TFA at 10 mL/min. ESI-MS $[M+H]^+$ m/z calcd. 450.2500 for $C_{25}H_{32}N_5O_3$, obsd. 450.2500 $^1H$ NMR (600 MHz, $CD_3CN$) δ 0.89 (t, 3H, CH3),1.20 (m,1H), 1.25-1.39 (m, 2H), 1.47-1.59 (m, 4H), 1.72 (m, 1H), 3.16 (dd, J= 6 and 9 Hz, 2H), 3.28 (t, J= 6 Hz, 1H), 3.95 (brs, 1H), 4.23 (t, J= 6Hz, 1H), 4.32 (brt, J= 6Hz, 1H), 4.36 (dd, J= 2 and 6Hz, 1H), 5.90 (brs, 1H), 6.66 (brs, 1H), 7.34 (t, J= 6Hz, 2H), 7.42 (t, J= 6Hz, 2H), 7.68 (brt, J= 6Hz, 2H), 7.83 (d, J= 6Hz, 2H).

.

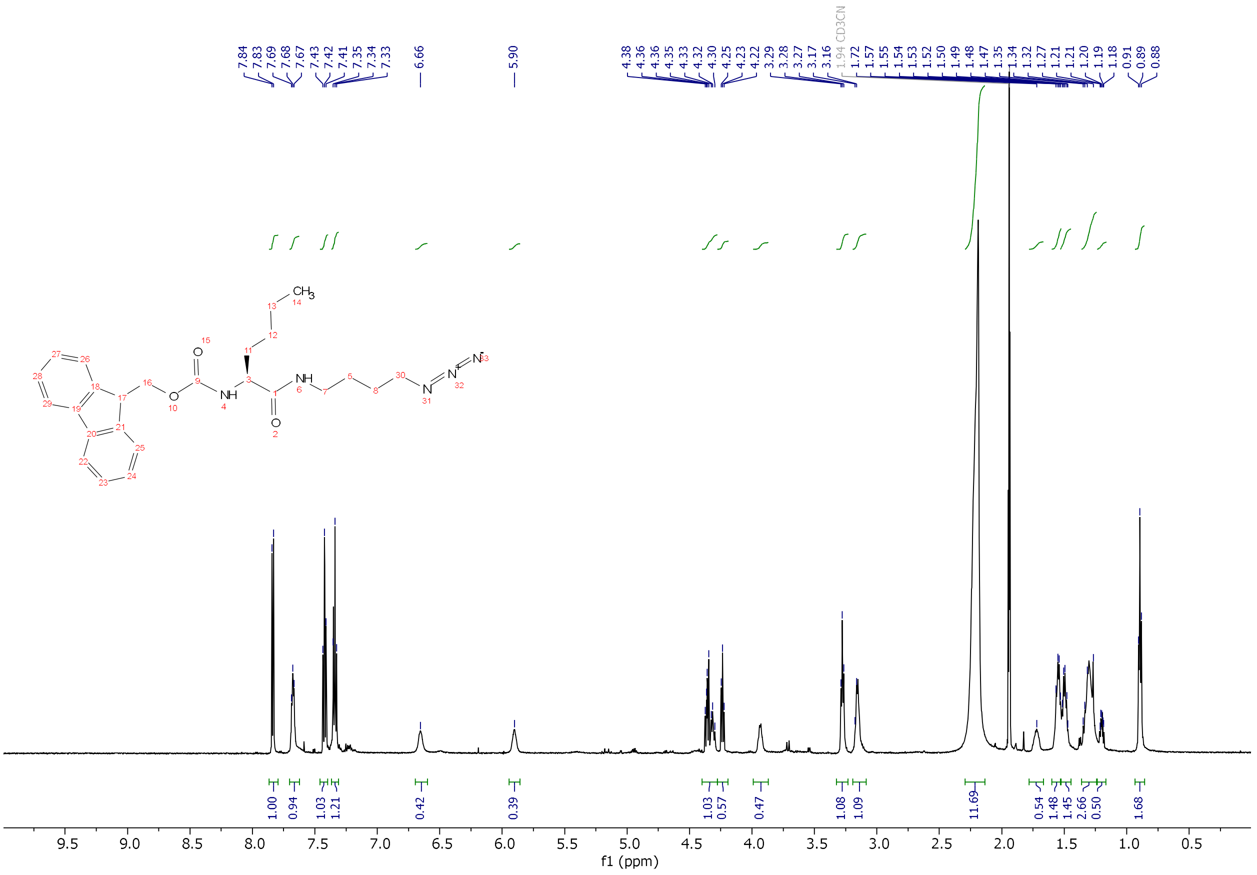

$^1$H-NMR of **4a** in $CD_3CN$.

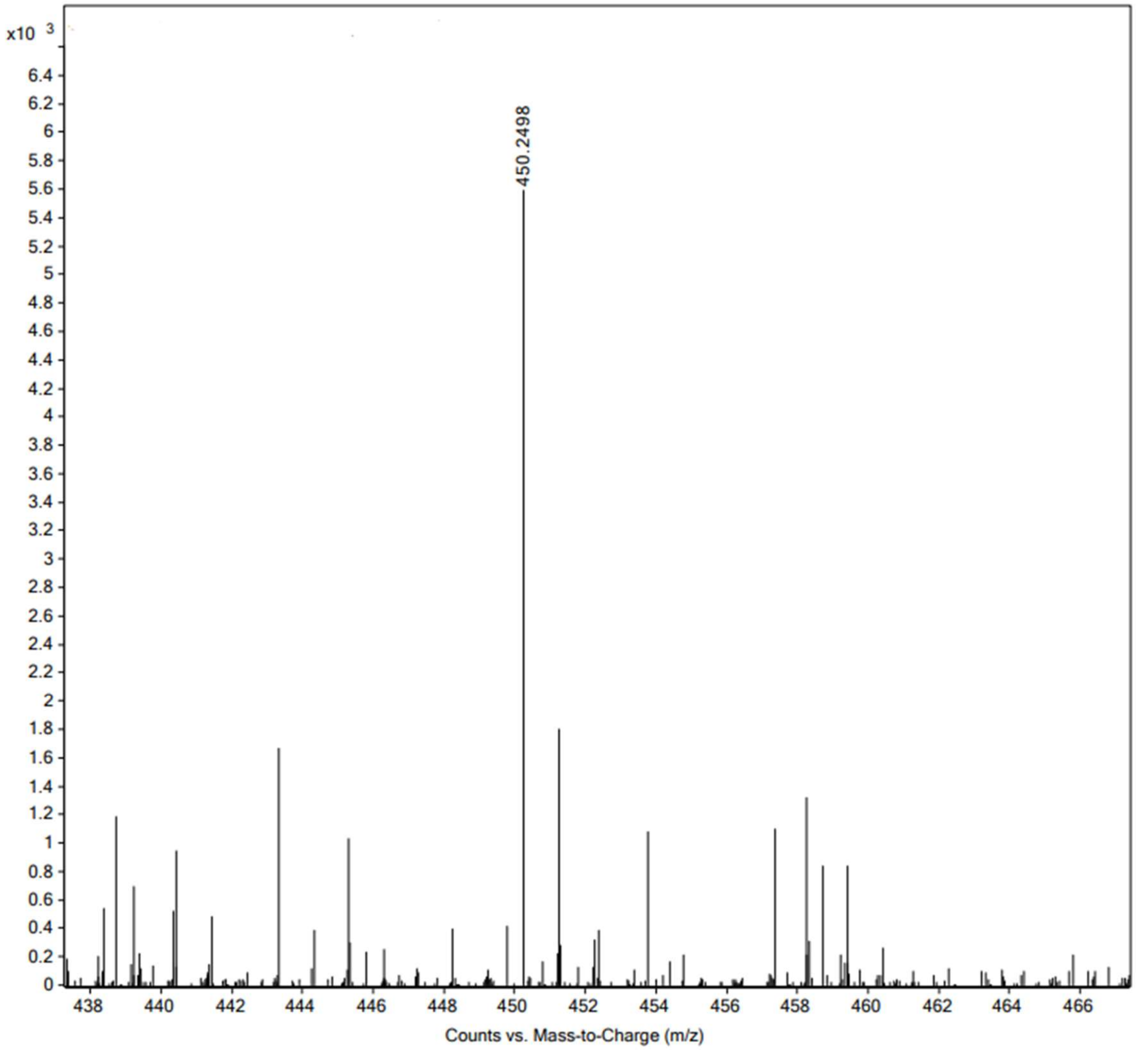


High resolution mass spectrum (ESI) of **4a.**

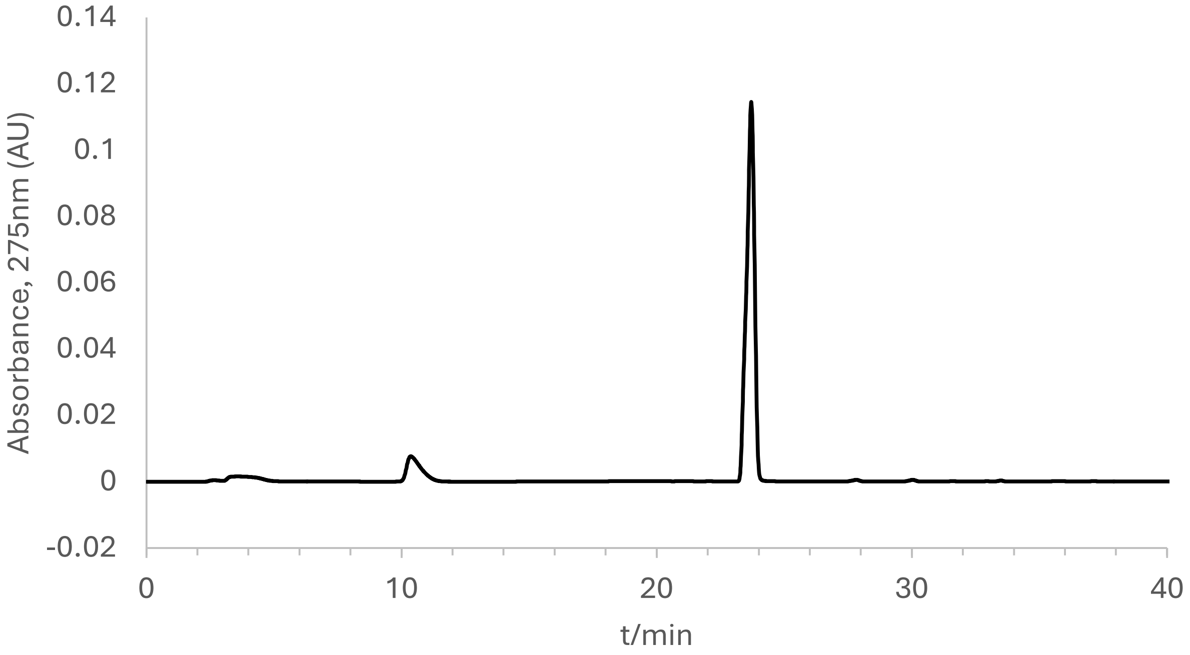


Analytical HPLC chromatogram of **4a**.

**Synthesis of 4b, 4c, 4d, and 4e or Fmoc-Lys $(Me)_x$- butyl azide where x = 0, 1, 2, 3**

4-azidobutylamine was coupled to either Fmoc-Lys (Boc)-OH or Fmoc-Lys (Boc, Me)-OH or Fmoc-Lys$(Me)_2$-OH.HCl or Fmoc-Lys$(Me)_3$-$OH^+Cl^-$ using 2 equivalents of the amino acid, 2 eq. of HBTU and 4 eq. DIEA in DMF. The coupling reaction was done at room temperature with agitation for 2h to link the two building blocks by an amide bond. Excess DMF solvent was removed by rotary evaporation as a DMF-toluene azeotrope until dry powder was obtained. The powder was resuspended in nanopure water to make a white suspension. The suspension was centrifuged at 4000g for 10min to separate the coupling reagents (aqueous layer) from the product (pellet). The pellet was resuspended in 20% HPLC grade acetonitrile in water (0.1% TFA) and injected into preparative RP-HPLC for purification on a Phenomenex Luna 10 µm C8(2) 100 Å (250 x 21.2 mm) using a 20 to 100% linear gradient of MeCN in $H_2O$ each containing 0.1% TFA at 10 mL/min.

**4b.**

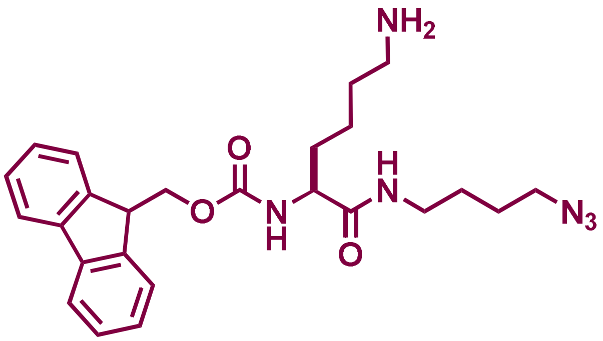


ESI-MS $[M+H]^+$ m/z calcd. 465.2609 for $C_{25}H_{33}N_6O_3$, obsd. 465.2609. $^1H$ NMR (600 MHz, $cdcl_3$) δ 1.30-171 (m, 3H), 2.91 (brs, 1H), 3.14 (m, 2H), 4.09 (t, J= 6Hz, 1H), 4.27 (m, 1H), 6.19 (brd, J=6 Hz, 1H), 7.23 (dt, J= 1 and 6Hz, 2H), 7.33 (t, J= 6Hz, 2H), 7.51 (brd, J= 6Hz, 2H), 7.69 (d, J= 6Hz, 2H), 7.99 (brs, 1H). $^{13}C$ NMR (151 MHz, $cdcl_3$) δ 22.15, 26.01, 26.39, 26.69, 31.90, 38.94, 39.46, 46.89, 50.81, 54.55, 67.28, 120.00, 125.04, 127.06, 127.80, 141.19, 143.62, 156.67, 161.57, 161.80, 172.27.

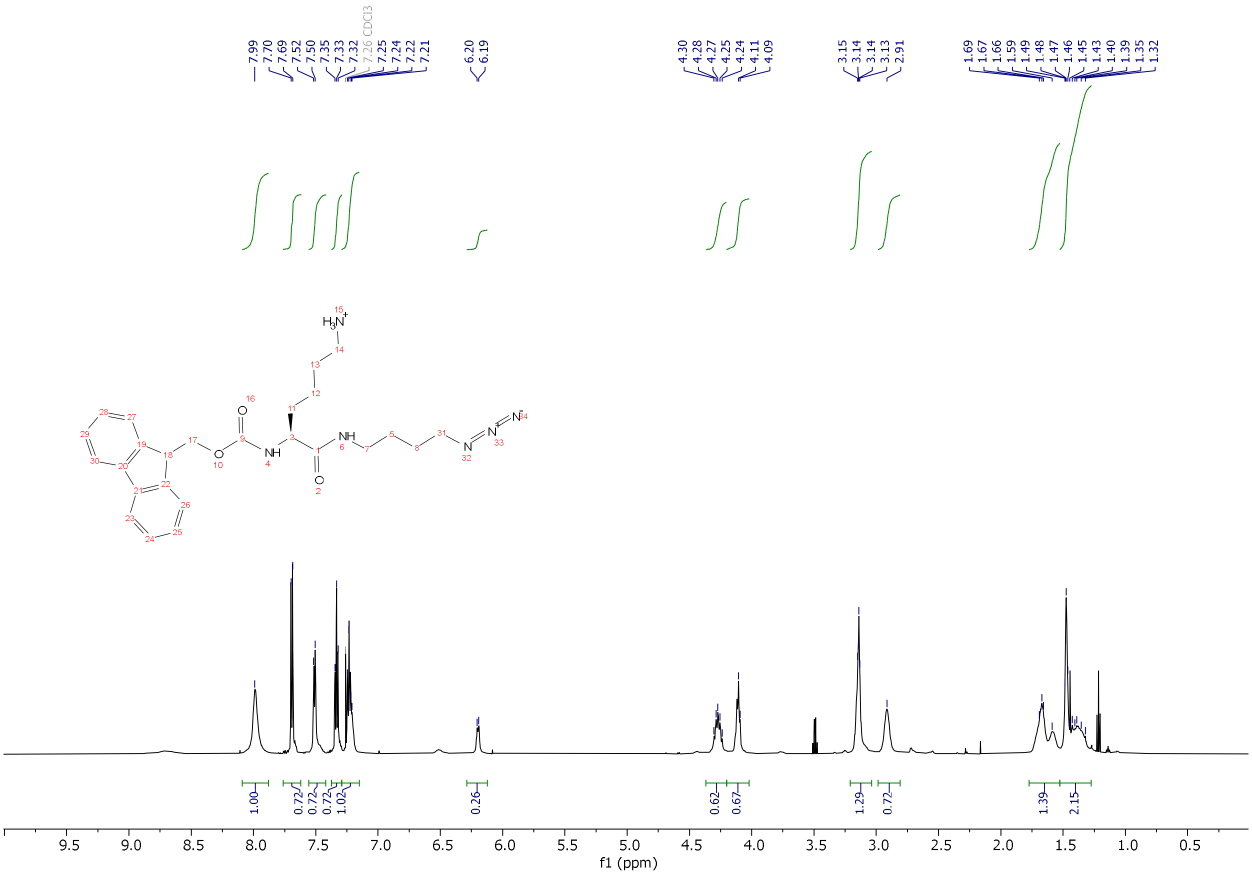


$^1$H-NMR of **4b** in $CDCl_3$.

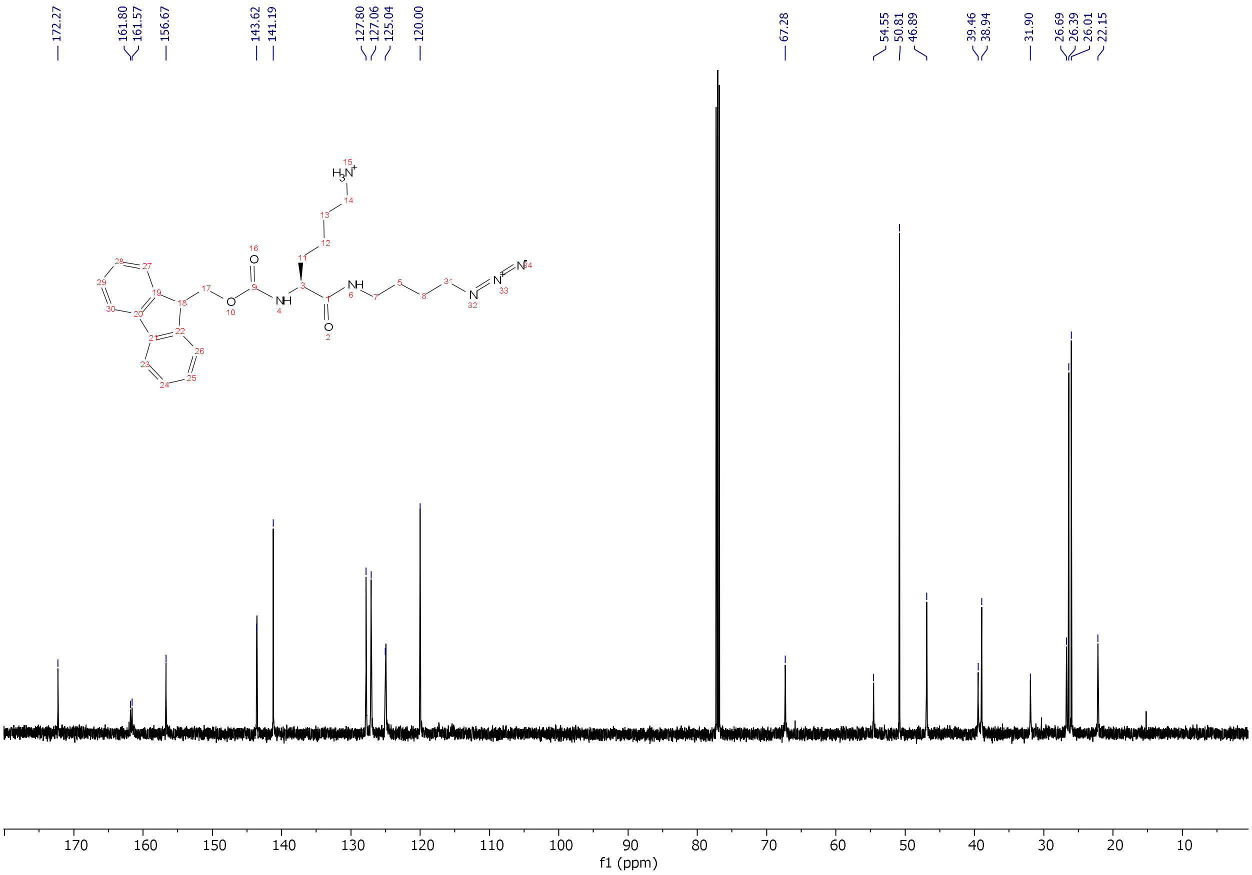


$^{13}$C-NMR of **4b** in $CDCl_3$.

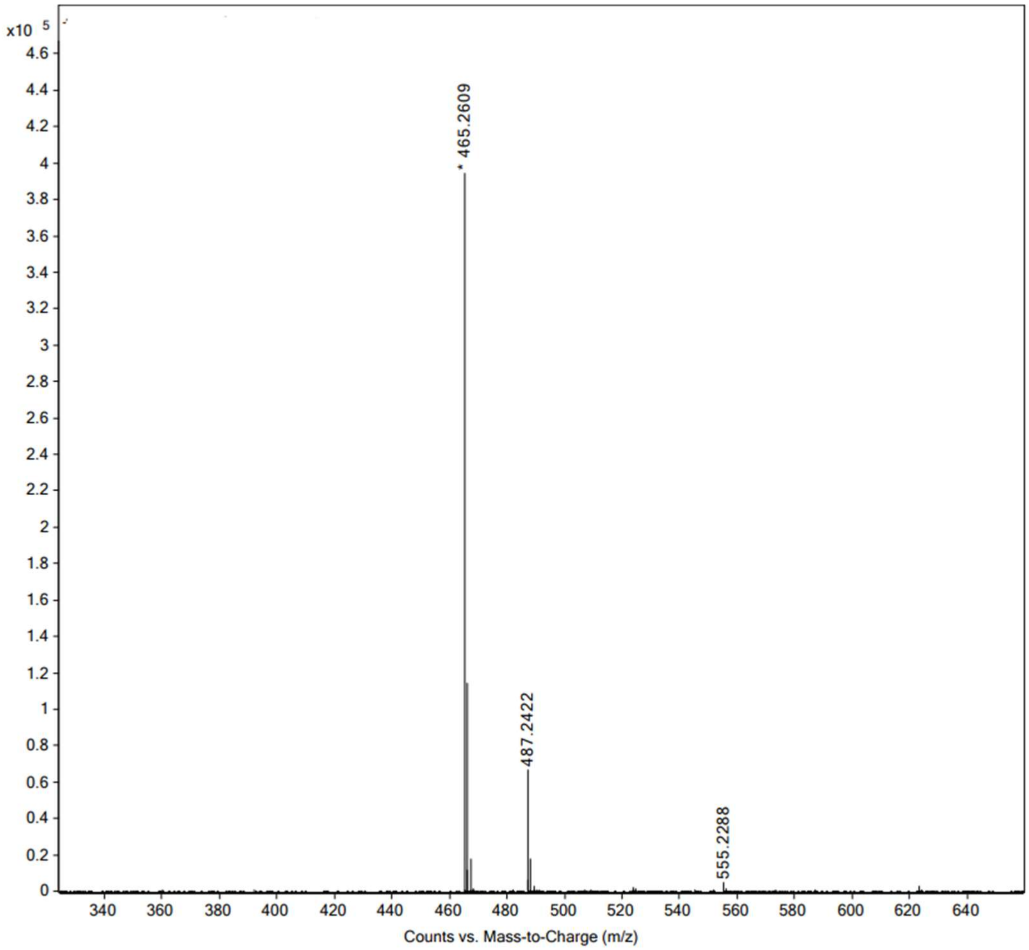


High resolution mass spectrum (ESI) of **4b**.

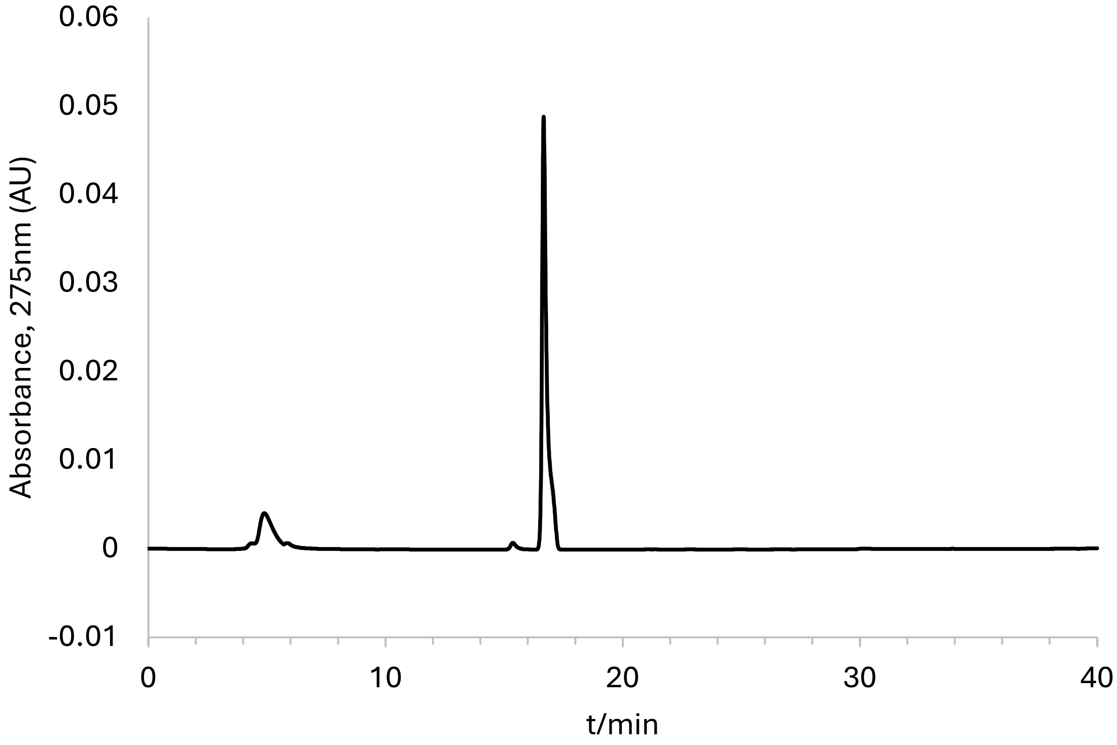


Analytical HPLC chromatogram of **4b**.

**4c.**

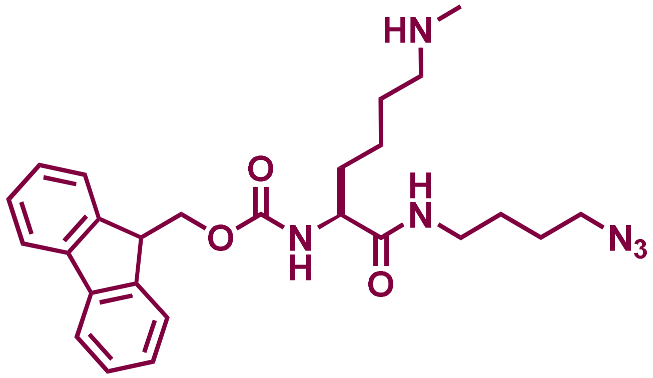


ESI-MS $[M+H]^+$ m/z calcd. 479.2765 for $C_{26}H_{35}N_6O_3$, obsd. 479.2779. $^1$H NMR (600 MHz, $CDCl_3$) δ 1.43 (m, 1H), 1.54 (br, 2H), 1.75-1.82 (m, 3H), 2.88 (m, 1H), 2.99 (m, 1H), 3.21 (brs, 1H), 4.17 (q, J= 6Hz, 2H), 4.30 (m, 1H), 6.01 (brd, J=6 Hz, 1H), 7.24 (brs, 1H), 7.27 (t, J= 6Hz, 2H), 7.38 (t, J= 6Hz, 2H), 7.54 (brd, J= 6Hz, 2H), 7.74 (d, J= 6Hz, 2H), 9.05 (brs, 1H), 9.39 (brs, 1H).

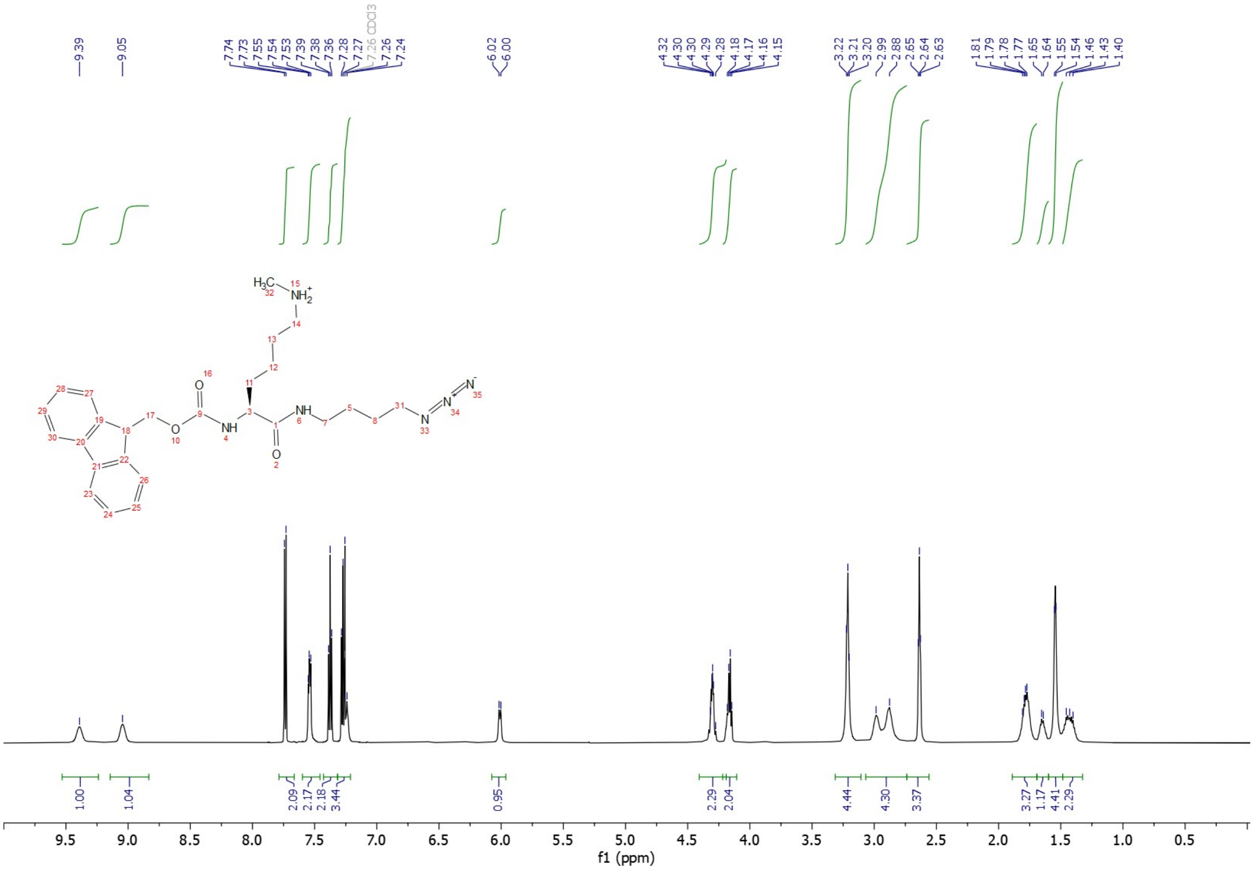

[1]H-NMR of **4c** in $CDCl_3$.

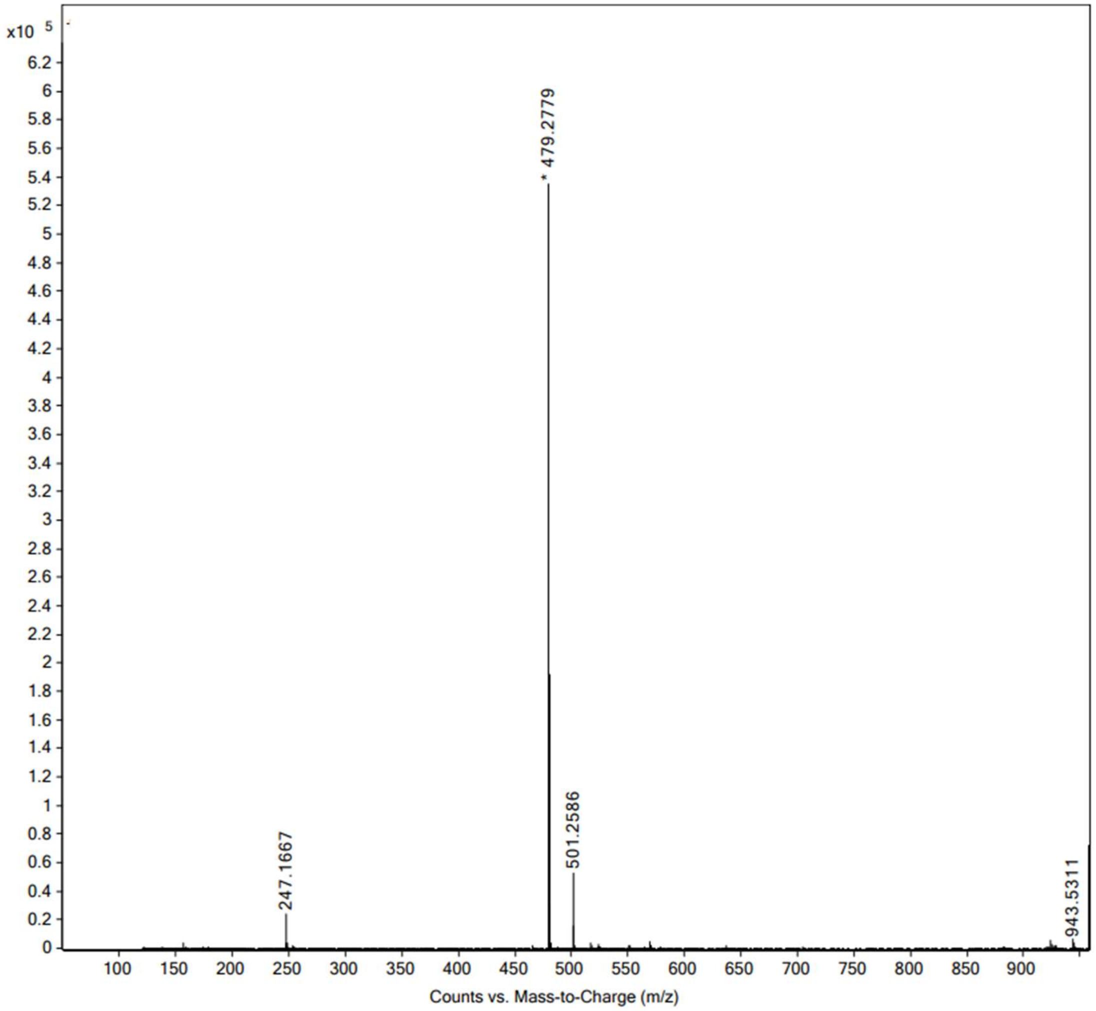


High resolution mass spectrum (ESI) of **4c**.

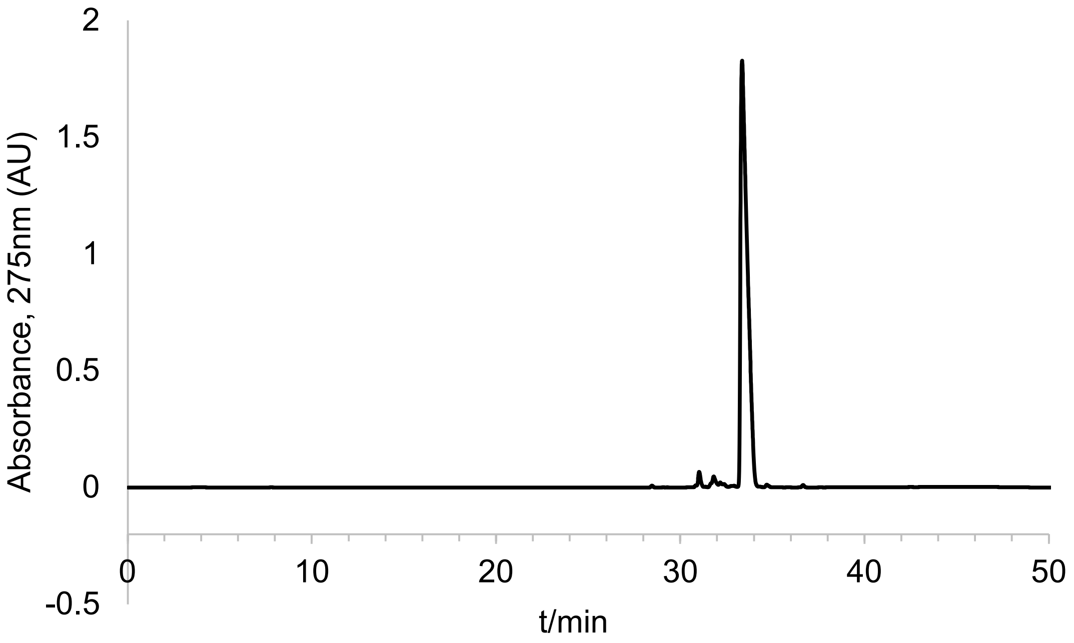


Analytical HPLC chromatogram of **4c.**

**4d.**

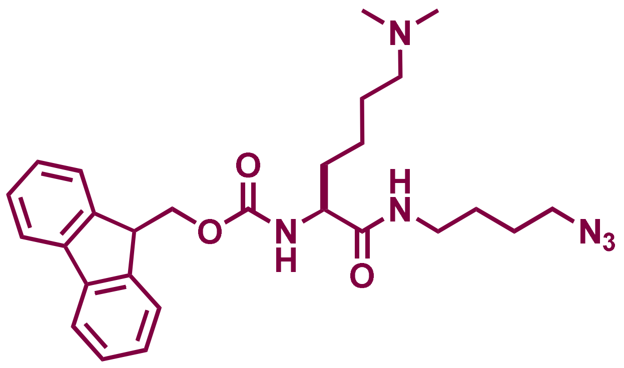

ESI-MS [M+H]+ m/z calcd. 493.2922 for $C_{27}H_{37}N_6O_3$, obsd. 493.2926. $^1H$ NMR (600 MHz, cdcl$_3$) δ 1.45 (m, 1H), 1.59 (m, 2H), 1.70 (m, 1H), 1.82 (m, 2H), 2.82 (brs, 6H), 3.10 (brs, 1H), 4.20 (t, J = 6Hz, 1H), 4.21 (m, 1H), 4.32 (m, 1H), 6.00 (brs, 1H), 7.30 (t, J= 6 Hz, 2H), 7.40 (t, J= 6 Hz, 2H), 7.59 (t, J= 6 Hz, 2H), 7.76 (d, J= 6 Hz, 2H), 10.33 (brs, 1H), 11.13 (brs, 1H). $^{13}C$ NMR (151 MHz, cdcl$_3$) δ 21.91, 23.52, 26.20, 26.55, 32.00, 39.11, 42.72, 43.47, 47.15, 51.05, 53.96, 57.71, 67.23, 114.89, 116.81, 120.10, 125.24, 127.18, 127.87, 141.37, 143.90, 156.58, 160.76, 161.01, 172.37.

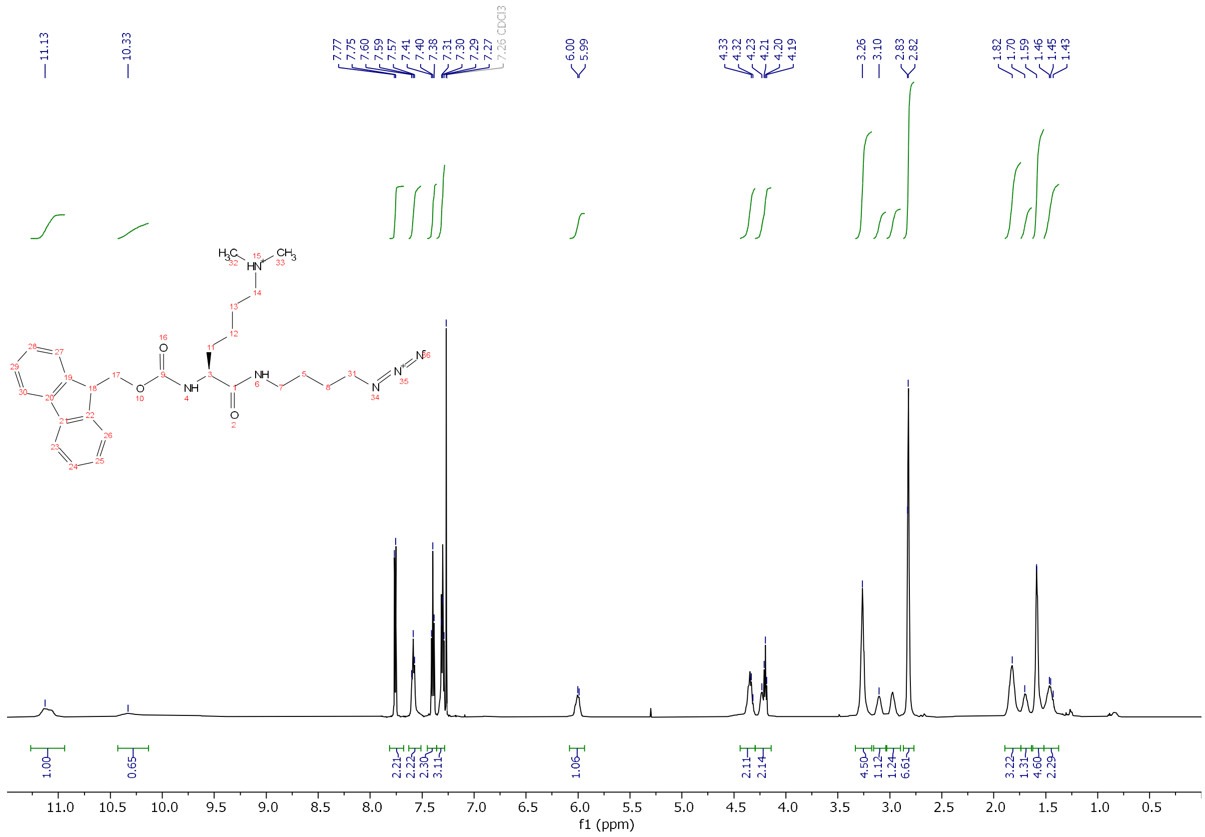

[1]H-NMR of **4d** in $CDCl_3$.

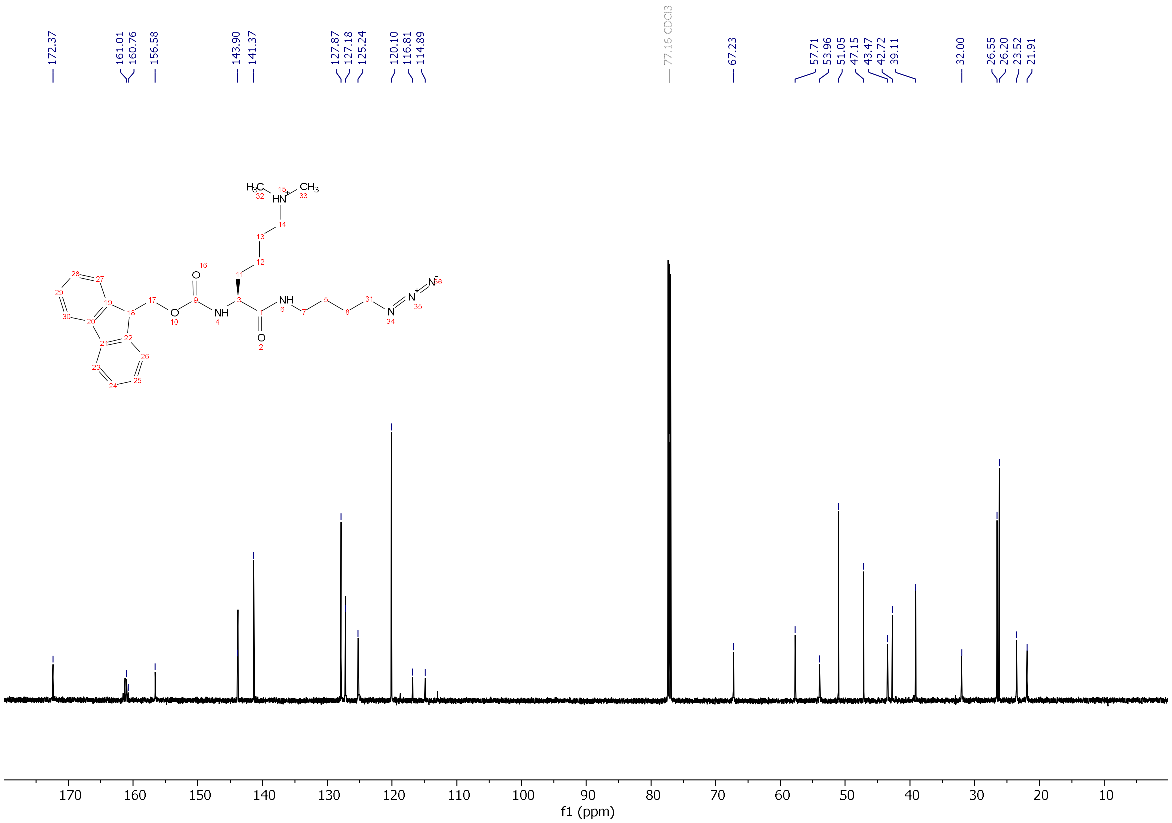


[13]C-NMR of **4d** in $CDCl_3$.

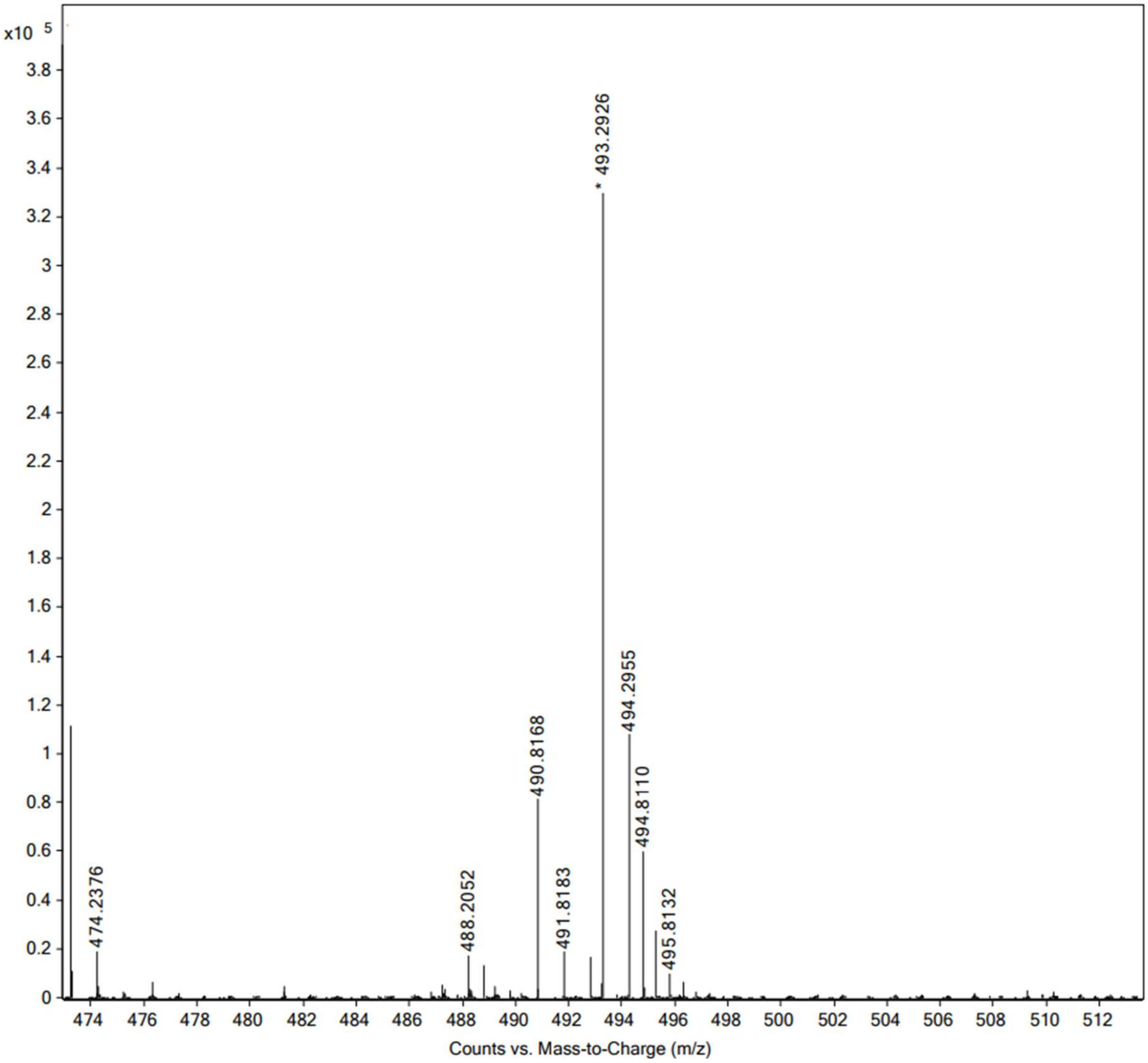


High resolution mass spectrum (ESI) of **4d**.

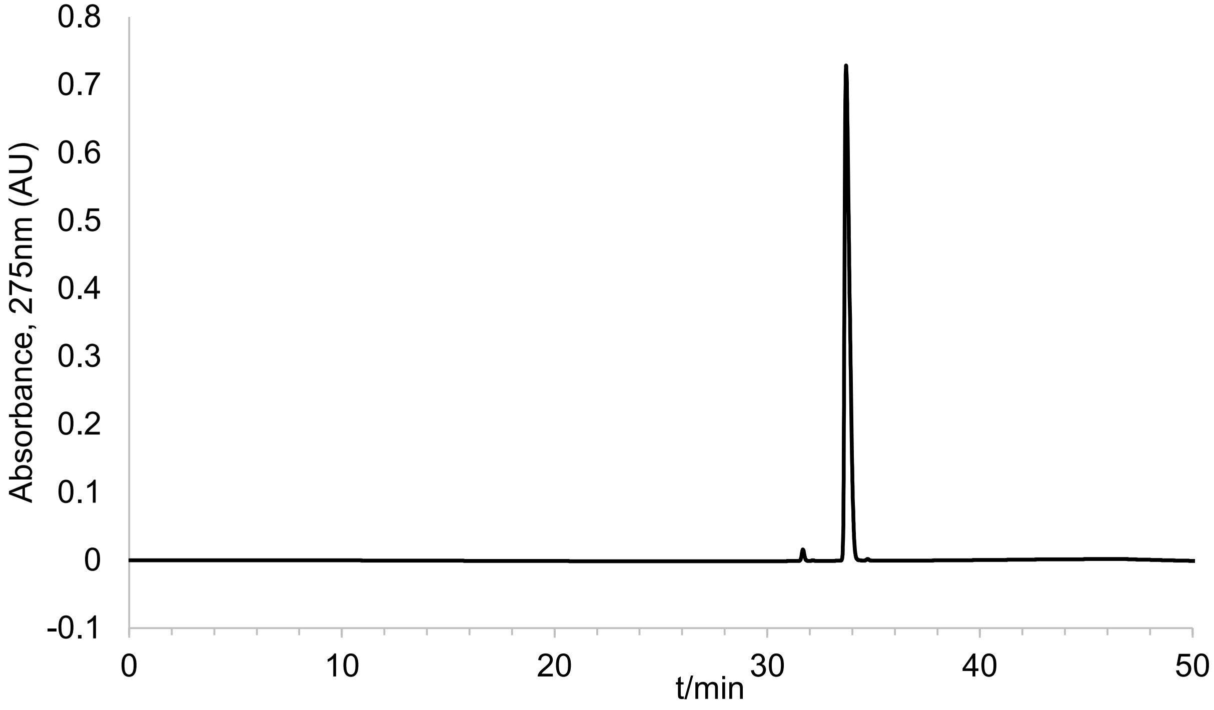


Analytical HPLC chromatogram of **4d**.

**4e.**

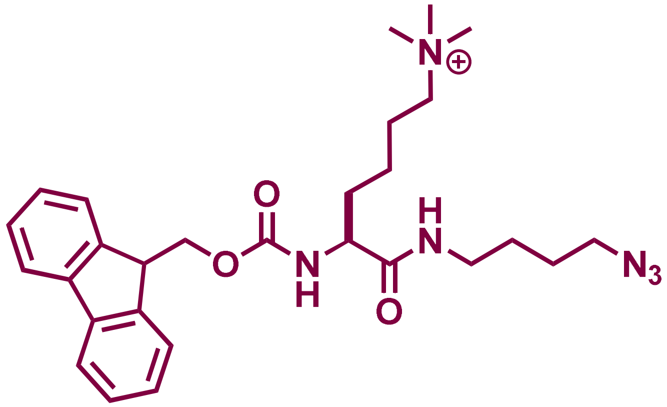


ESI-MS [M+H]+ m/z calcd. 507.3078 for $C_{28}H_{39}N_6O_3$, obsd. 507.3081. $^1$H NMR (600 MHz, $CDCl_3$) δ 1.52 (m, 1H), 1.64 (m, 2H), 1.84 (m, 1H), 2.02 (m, 1H), 3.27 (t, J= 6Hz, 2H), 3.29, 3.35 (brs, 9H), 3.42 (m, 1H), 3.79 (m, 1H), 4.19 (t, J= 6 Hz, 1H), 4.29 (m, 1H), 4.42 (m,1H), 5.92 (brs, 1H), , 7.30 (dt, J= 1, 6Hz, 2H), 7.38 (dt, J= 1, 6Hz, 2H), 7.60 (d, J= 4, 6Hz, 2H), 7.74 (d, J= 6Hz, 2H).

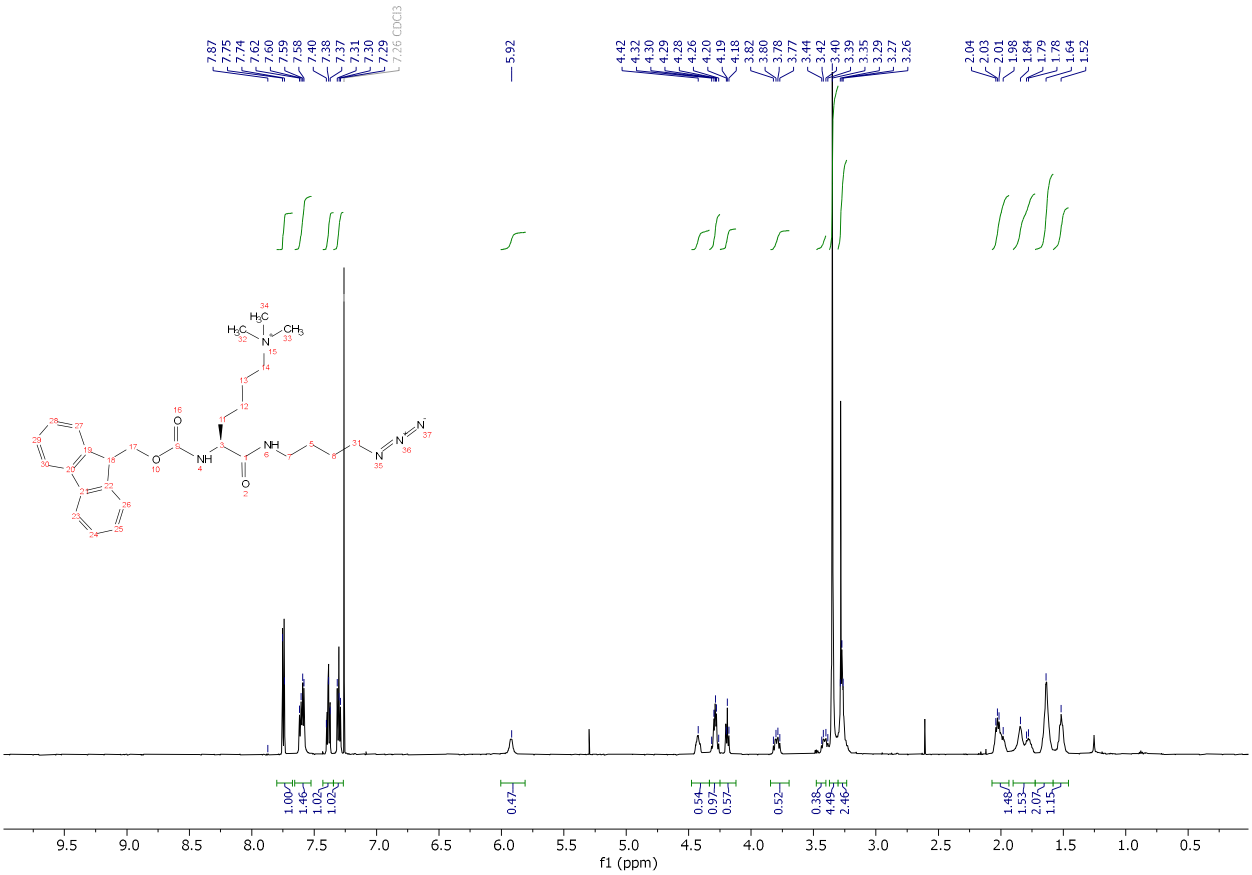

$^1$H-NMR of **4e** in $CDCl_3$.

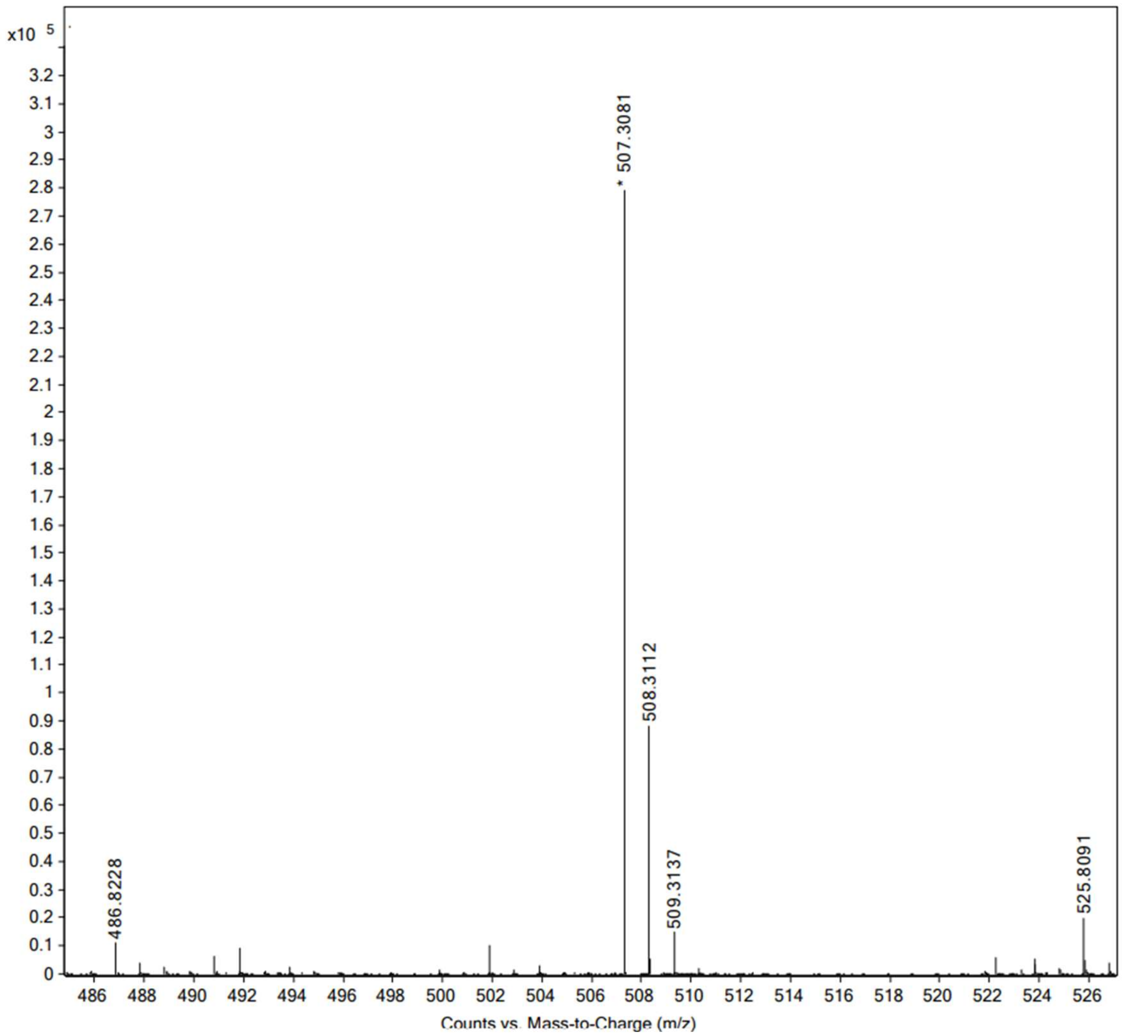


High resolution mass spectrum (ESI) of **4e**.

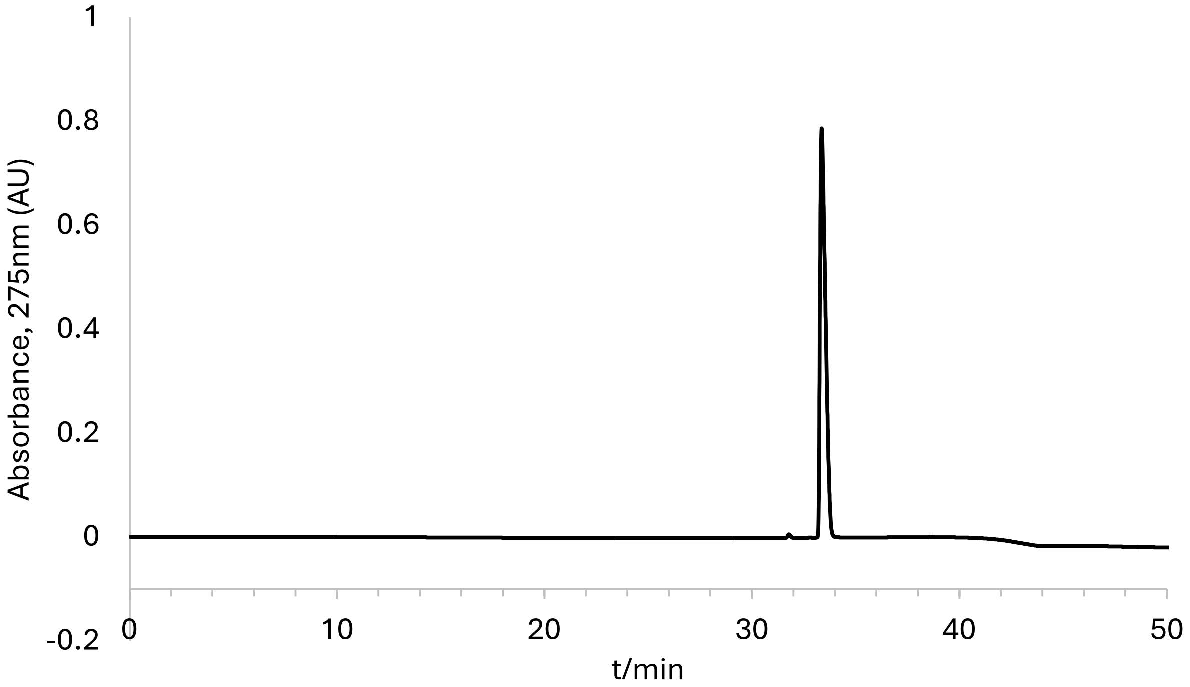


Analytical HPLC chromatogram of **4e**.

**Synthesis of 5a.**

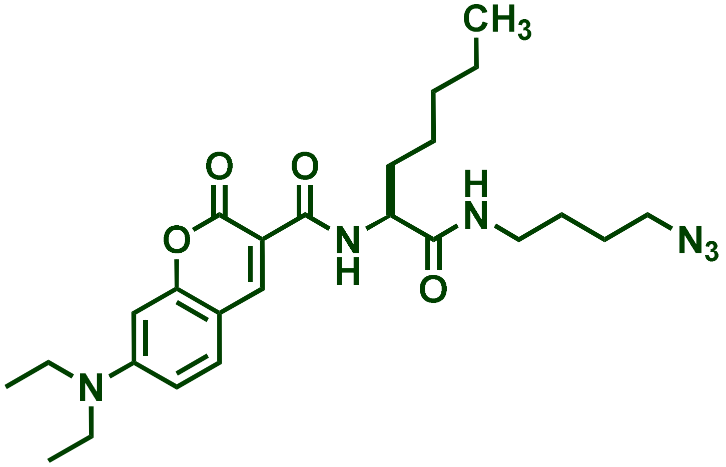

100 mg of chlorotrityl chloride resin (CTC resin) of loading capacity 1.5 mmol/g was swollen in dry DCM after which 1 eq Fmoc-Nle-OH (Nle = norleucine) was loaded on the resin as a solution of 5 mL of dry DCM mixed with 4 eq of dry DIPEA. The resin was shaken for 1hr at room temperature and washed 3X with a cocktail of DCM/MeOH/DIPEA (17:2:1) and then finally with DCM. The Fmoc group was removed using 20% piperidine and following wash steps with DCM and MeOH, 7-(diethylamino) coumarin-3-carboxylic acid was coupled to the norleucine using 4 equivalents of the coumarin and 4 equivalents each of Oxyma and DIC; the coupling reaction was done at room temperature with shaking for 2 h to make coumarin-Nle on trityl chloride resin. The resin was washed using alternating DCM and MeOH and finally rinsed with DCM. This coumarin-Nle conjugate was cleaved off the resin using 1% TFA in dry DCM. The cleavage cocktail was blown off and the resulting oil pellet was washed with ether 3X and dried. 4-azidobutylamine was coupled to the coumarin-Nle conjugate, in solution, using 2 equivalents of azidobutylamine, 2 eq. of HBTU and 4 eq. DIEA in DMF. The coupling reaction was done at room temperature with agitation for 2h to make Coumarin-Nle-butyl azide. Excess DMF solvent was removed by rotary evaporation as a DMF-toluene azeotrope until dry powder was obtained. The powder was resuspended in nanopure water to make a white suspension. The suspension was centrifuged at 4000g for 10min to separate the coupling

reagents (aqueous layer) from the product (pellet). The pellet was resuspended in 20% HPLC grade acetonitrile in water (0.1% TFA) and injected into preparative RP-HPLC for purification on a Phenomenex Luna 10 µm C8(2) 100 Å (250 x 21.2 mm) using a 20 to 100% linear gradient of MeCN in $H_2O$ each containing 0.1% TFA at 10 mL/min. ESI-MS $[M+H]^+$ m/z calcd. 471.2714 for $C_{24}H_{35}N_6O_4$, obsd. 471.2718. $^1H$ NMR (600 MHz, $CDCl_3$) δ 0.89 (t, J= 6.0 Hz, 3H)1.18 (t, J= 6 Hz, 6H), 1.30-1.36 (m, 2H), 1.46-1.59 (m, 4H), 1.66-1.71 (m, 1H), 1.78-1.85 (m, 1H), 3.13-3.22 (m, 2H) 3.29 (t, J= 6 Hz, CH), 3.46 (q, J= 6 Hz, 1H, $CH_2$) , 4.40 (dq, J= 6, 12 Hz, 1H, CH), 6.55 (d, J= 2 Hz, 1H), 6.76 (dd, J= 2 and 12 Hz, 1H), 7.51 (d, J= 12 Hz, 1H), 8.60 (s, 1H), 9.03 (d, J= 12 Hz, 1H). $^{13}C$ NMR (151 MHz, $CDCl_3$) δ 11.67, 13.23, 22.13, 25.83, 26.42, 27.49, 32.31, 32.34, 38.11, 38.23, 44.67, 50.78, 53.55, 53.58, 96.10, 108.05, 109.84, 110.19, 117.30, 131.34, 147.73, 152.92, 157.79, 162.51, 162.53, 171.62.

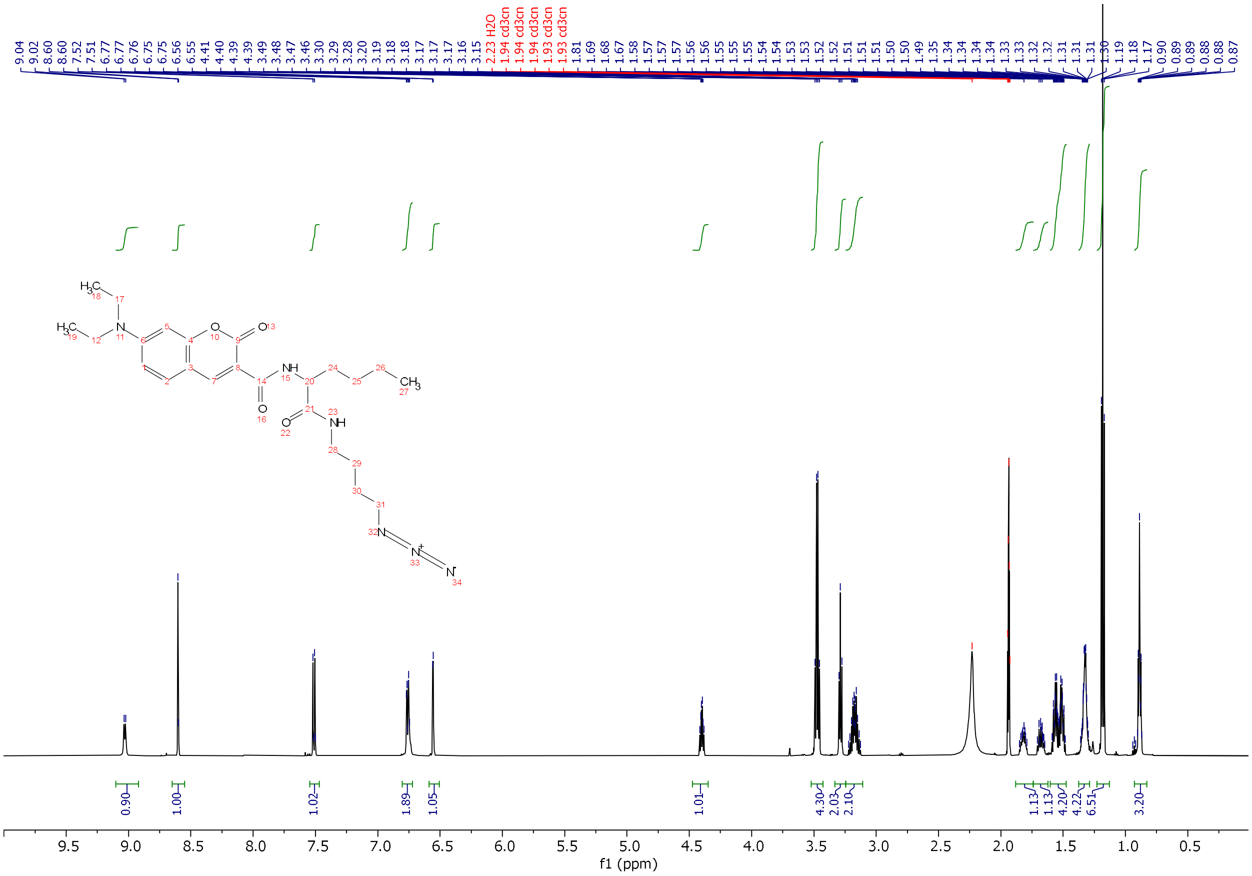

[1]H-NMR of **5a** in $CDCl_3$.

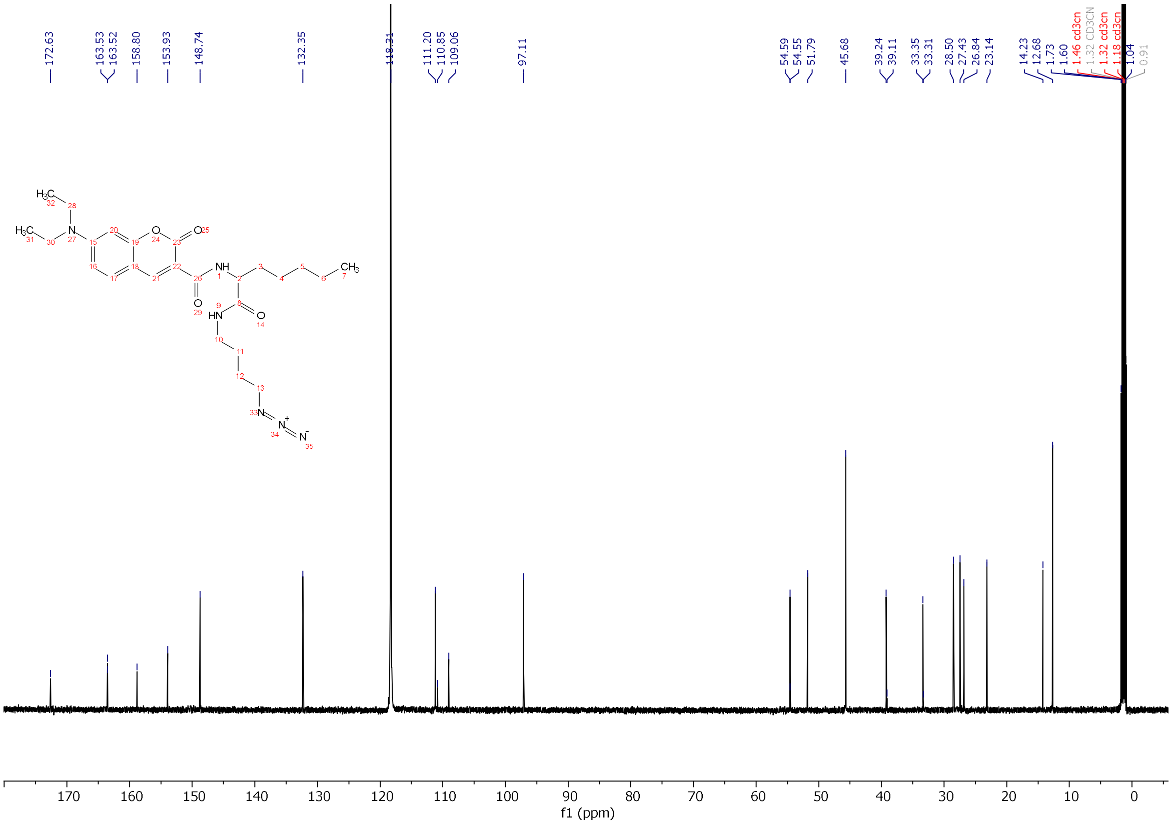


$^{13}$C-NMR of **5a** in $CDCl_3$.

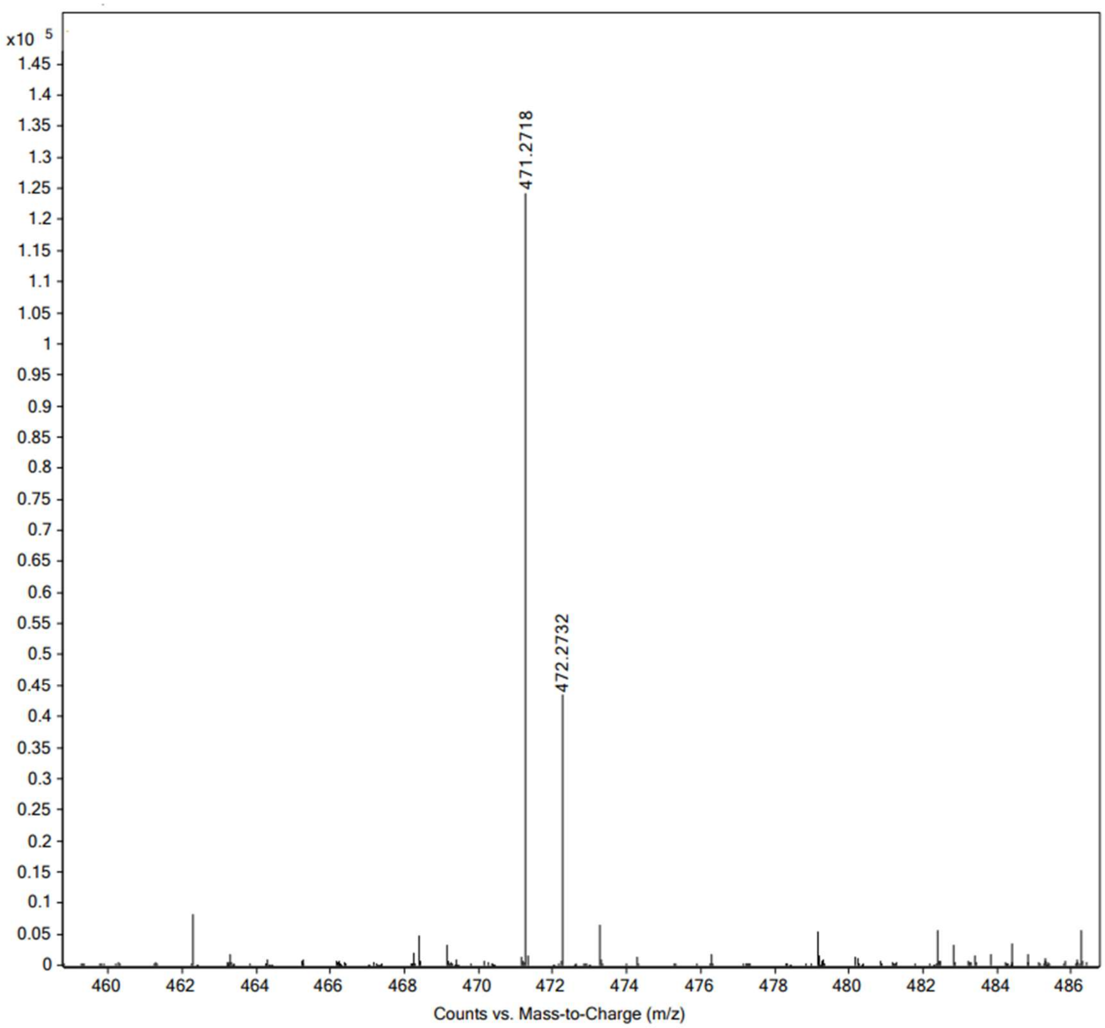


High resolution mass spectrum (ESI) of **5a**

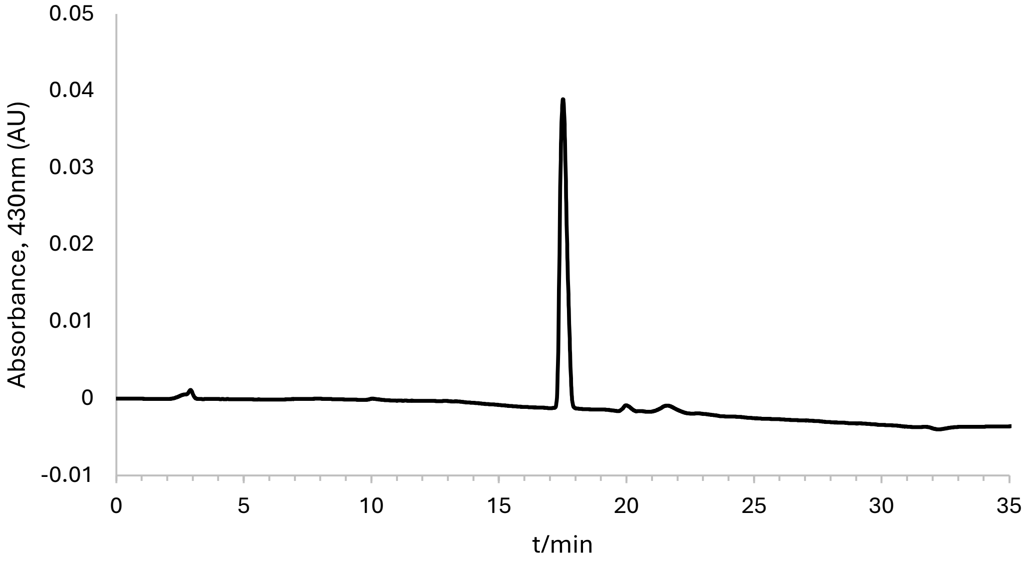


Analytical HPLC chromatogram of **5a**.

**Synthesis of 5b**.

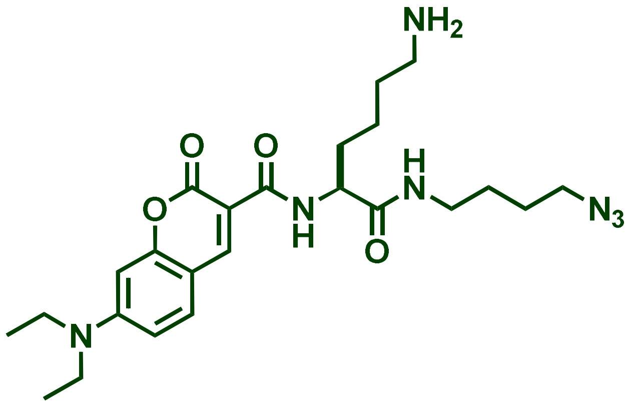


0.2 mol of a-Fmoc-e-$^t$Boc-protected lysine (Me) amino acid was mixed with 1-amino-4-butyl azide (0.22 mmol) in $CH_2Cl_2$ (2.0 mL) at room temperature. To this solution was added EDC (0.25 mmol) and the mixture was stirred at RT followed by LC-MS analysis upon completion of the reaction. The mixture was further diluted with $CH_2Cl_2$ (20 mL) and washed with water, dil. $NaHCO_3$ and brine. The $CH_2Cl_2$ layer was filtered through a $Na_2SO_4$ plug on a funnel and concentrated under reduced pressure to give a solid residue which was carried forward to the next reaction without purification or characterization. The left-over residue was dissolved in methanol (2 mL) and to it, was added LiOH (5M, 0.5 mL), the mixture was stirred at RT overnight and then, the next day, the reaction mixture was analyzed by mass spectrometry to confirm that the starting material was completely consumed to afford the desired Fmoc-deprotected amine. The mixture was carefully acidified with dil. HCl, concentrated under reduced pressure, followed by freezing and lyophilization to yield a solid residue which was used as is for the next reaction. The amine residue was dissolved in $CH_2Cl_2$ and then coumarin (0.18 mmol) mixed with EDC (0.2mmol) was added to the mixture followed by stirring at RT; the progress of the reaction was followed by LC-MS. Upon formation of the product (after 4-6hrs), the reaction mixture was further diluted with $CH_2Cl_2$ (20 mL) and washed with water, dil. $NaHCO_3$ and brine. The $CH_2Cl_2$ layer was filtered through a $Na_2SO_4$ plug on a funnel and concentrated under reduced pressure to give a solid residue which was Boc-deprotected upon treatment with conc. HCl (0.5mL) at RT overnight. The next day, the reaction mixture was diluted with acetonitrile (5mL) and carefully neutralized with $Na_2SO_4$. The solid was removed by filtration through a 0.2 mm filter, and the remaining solution was injected into preparative RP-HPLC for purification on a Phenomenex Luna 10 μm C8(2) 100 Å (250 x 21.2 mm) using a 20 to 100% linear gradient of MeCN in $H_2O$ each containing 0.1% TFA at 10 mL/min. ESI-MS $[M+H]^+$ m/z calcd. 486.2823 for $C_{25}H_{36}N_7O_4$, obsd. 486.2824. $^1$H-NMR (600 MHz, $CD_3CN$) d 1.18 (t, J= 6 Hz, 6H), 1.41-1.58 (m, 4H), 1.67-1.75 (m, 1H), 1.48-1.90 (m, 1H), 2.93-3.02 (m, 2H) 3.19 (q, J= 6Hz, 2H) 3.29 (t, J= 6 Hz, CH), 3.46 (q, J= 6 Hz, 1H, $CH_2$) , 4.45 (d, J= 6 Hz, 1H), 6.50 (d, J= 2 Hz, 1H), 6.74 (dd, J= 2 and 12 Hz, 1H), 6.80 (brs, 1H) 7.06 (brt, J= 12 Hz, 1H), 8.55 (s, 1H, ArH), 9.15 (d, J= 6 Hz, 1H).

$^{13}$C NMR (151 MHz, $CD_3CN$) δ 11.69, 21.95, 25.73, 26.16, 31.97, 38.34, 39.73, 44.74, 50.77, 108.01, 109.18, 110.35, 131.41, 147.91, 153.06, 157.76, 162.51, 171.88.

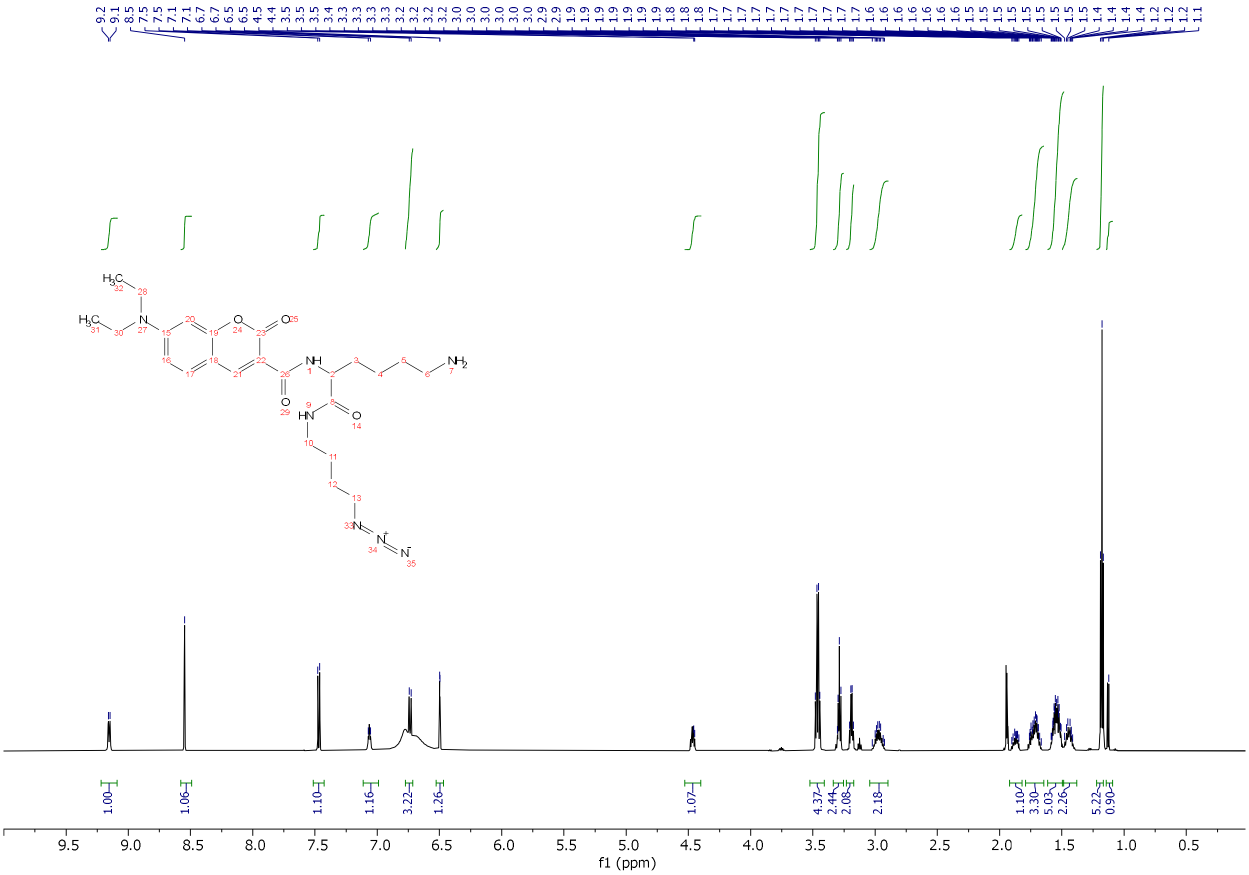

[1]H-NMR of **5b** in $CD_3CN$.

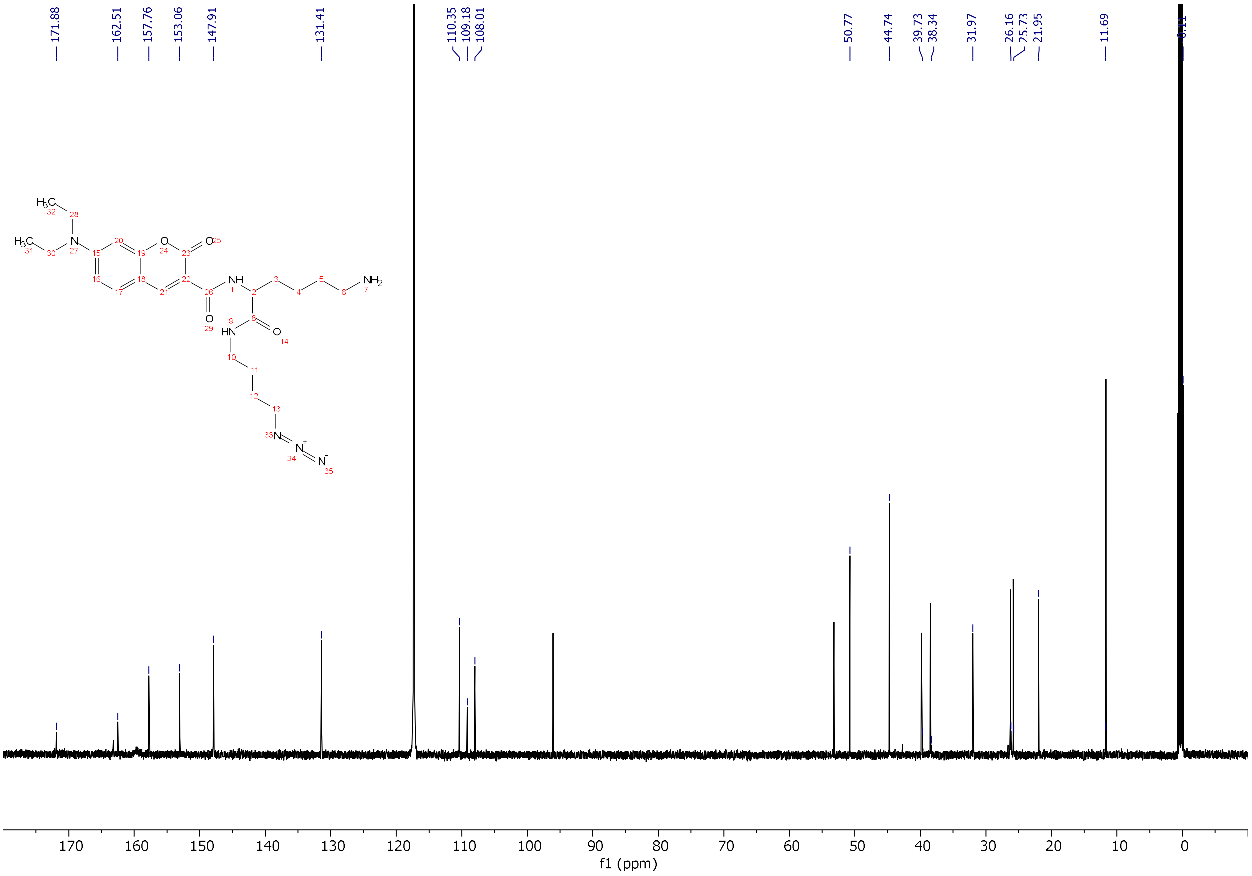


$^{13}$C-NMR of **5b** in $CDCl_3$.

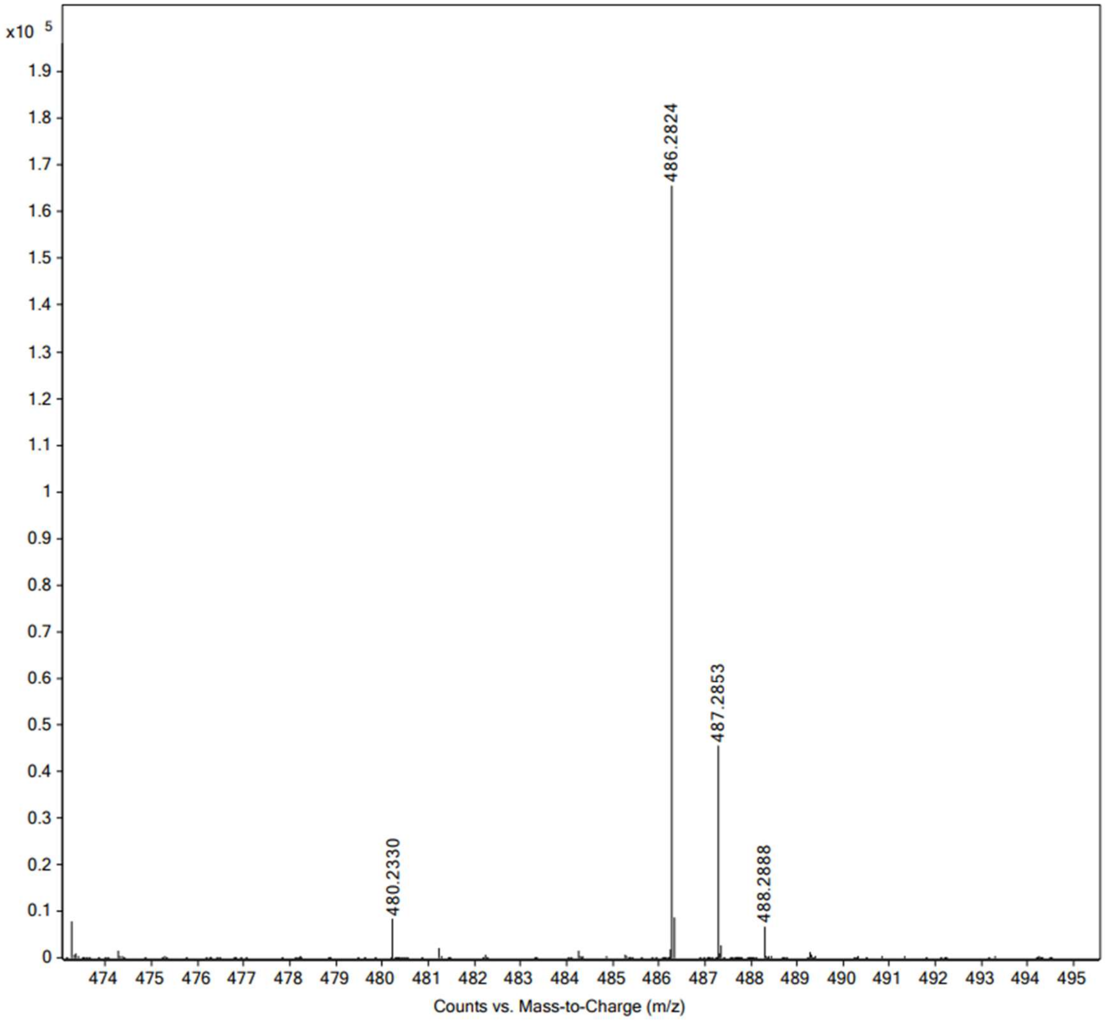


High resolution mass spectrum (ESI) of **5b**.

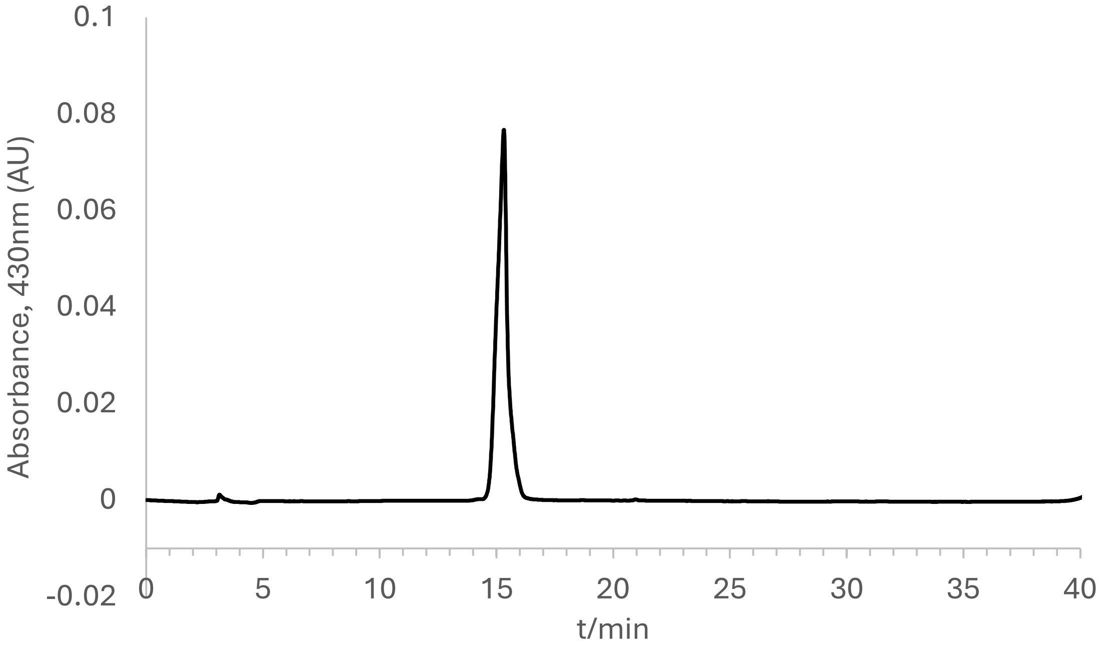


Analytical HPLC chromatogram of **5b**.

**Synthesis of 5c.**

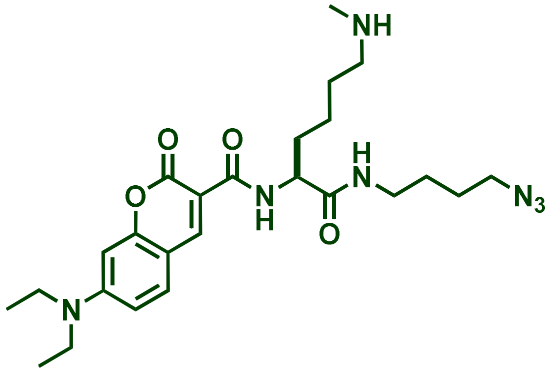


0.2 mmol of a-Fmoc-protected lysine (Me)$_2$ amino acid (96.4mg) was mixed with 1-amino-4-butyl azide (29 mg, 0.25mmol) in $CH_2Cl_2$ (2.0 mL) at room temperature. To this solution was added EDC (0.25 mmol) and the mixture was stirred at RT followed by LC-MS analysis upon completion of the reaction. The mixture was further diluted with $CH_2Cl_2$ (20 mL) and washed with water, dil. $NaHCO_3$ and brine. The $CH_2Cl_2$ layer was filtered through a $Na_2SO_4$ plug on a funnel and concentrated under reduced pressure to give a solid residue which was carried forward to the next reaction without purification or characterization. The left-over residue was dissolved in methanol (2 mL) and to it was added LiOH (5M, 0.5 mL), the mixture was stirred at RT overnight and then, the next day, the reaction mixture was analyzed by mass spectrometry to confirm that the starting material was completely consumed to afford the desired Fmoc-deprotected amine. The mixture was carefully neutralized with dil. HCl (0.1M), concentrated under reduced pressure, followed by freezing and lyophilization to yield a solid residue which was used as is for the next reaction. The amine residue was dissolved in $CH_2Cl_2$ and then coumarin (0.2 mmol) mixed with EDC (0.25mmol) was added to the mixture followed by stirring at RT; the progress of the reaction was followed by LC-MS. Upon formation of the product after 6hrs, the reaction mixture was further diluted with $CH_2Cl_2$ (20 mL) and washed with water, dil. $NaHCO_3$ and brine. The $CH_2Cl_2$ layer was filtered through a $Na_2SO_4$ plug on a funnel and concentrated under reduced pressure to give a residue. This residue was dissolved in 200uL of MeCN and treated concentrated HCl (0.5mL), the mixture was stirred overnight, the LCMS analysis indicated deprotection of Boc group and formation of desired product. The mixture was diluted with 3.0 mL MeCN and concentrated under reduced pressure. Left over residue diluted with $CHCl_3$ (3 mL), carefully neutralized by adding solid $Na_2CO_3$, and solution was filtered through bed of celite, concentrated and purified as follows: The residue was dissolved in 1:1 ($MeCN:H_2O$, ~10mL). The solution was filtered through a 0.2 m filter, and filtrate was injected into preparative RP-HPLC for purification on a Phenomenex Luna 10 µm C8(2) 100 Å (250 x 21.2 mm) using a 20 to 100% linear gradient of MeCN in $H_2O$ each containing 0.1% TFA at 10 mL/min. Homogenous fractions showing absorbance at 220 and 410nm were collected, concentrated under reduced pressure using rotary evaporator to remove as much MeCN as possible, remaining solution was frozen at -80 °C overnight. Next day the frozen solution was lyophilized to obtained dried light orange solid. ESI-MS $[M+H]^+$ m/z 500.2980 calcd. 500.2980 for $C_{25}H_{38}N_7O_4$ obsd. 500.2981. $^1H$-NMR (600 MHz, $CD_3CN$) d 1.38 (t, J= 6 Hz, 6H), 1.41-1.58 (m, 4H), 1.67-1.75 (m, 1H), 1.48-1.90 (m, 1H), 2.94 (brt, 3H), 3.19-3.24 (m, 2H) 3.41 (q, J= 6Hz, 2H) 3.52 (t, J= 6 Hz, CH), 3.68 (q, J= 6 Hz, 4H, $CH_2$) , 4.68 (q, J= 6 Hz, 1H), 6.72 (s,1H), 6.95 (d, J= 6 Hz, 1H) 7.32 (brt, J= 12 Hz, 1H), 7.42 (brs, 1H), 7.69 (d, J= 6 Hz, 1H), 8.13 (brs, 1H), 8.76 (s, 1H), 9.38 (d, J= 6 Hz, 1H).

$^{13}C$ NMR (151 MHz, $CD_3CN$) δ 12.40, 22.81, 25.69, 26.55, 26.99, 32.63, 33.75, 39.28, 45.52, 49.91, 51.50, 54.01, 96.86, 108.78, 109.86, 111.14, 115.59, 132.17, 148.64, 153.79, 158.50, 163.23, 163.97, 172.75.

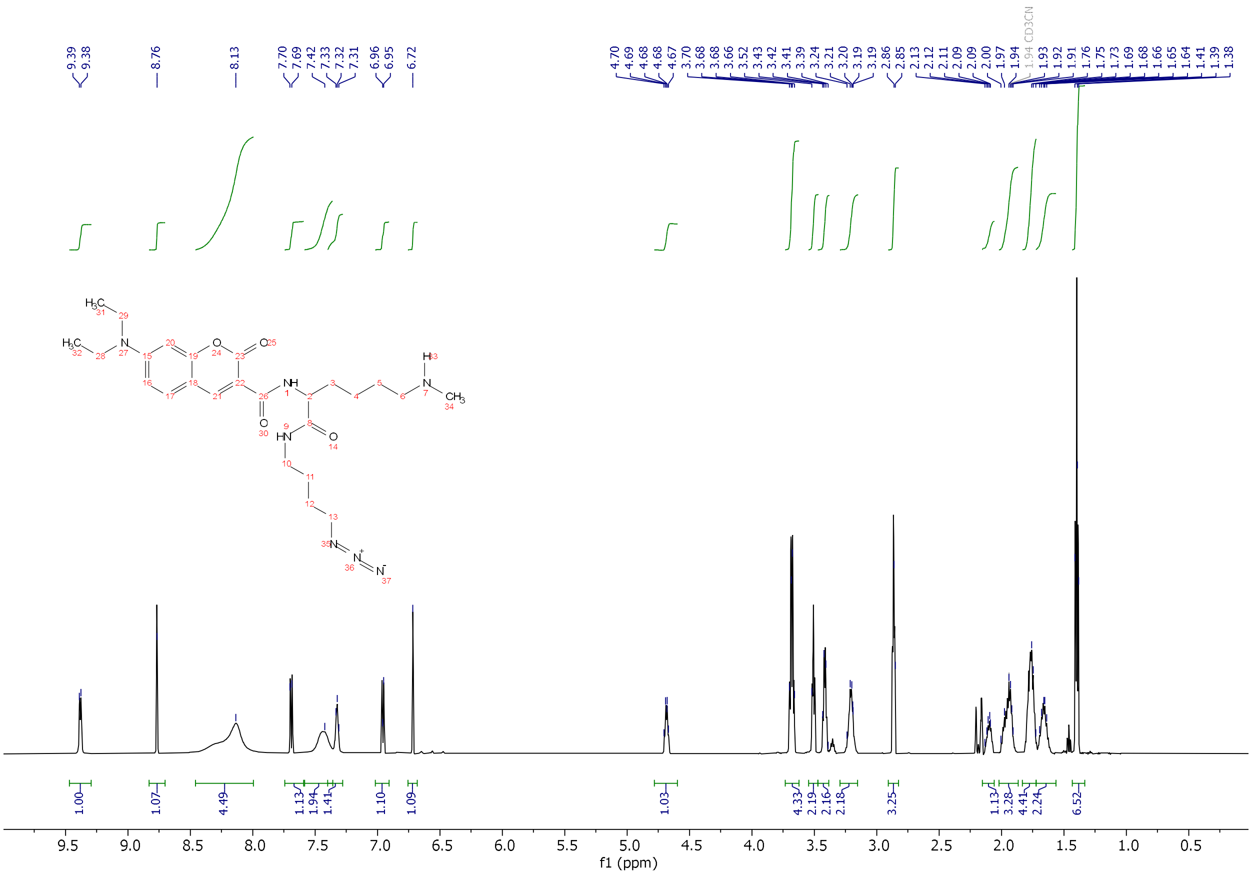

[1]H-NMR of **5c** in $CD_3CN$.

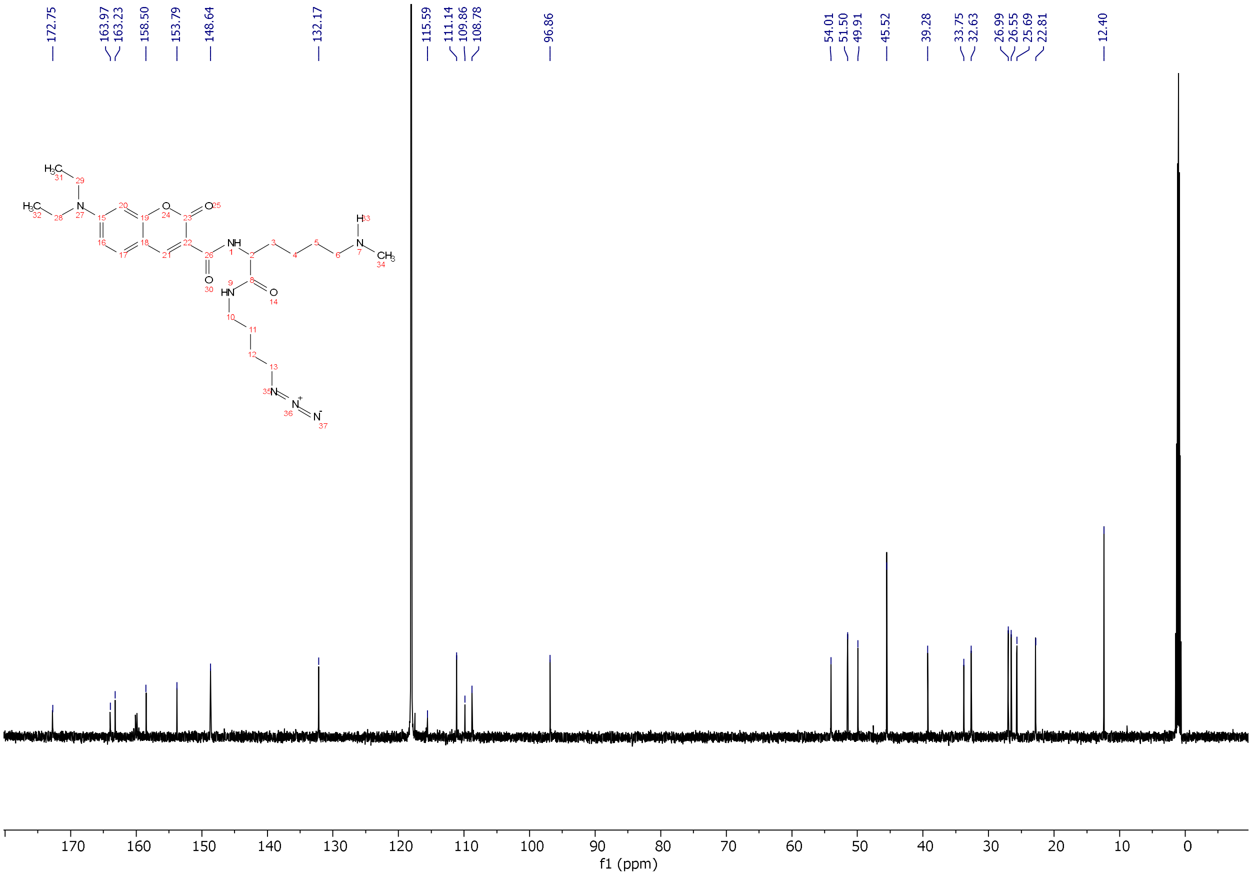


[13]C-NMR of **5c** in $CD_3CN$.

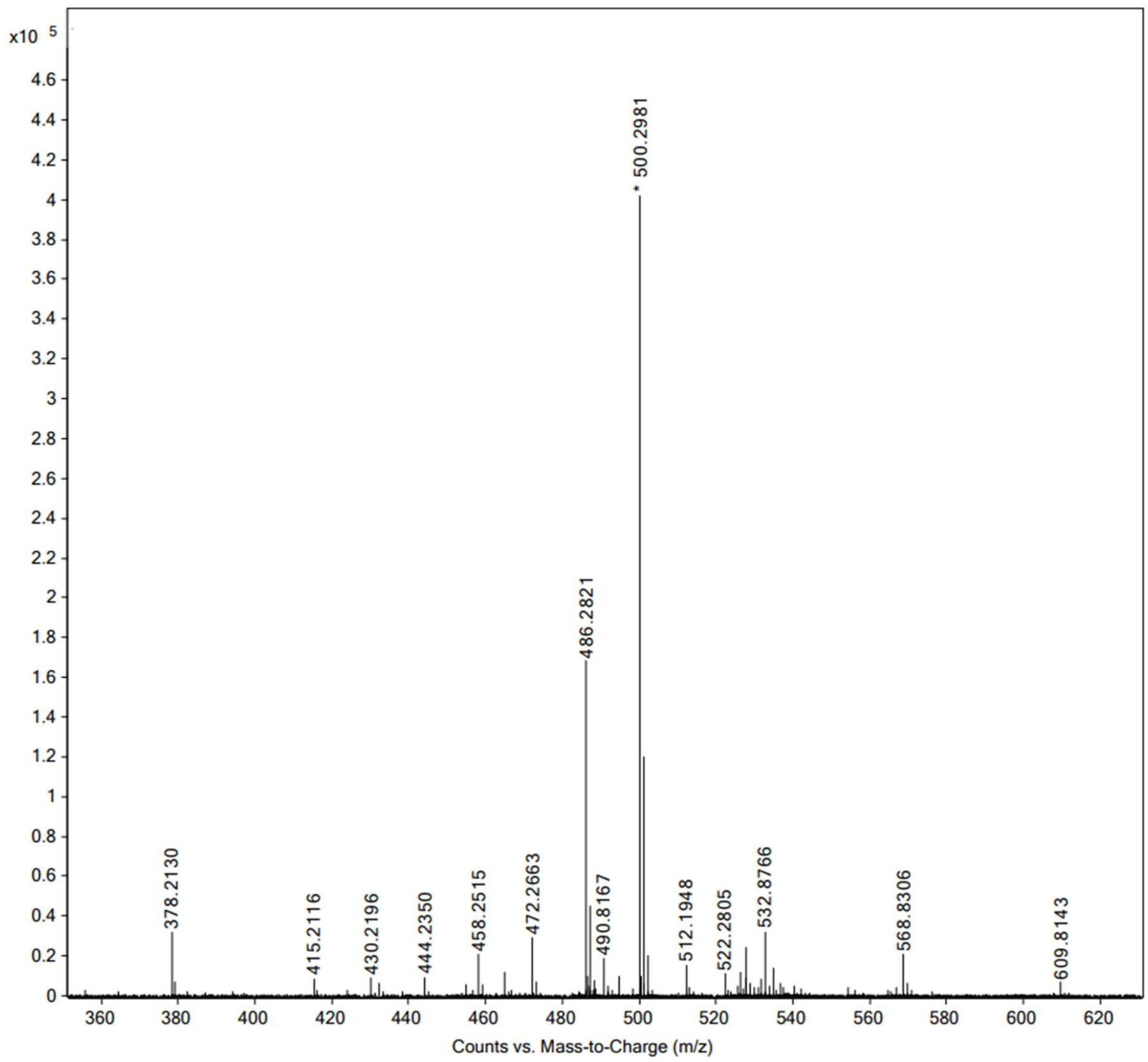


High resolution mass spectrum (ESI) of **5c**.

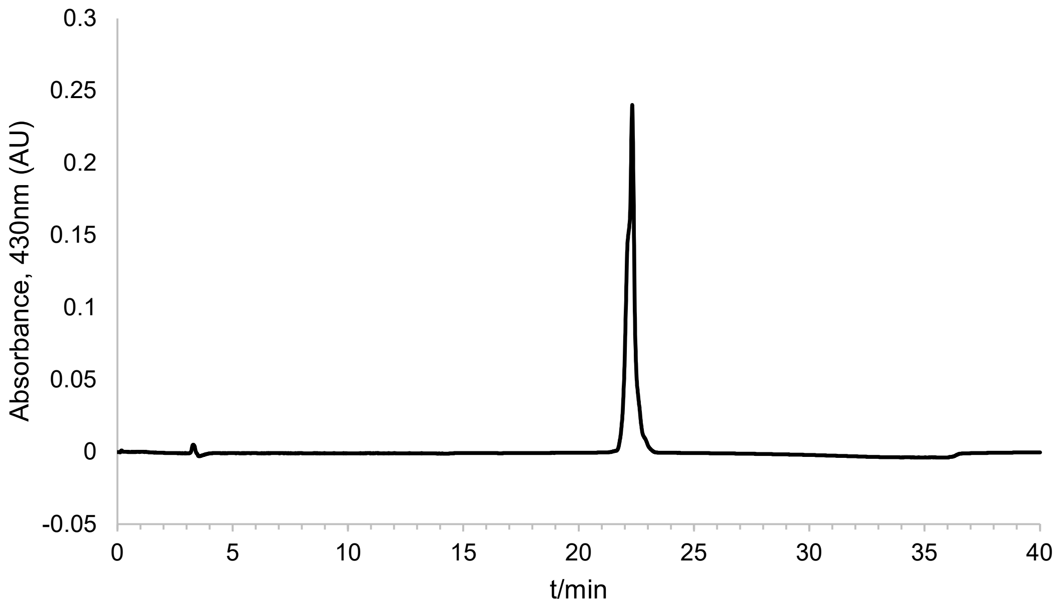


Analytical HPLC chromatogram of **5c**.

**Synthesis of 5d.**

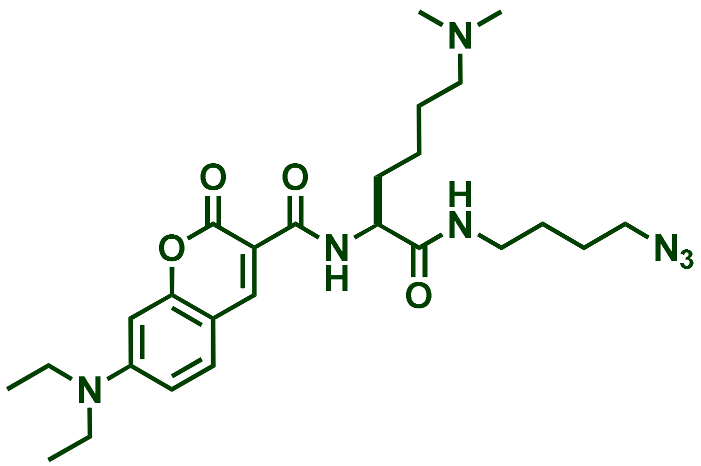


0.2 mmol of a-Fmoc-protected lysine $(Me)_2$ amino acid (79.2mg) was mixed with 1-amino-4-butyl azide (29 mg, 0.25mmol) in $CH_2Cl_2$ (2.0 mL) at room temperature. To this solution was added EDC (0.25 mmol) and the mixture was stirred at RT followed by LC-MS analysis upon completion of the reaction. The mixture was further diluted with $CH_2Cl_2$ (20 mL) and washed with water, dil. $NaHCO_3$ and brine. The $CH_2Cl_2$ layer was filtered through a $Na_2SO_4$ plug on a funnel and concentrated under reduced pressure to give a solid residue which was carried forward to the next reaction without purification or characterization. The left-over residue was dissolved in methanol (2 mL) and to it was added LiOH (5M, 0.5 mL), the mixture was stirred at RT overnight and then, the next day, the reaction mixture was analyzed by mass spectrometry to confirm that the starting material was completely consumed to afford the desired Fmoc-deprotected amine. The mixture was carefully acidified with dil. HCl, concentrated under reduced pressure, followed by freezing and lyophilization to yield a solid residue which was used as is for the next reaction. The amine residue was dissolved in $CH_2Cl_2$ and then coumarin (0.18 mmol) mixed with EDC (0.2mmol) was added to the mixture followed by stirring at RT; the progress of the reaction was followed by LC-MS. Upon formation of the product after 6hrs, the reaction mixture was further diluted with $CH_2Cl_2$ (20 mL) and washed with water, dil. $NaHCO_3$ and brine. The $CH_2Cl_2$ layer was filtered through a $Na_2SO_4$ plug on a funnel and concentrated under reduced pressure to give a residue. The residue was dissolved in 1:1 (MeCN:$H_2O$, 8mL). The solution was filtered through a 0.2 m filter, and filtrate was injected into preparative RP-HPLC for purification on a Phenomenex Luna 10 μm C8(2) 100 Å (250 x 21.2 mm) using a 20 to 100% linear gradient of MeCN in $H_2O$ each containing 0.1% TFA at 10 mL/min. Homogenous fractions showing absorbance at 220 and 410nm were collected, concentrated under reduced pressure using rotary evaporator to remove as much MeCN as possible, remaining solution was frozen at -80 $^{o}$C overnight. Next day the frozen solution was lyophilized to obtained dried light orange solid. ESI-HRMS $[M+H]^+$ m/z calcd. 514.3136 for $C_{26}H_{40}N_7O_4$, obsd. 514.3137. $^1$H-NMR (600 MHz, $CD_3CN$) 1.19 (t, J= 6 Hz, 6H), 1.36-1.41 (m, 2H), 1.51-1.58 (m, 4H), 1.70-1.77 (m, 3H), 1.85-1.90 (m, 1H), 2.78 (brs, 6H), 2.99-3.06 (m, 2H) 3.19 (q, J= 6Hz, 2H) 3.29 (t, J= 6 Hz, 2H), 3.47 (q, J= 6 Hz, 4H, $CH_2$), 4.47 (d, J= 6 Hz, 1H), 6.53 (d, J= 2 Hz, 1H), 6.75 (dd, J= 2 and 12 Hz, 1H), 6.99 (brs, 1H), 7.50 (d, J= 6, 1H), 8.58 (s, 1H, ArH), 8.70 (brs, 1H), 9.11 (d, J= 6 Hz, 1H). $^{13}$C NMR (151 MHz, $CD_3CN$) δ 12.69, 22.64, 23.08, 24.57, 26.84, 27.35, 32.95, 39.42, 43.69, 43.76, 45.73, 51.79, 54.14, 58.54, 97.10, 109.04, 110.43, 110.46, 111.32, 118.33, 132.42, 148.87, 154.04, 158.81, 160.36, 160.61, 163.52, 163.92, 172.62.

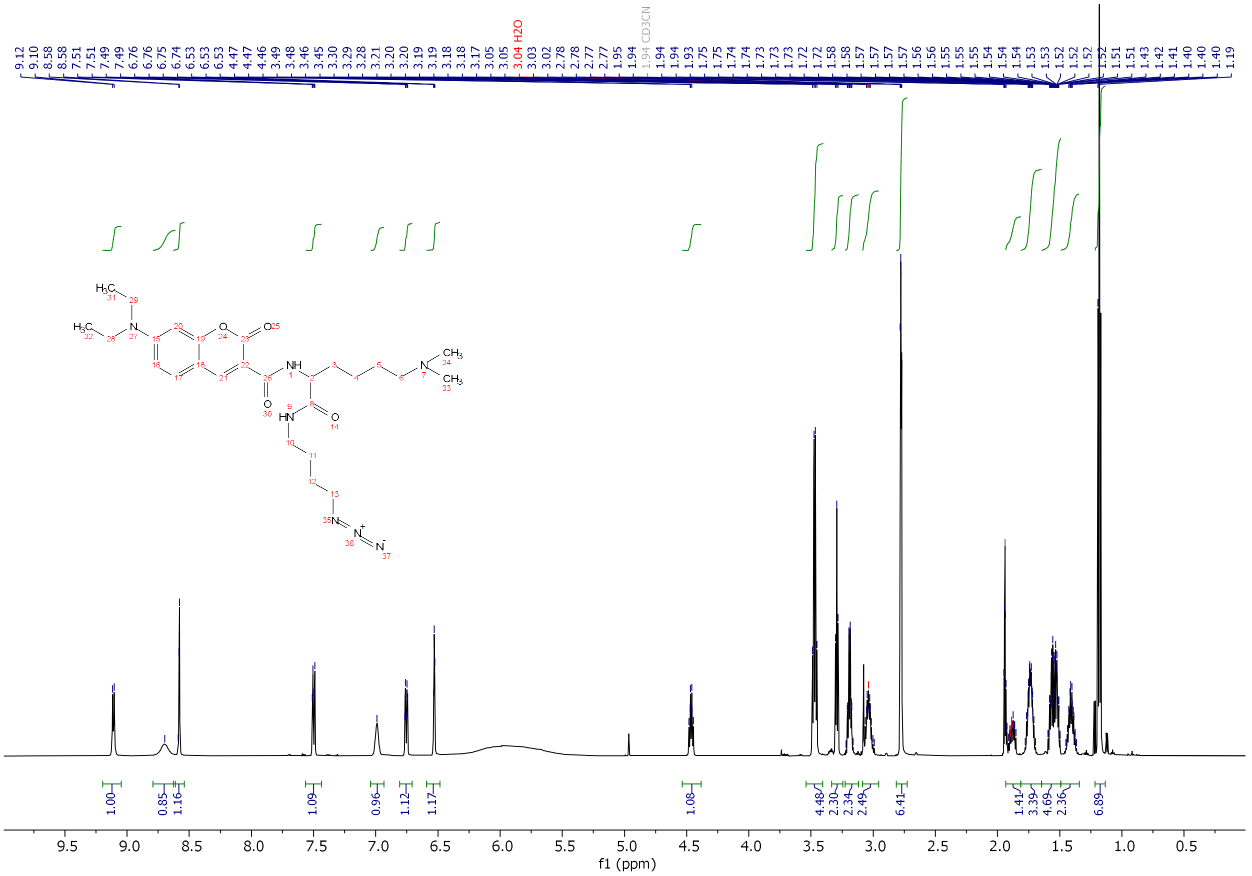

[1]H-NMR of **5d** in $CD_3CN$.

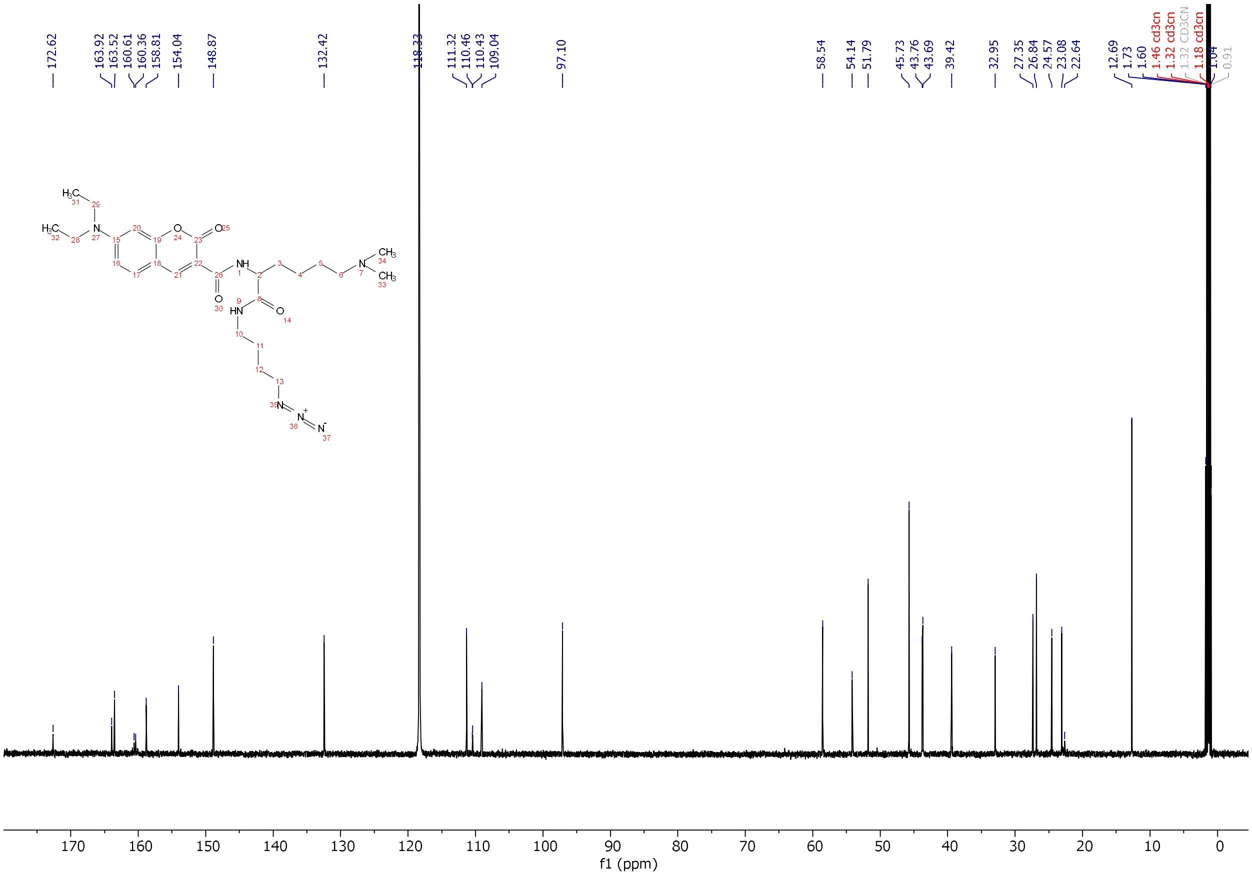

$^{13}$C-NMR of **5d** in $CD_3CN$.

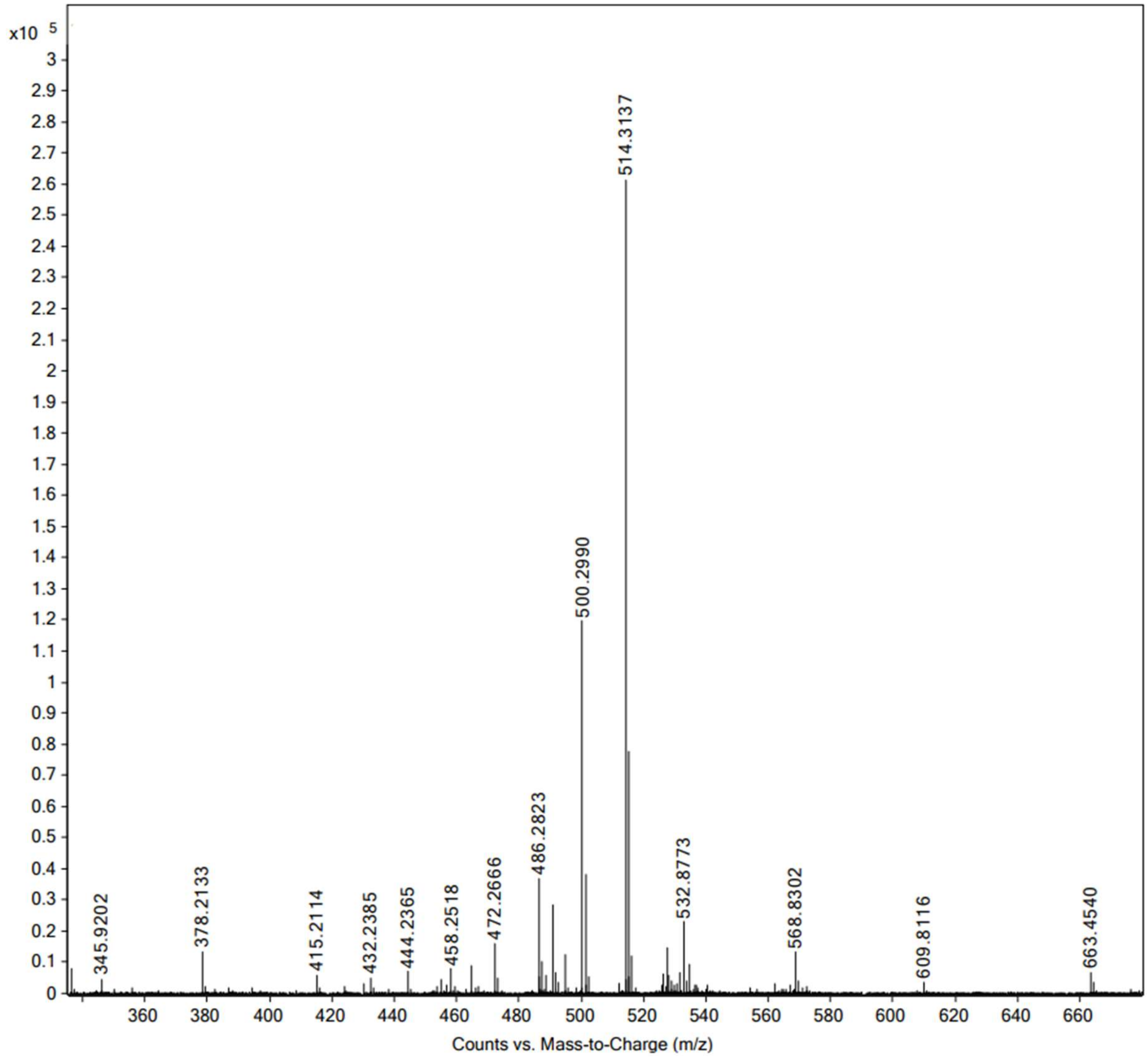


High resolution mass spectrum (ESI) of **5d**.

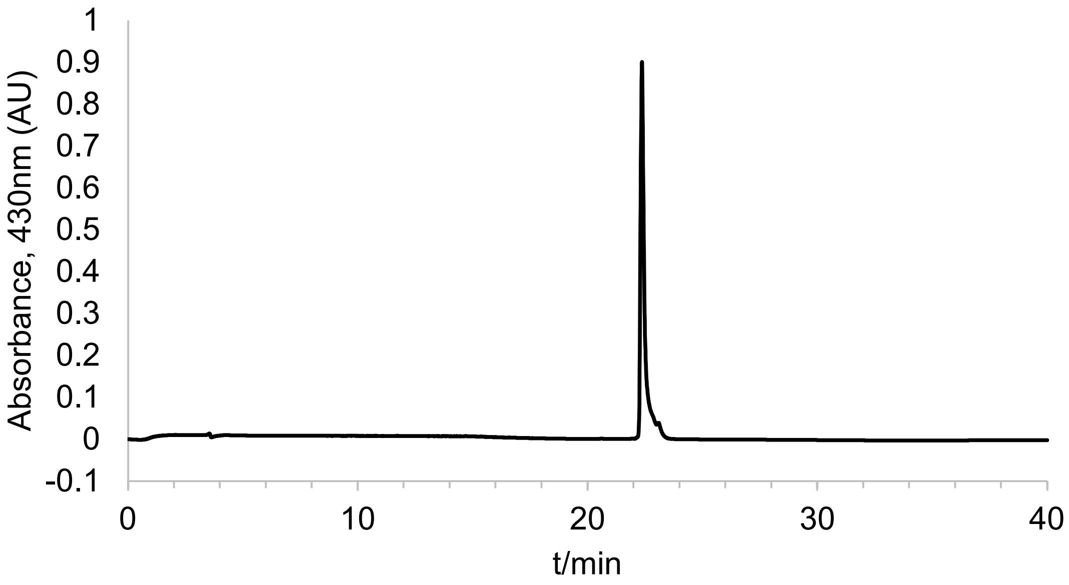


Analytical HPLC chromatogram of **5d**.

**Synthesis of 5e**.

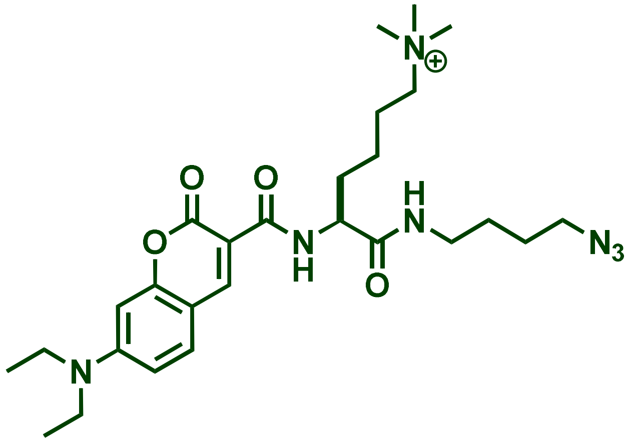

The solution of compound **5d** (25.5mg, 0.05mmol) was dissolved in MeCN (1 mL) and solid $NaHCO_3$ (20.0mg) was added to it and sealed with rubber septum. To this mixture methyl iodide (50 uL) was added via syringe and the mixture was stirred overnight at RT. Next the LCMS analysis indicated starting material was consumed to form the desired quaternary ammonium product. The mixture was concentrated under reduced pressure to give a residue, which was dissolved in minimum amount of $CHCl_3$ (~100uL) and loaded on preparative TLC plate (1000 m, Silica Gel, Analtech), the plate was run with $CHCl_3$:MeOH (95:5)The lowest fluorescent band observed by UV light was removed and eluted with ($CHCl_3$:MeOH; 9:1, 0.5mL x 3). The combined eluent was concentrated under reduced pressure to yield orange oily residue, which analyzed by analytical HPLC and characterized by $^1$H, $^{13}$C-NMR and HRMS. ESI-HRMS $[M]^+$ m/z calcd. 528.3293 for $C_{27}H_{42}N_7O_4$, obsd. 528.3293. $^1$H-NMR (600 MHz, $CD_3CN$) 1.18 (t, J= 6 Hz, 6H), 1.38-1.42 (m, 2H), 1.55-1.61 (m, 4H), 1.76-1.83 (m, 3H), 2.38 (m, 2H), 3.10 (brs, 9H), 3.16-3.23 (m, 2H) 3.19 (q, J= 6Hz, 2H), 3.32 (t, J= 6 Hz, 2H), 3.47 (q, J= 6 Hz, 4H, $CH_2$), 4.61 (m, 1H), 6.54 (brs, 1H), 6.77 (dd, J= 2 and 12 Hz, 1H), 7.40 (brs, 1H), 7.52 (d, J= 6, 1H), 8.08 (s, 0.5H), 8.61 (s, 1H), 9.11 (d, J= 6 Hz, 1H). $^{13}$C NMR (151 MHz, $CD_3CN$) δ 10.43, 11.72, 13.33, 21.91, 22.22, 22.66, 23.72, 25.92, 26.35, 28.69, 30.28, 32.20, 38.28,

38.77, 44.71, 50.81, 53.01, 66.24, 67.37, 96.05, 108.02, 110.29, 129.38, 131.40, 147.91, 152.98, 157.77, 162.49, 162.75, 171.49.

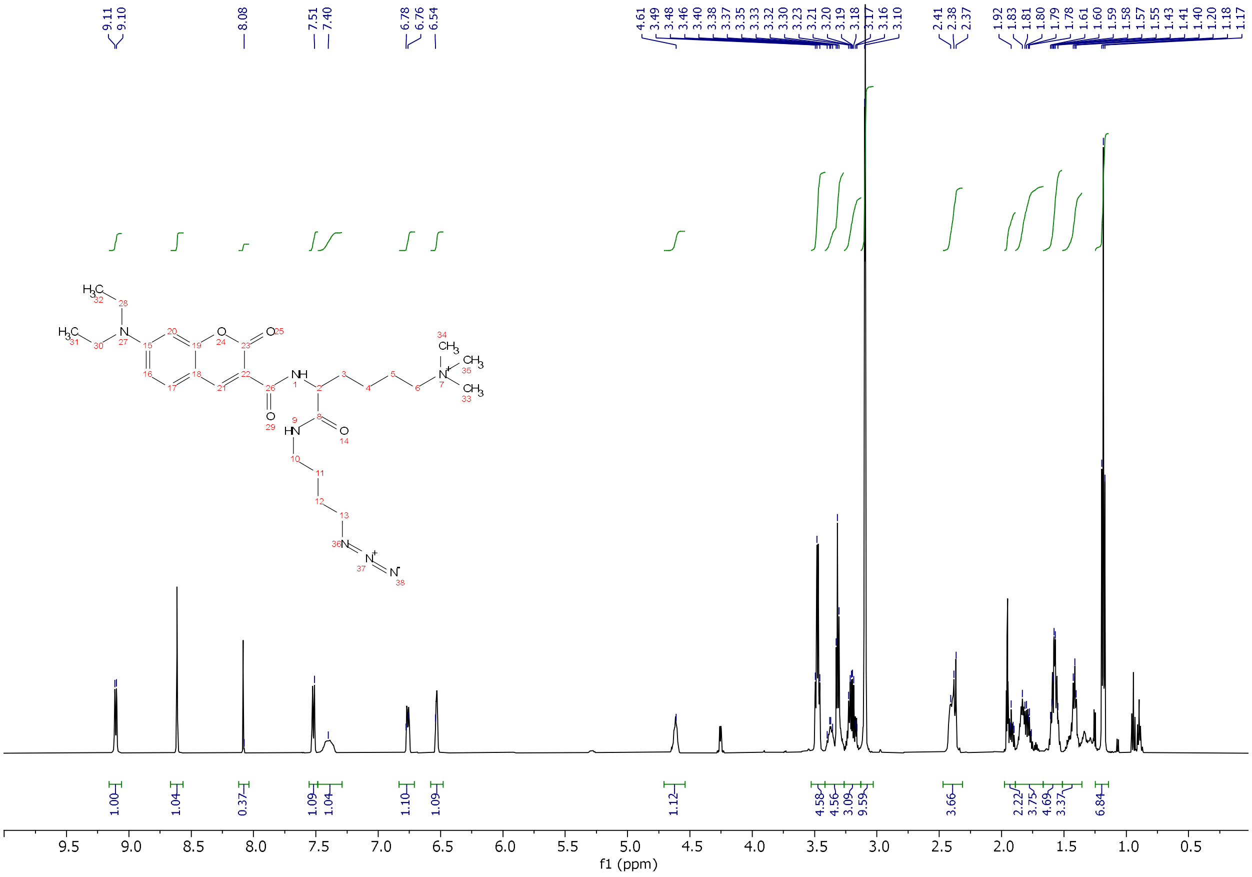

$^1$H-NMR of **5e** in $CD_3CN$.

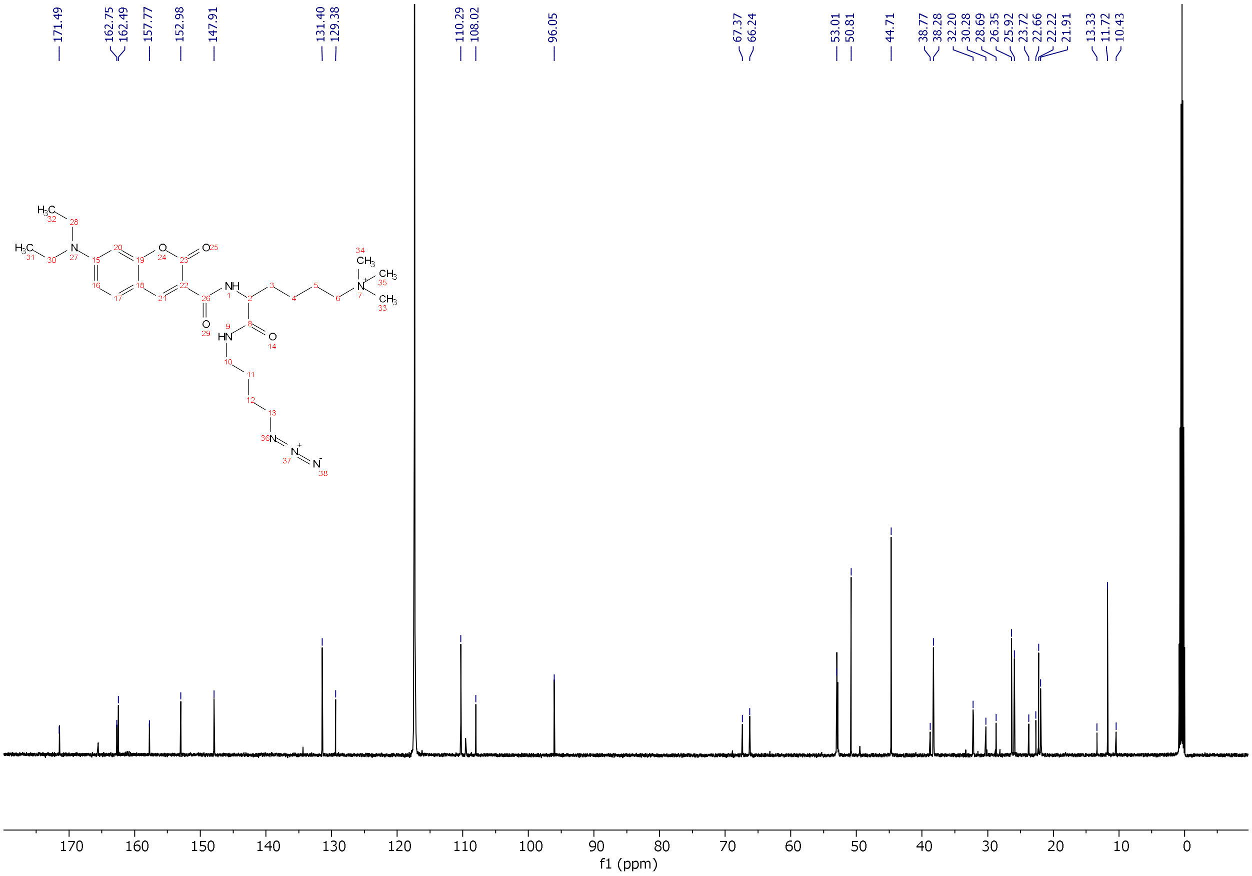


$^{13}$C-NMR of **5e** in $CD_3CN$.

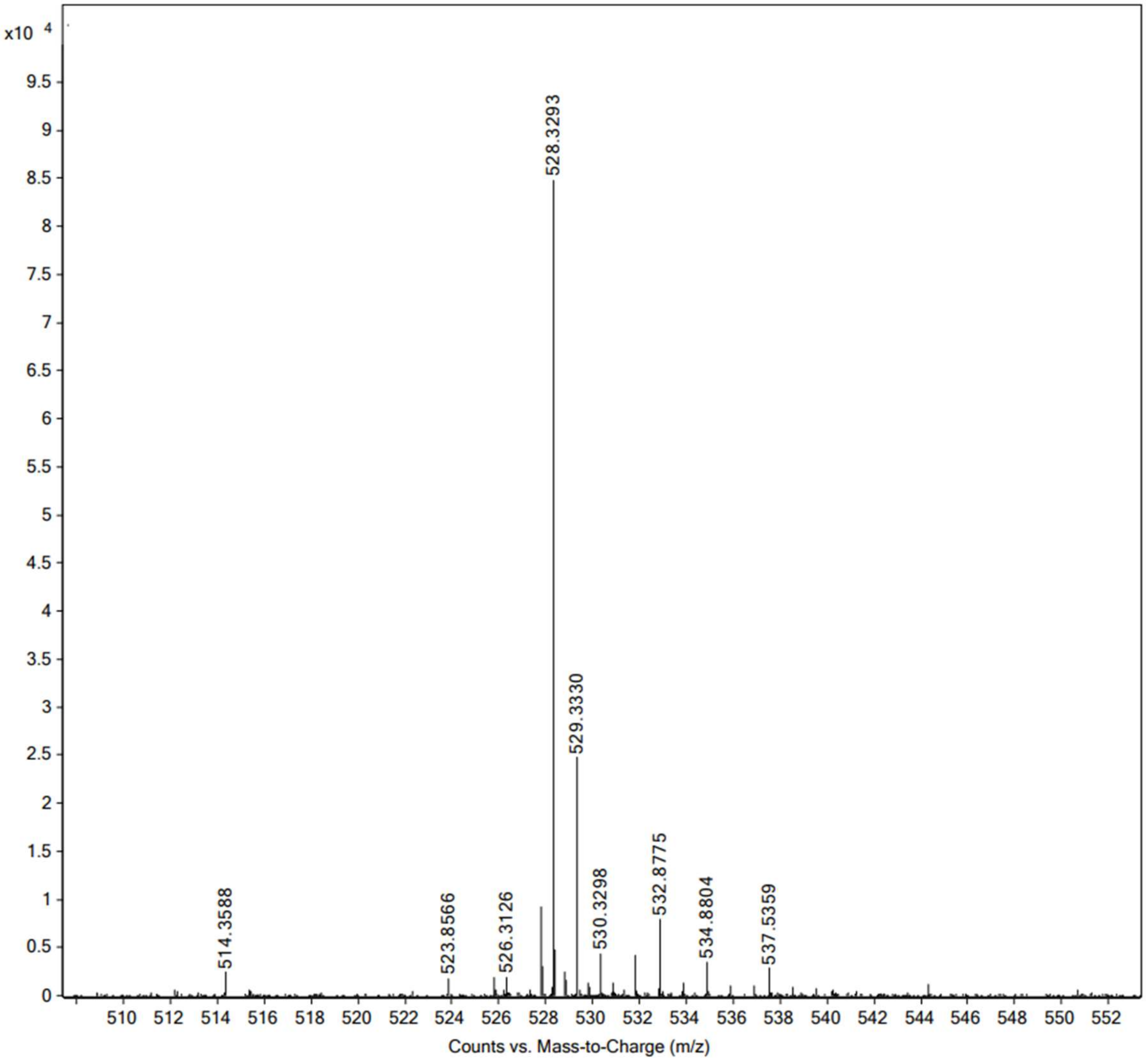


High resolution mass spectrum (ESI) of **5e**.

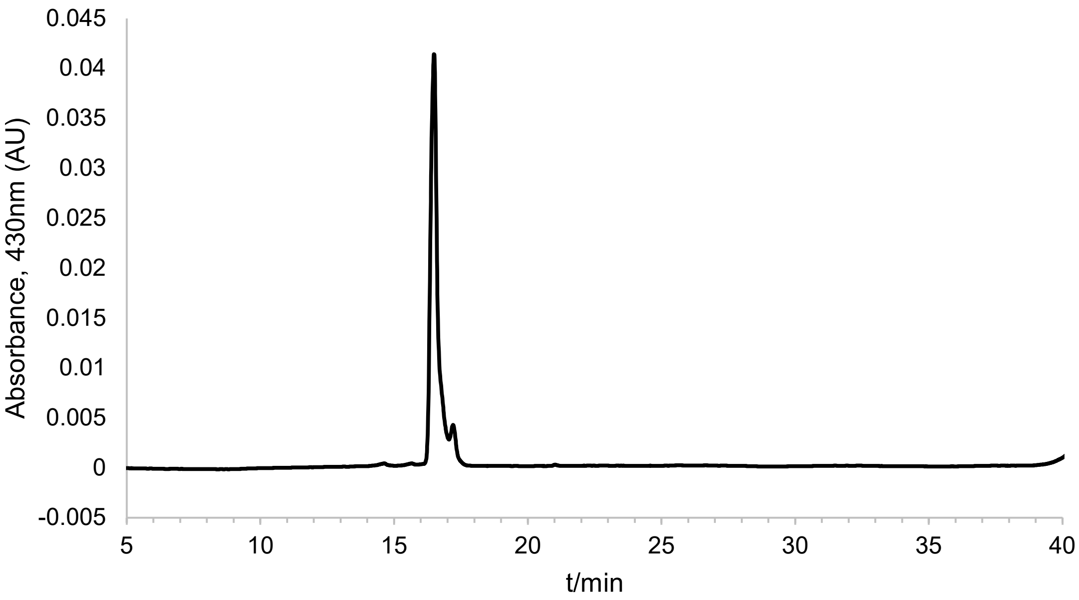


Analytical HPLC chromatogram of **5e**.

**Synthesis of vanAZ1.**

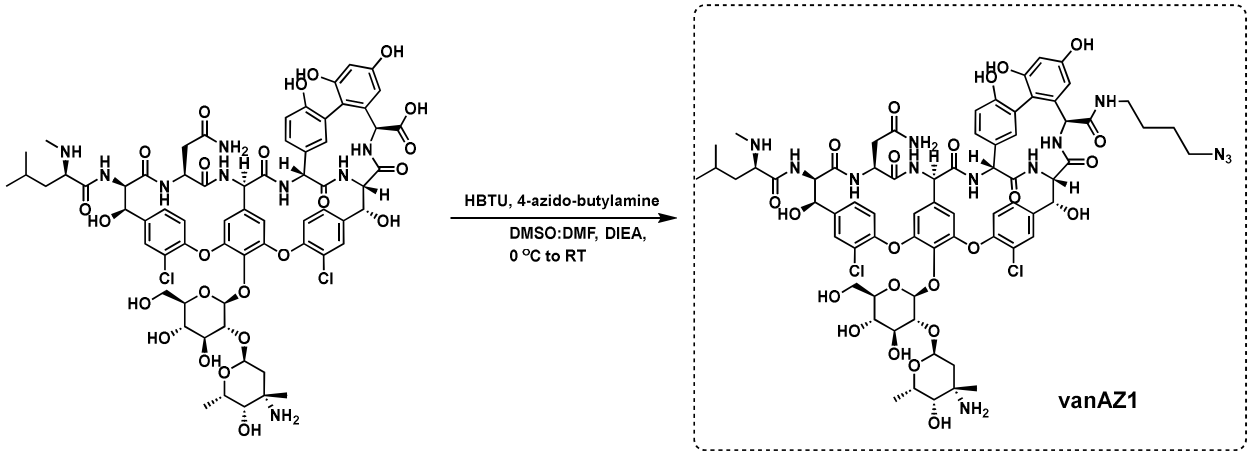


Following reported procedure[1] vancomycin-HCl (MW: 1485, 51.0 mg, 0.0343 mmol, Chem-Impex, Cat # 00315, Lot# 001204-1802003) was dissolved in 1:1 DMSO:DMF, 2 mL and to it was added 4-azido-butylamine (10.0 mL solution in $CH_2Cl_2$ of 1:1 conc, MW: 114.15, 5.0 mg, 0.043 mmol), the solution was cooled in an ice water bath and stirred. While stirring, a solution of HBTU (15.2 mg, 0.04 mmol), followed by DIEA (50 mL) were added. The ice bath was allowed to warm up to room temperature and the mixture was stirred for 6 h. The crude reaction mixture was subjected to analytical HPLC and this analysis showed that the starting material was completely consumed ($R_t$ ~ 12 min) to form a new nonpolar product with Rt ~18 min. The reaction mixture was then diluted with Millipore DD water, 10.0 mL, and the mixture was filtered through 0.4 µ filter. The clear solution was injected to reverse phase preparative HPLC under the same conditions (80:20; water: methanol containing 0.1%TFA) to isolate single peak fractions. The fractions were analyzed by MALDI-TOF prior to combining; fractions showing the desired mass were combined and evaporated under reduced pressure with a rotary evaporator. The leftover solution was frozen in -80 °C freezer and lyophilized for 24 hrs to dry solid (37.5 mg, yield 79%), characterized by $^1$H-NMR and MALDI-TOF mass spectrometry. Calculated m/z 1543.5102 for $C_{70}H_{84}Cl_2N_{13}O_{23}$ $[M+H]^+$; MALDI-TOF observed m/z 1544.985.

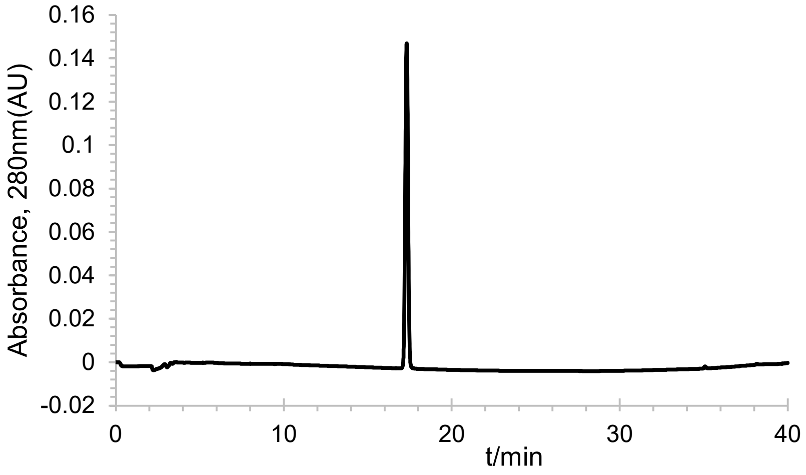


Analytical HPLC chromatogram of **vanAZ1**.

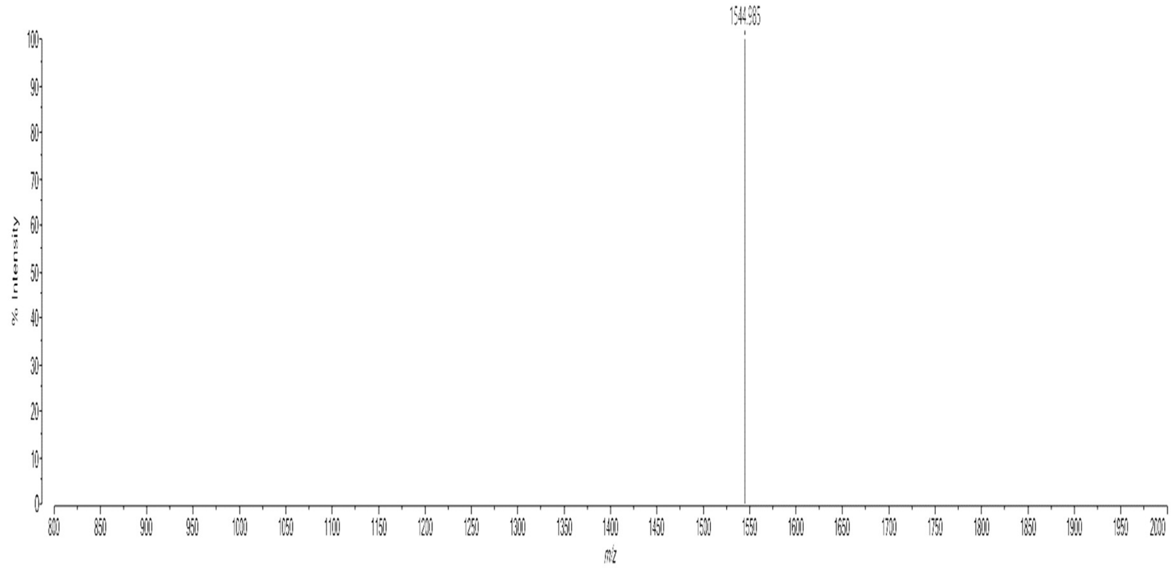


MALDI-TOF spectrum of **vanAZ1**.

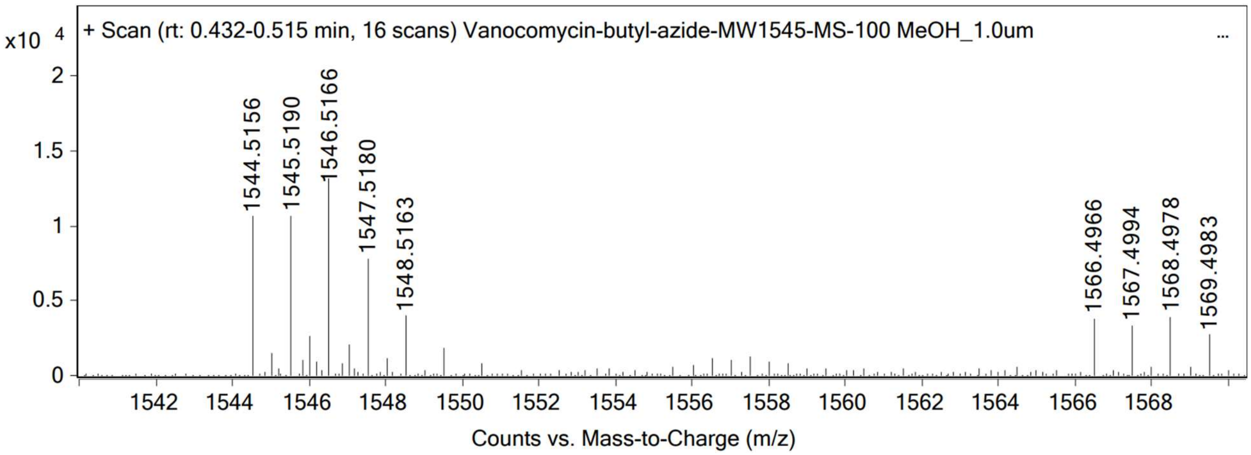


High resolution mass spectrum (ESI) of **vanAZ1**.

**Synthesis of rifAZ1.**

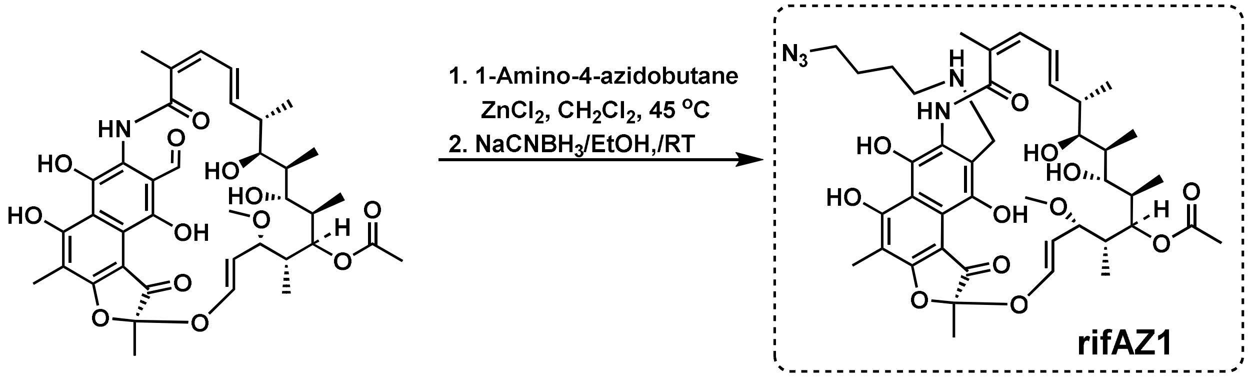


Rifamycin aldehyde (72.5 mg. 0.1 mmol) was dissolved in 5.0mL of anhydrous methylene chloride in 50 mL RB flask equipped with reflux condenser.[2] To this solution was added 1-amino-4-azidobutane (17.1 mg, 0.15 mmol), followed by $ZnCl_2$ (11.0 mg, at room temperature and the mixture was refluxed at 45 °C in an oil bath for 45 minutes. The mixture was then allowed to cool to room temperature. The solvents were evaporated, and the crude material was used as-is for reduction of iminium. The crude material was dissolved in ethanol (200% proof, 2.5 mL), sodium cyanoborohydride (10.7 mg, 0.17 mmol) was added to it and the mixture was stirred at room temperature for 2 h. TLC analysis of the crude material indicated complete conversion of the starting material ($R_t$ ~ 0.4) to a polar new compound (Rt ~0.2) in EtOAc:methanol (95:5) . The volatiles were removed under vacuo using a rotary evaporator. The remaining residue was dissolved in EtOAc and purified by column chromatography over silica gel using gradient ranging from ethyl acetate to 3% methanol EtOAc. Fractions showing homogeneity on TLC were combined and concentrated under reduced pressure to yield a red solid (65.1 mg, 79%). Further purification with reverse phase HPLC (80:20; Water:Methanol) afforded pure a compound that was used for the biological assay. The product was characterized by $^1$H-NMR and MALDI-TOF MS. Mass: m/z 824.4076 for $C_{42}H_{58}N_5O_{12}$ $[M+H]^+$ ; ESI MS observed m/z 824.4329.

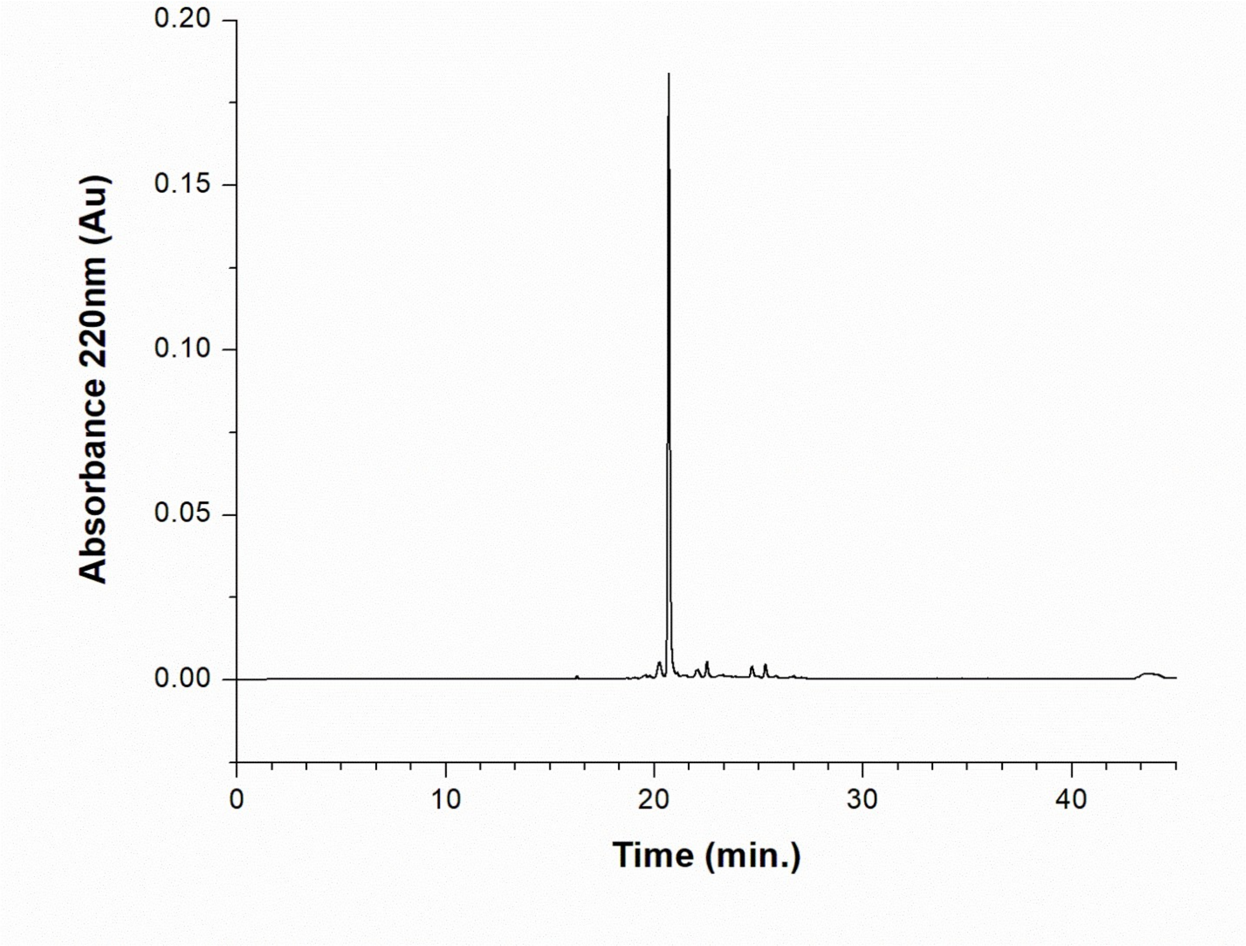


Analytical HPLC chromatogram of **rifAZ1**.

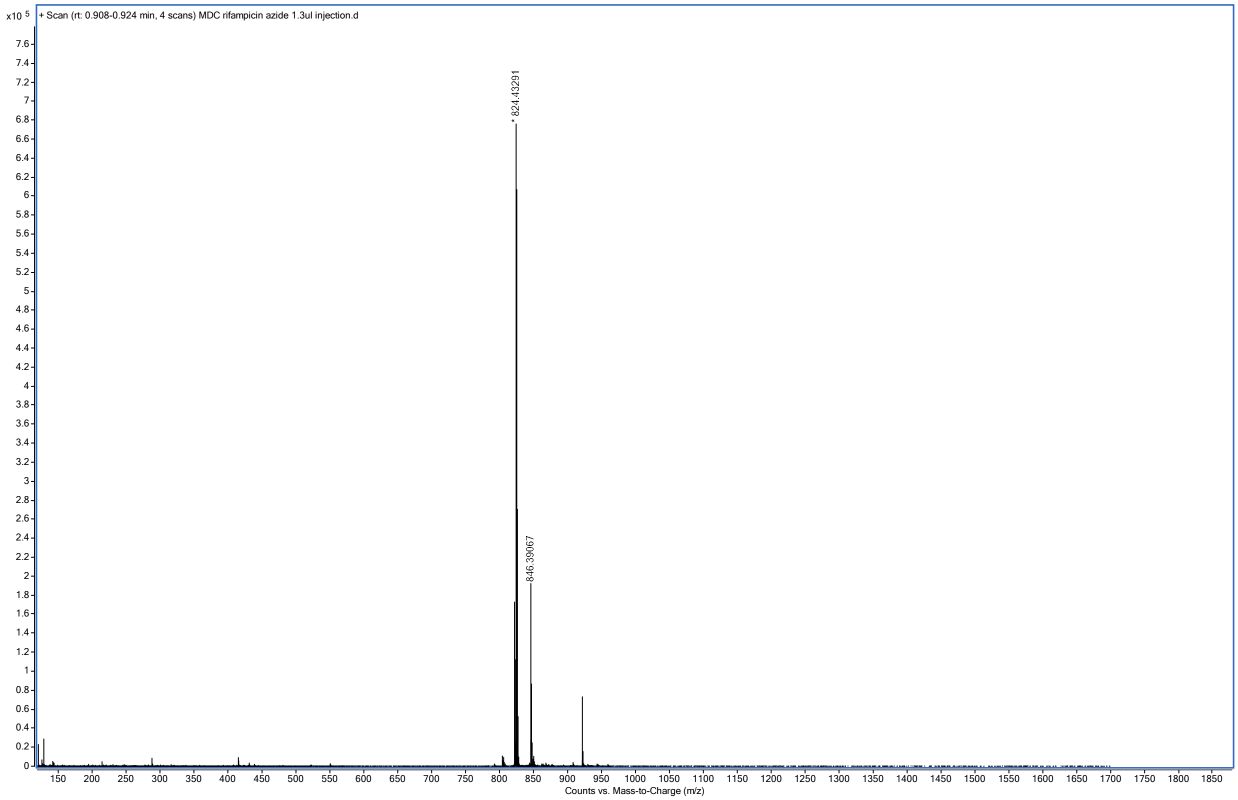


High resolution mass spectrum (ESI) of **rifAZ1**.

**Synthesis of triAZ1.**

Following a published protocol, **triAZ1** was synthesized and characterized as follows:[2]

NMR $^1$H (600 MHz, $CD_3OD$): δ 7.22 (s, 1H), 6.56 (s, 2H), 8.06 (d, J=6 Hz, 1H), 3.99 (t, J=6 Hz, 2H), 3.84 (s, 3H), 3.66 (s, 1H), 3.73 (dd, J= 3, 6 Hz, 1H), 3.58 (t, J= 3, 6 Hz, 1H), 1.91 (m, 2H). $^{13}$C NMR (151 MHz, $CDCl_3$) δ δ 166.2, 156.1, 155.0, 140.3, 137.0, 133.7, 111.0, 107.1, 71.0, 56.6, 33.0, 30.6. MS (ESI): m/z 360.5 for $C_{16}H_{22}N_7O_3$ $[M + H]^+$.

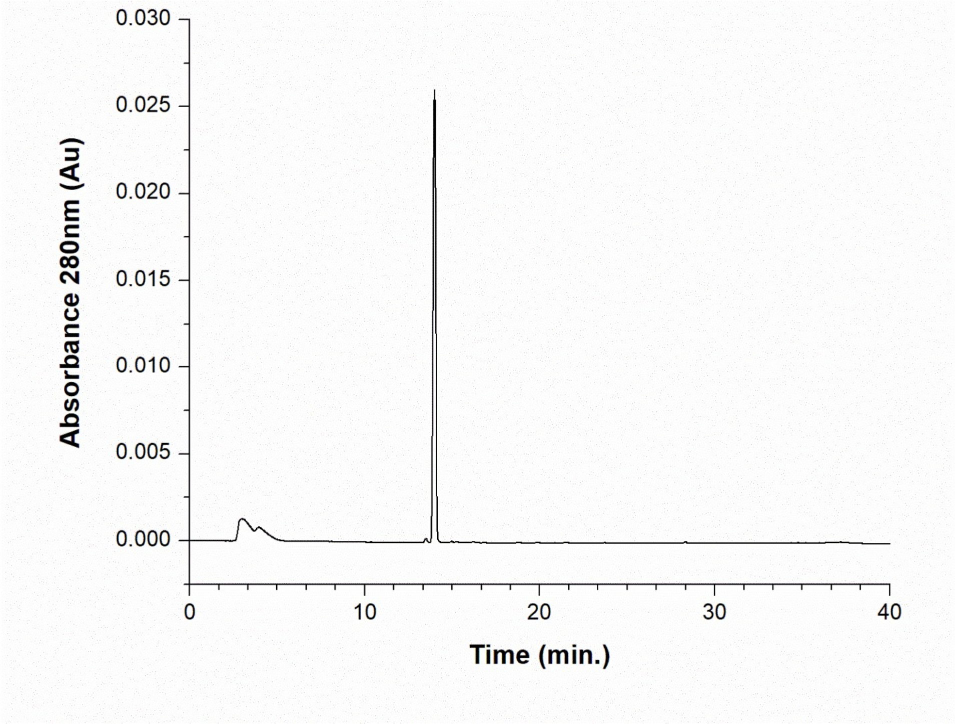


Analytical HPLC chromatogram of **triAZ1**.

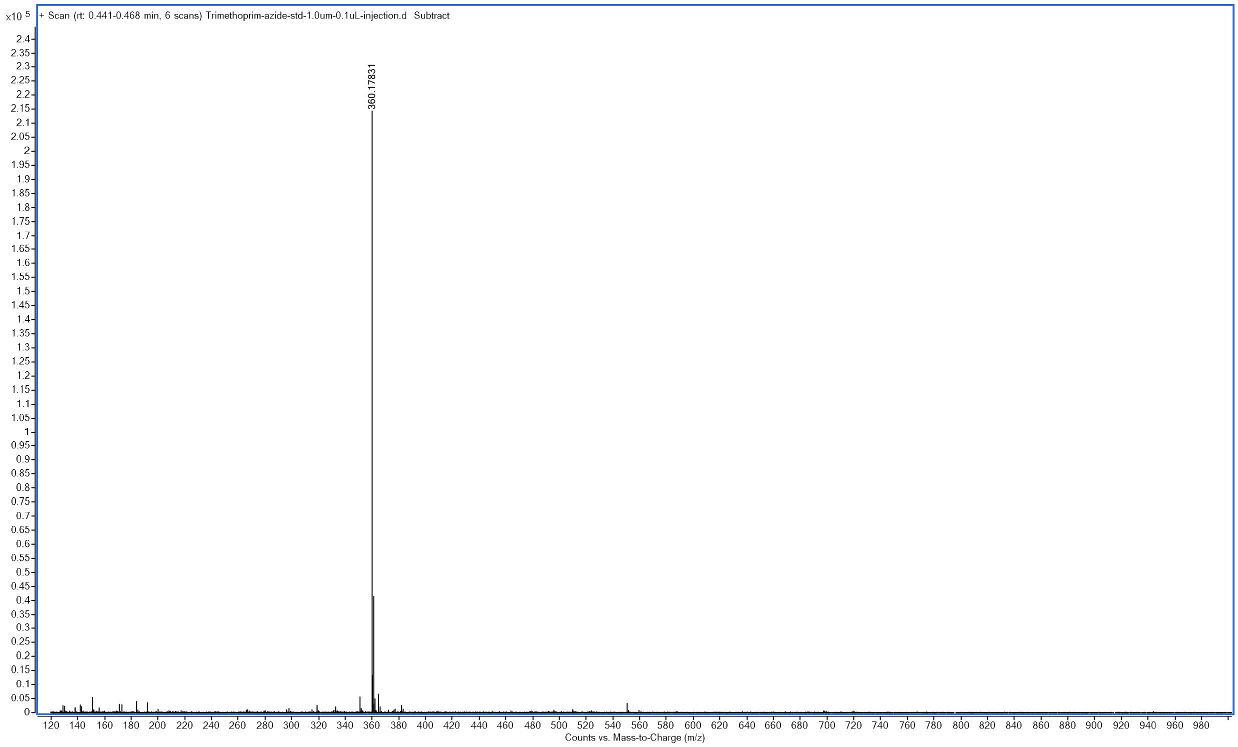


High resolution mass spectrum (ESI) of **triAZ1**.

**Synthesis of moxAZ1**.

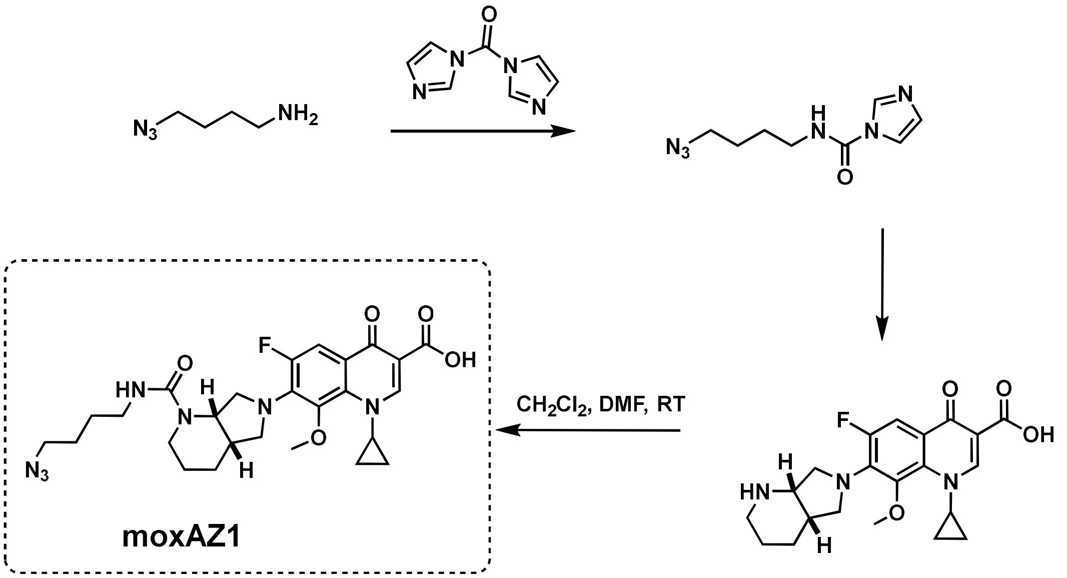


First 4-azidobutyl amine (114 mg, 1 mmol) in anhydrous $CH_2Cl_2$ (3 mL) was mixed with CDI (165.0 mg, 1 mmol). The mixture was stirred at room temperature overnight. The next day, 100 µL of the homogenous solution from this reaction (~0.33 mmol) was added to the clear solution of moxifloxacin (137 mg, 0.33 mol) in DMF and DIEA (50 µL). The mixture was further stirred at room temperature for 6 h. The reaction mixture was analyzed by TLC and ESI-mass for the formation of the product. At the end of the stirring, the reaction mixture was diluted with $CH_2Cl_2$ (20 mL), acidified with citric acid and washed with water (10 mL x 3), and finally with brine. The organic layer was separated and dried over $Na_2SO_4$ and was concentrated under reduced pressure using rotary evaporator. The residue was dissolved in a minimum amount of $CH_2Cl_2$ and loaded on a silica gel column for purification. Using initial $CH_2Cl_2$ and then $CH_2Cl_2$:MeOH (98:2), the desired compound was eluted. The TLC homogenous fractions were combined and concentrated under reduced pressure (109 mg, 61%). The compound was analyzed by analytical HPLC, characterized by high resolution mass spectroscopy and $^1$H-NMR as follows: HRMS observed m/z 542.2527; expected m/z 542.2522 for: $C_{26}H_{32}FN_7O_5$ $[M+H]^+$. $^1$H NMR (600 MHz, $CDCl_3$) δ 0.85 (m, 1H), 1.06-1.15 (m, 2H), 1.29 (m, 1H), 1.54 (m, 1H), 1.62 (m, 1H), 1.64-1.87 (m, 6H), 2.32 (m, 1H), 3.02 (d (m, 1H)t, J=2 and 6 Hz, 1H) 3.26-3.35 (m, 5H),

3.49 (t, J= 6 Hz, 1H), 3.57 (s, 3H, $CH_3$), 3.72 (brd, J= 6 Hz, 1H), 3.83 (dt, J= 1 and 6Hz, 1H), 3.97 (m, 1H), 4.10 (m, 1H), 4.80 (q, J= 6Hz, 1H), 7.74 (d, J=12 Hz, 1H), 8.76 (s, 1H). $^{13}C$ NMR (151 MHz, $cdcl_3$) δ 8.74, 10.64, 24.16, 25.04, 26.44, 27.63, 35.78, 40.59, 40.61, 48.75, 51.30, 52.58, 56.46, 107.73, 107.96, 108.12, 118.81, 134.52, 137.36, 137.43, 141.08, 149.80, 152.97, 154.63, 157.95, 167.17, 176.79, 187.14.

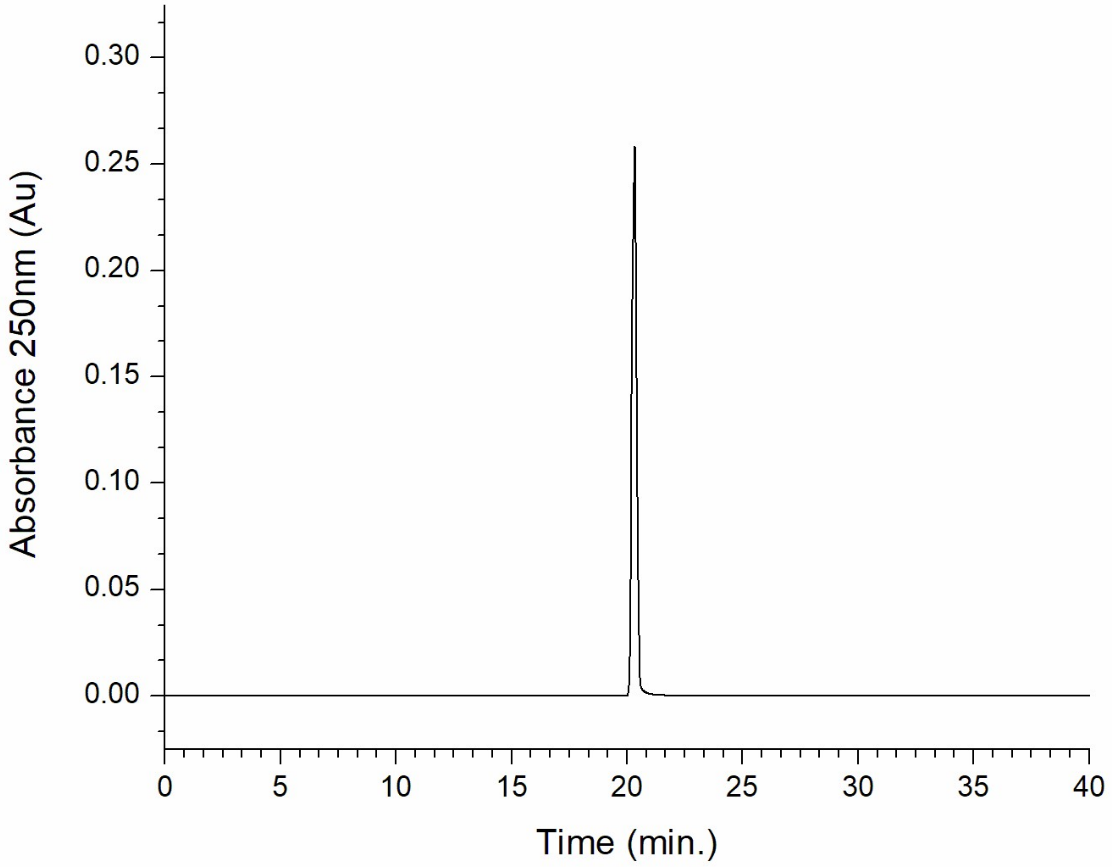


Analytical HPLC chromatogram of **moxAZ1**.

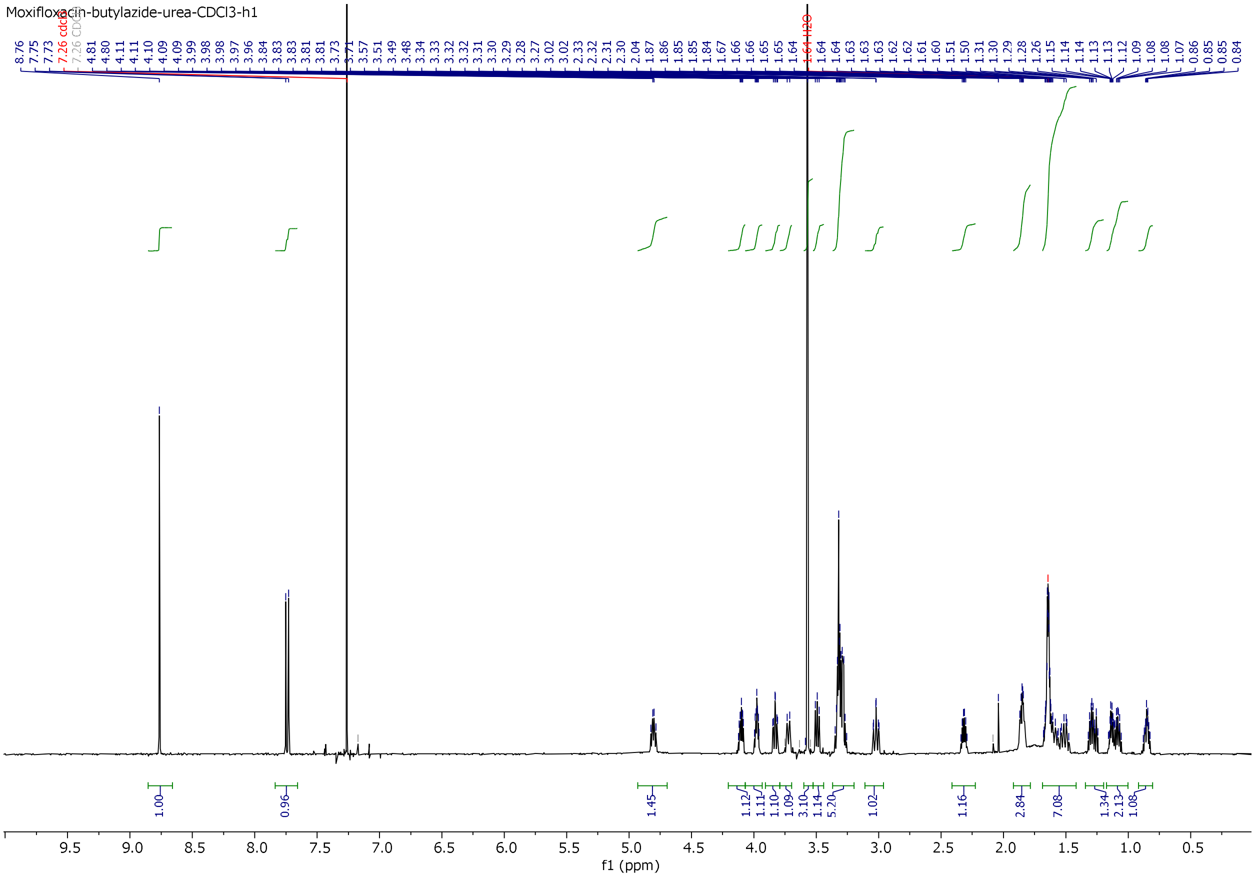

[1]H-NMR of **moxAZ1**.

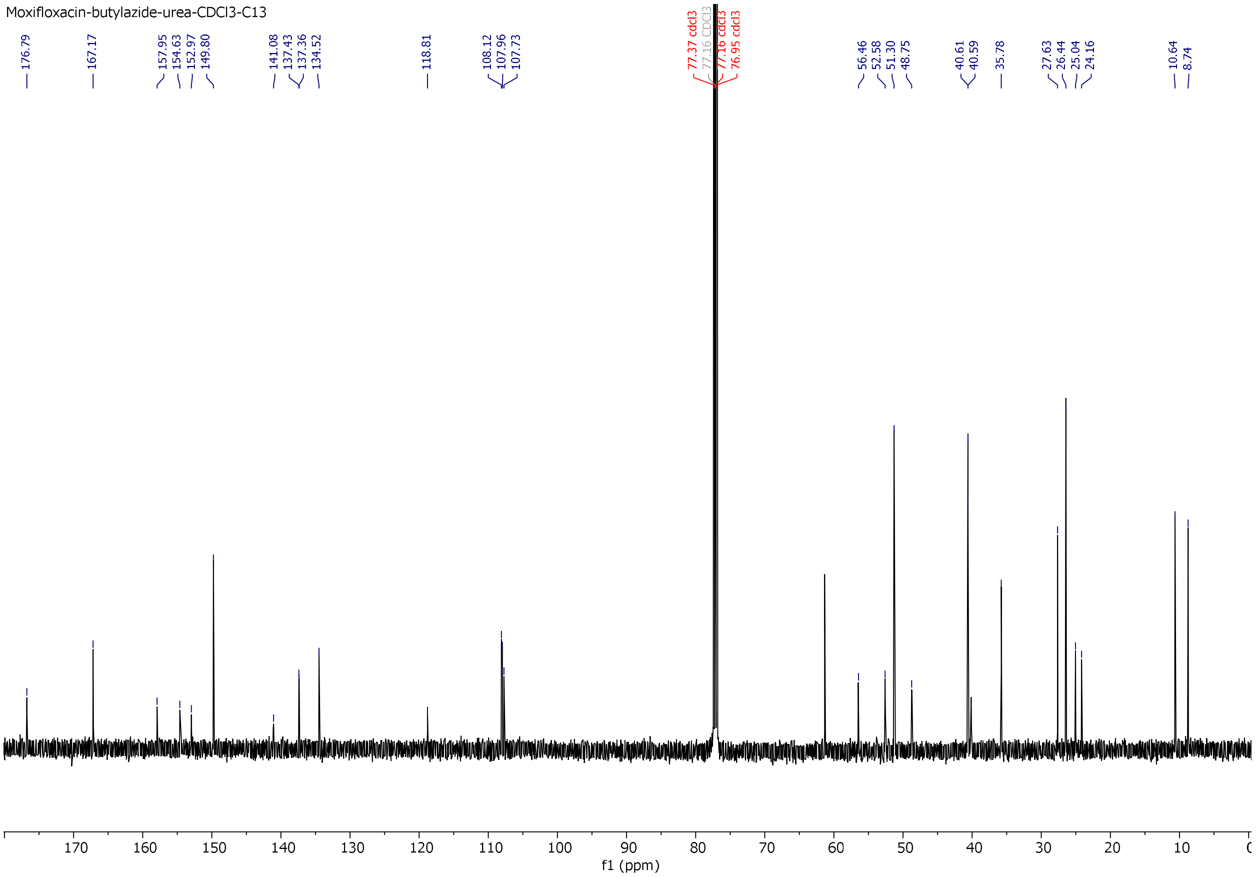


$^{13}C$-NMR of **moxAZ1**.

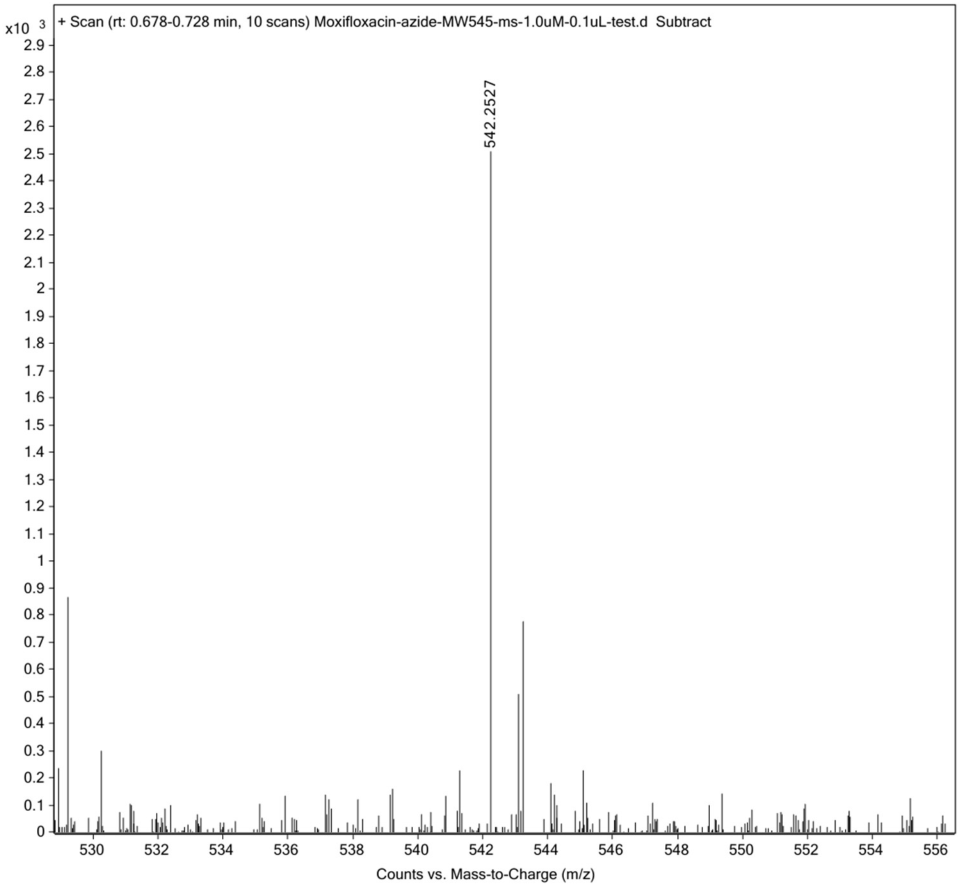


High resolution mass spectrum (ESI) of **moxAZ1**.

**Synthesis of novAZ1**.

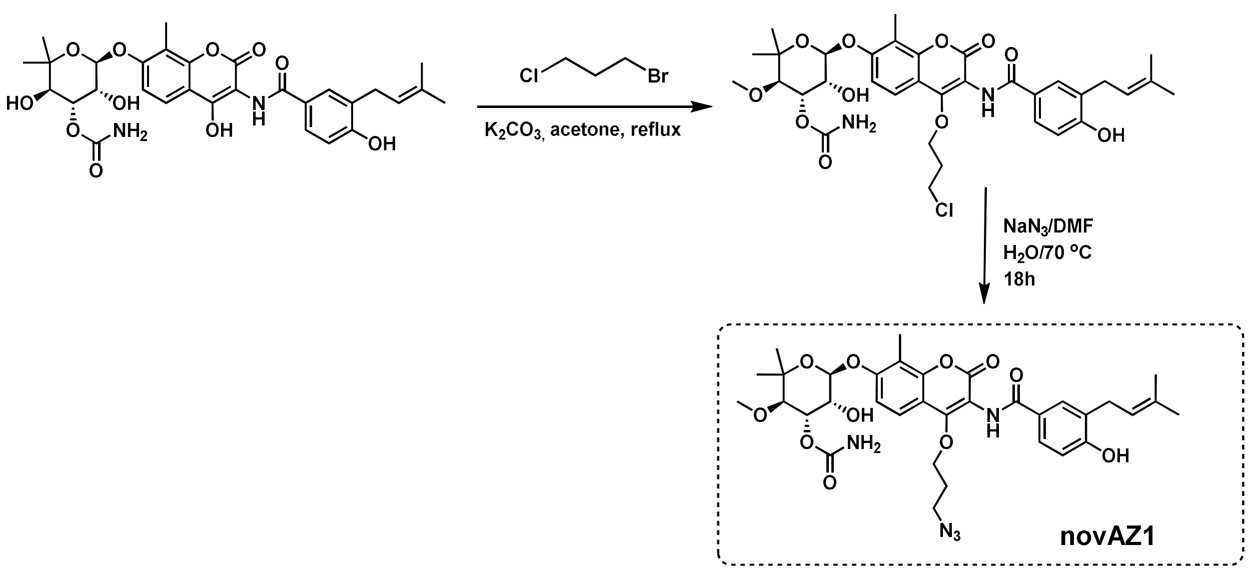

Novobiocin (122.4 mg, 0.2mmol, AK Scientific, Cat. # X5037) was suspended in acetone (10.0 mL) and to this solution, was added $K_2CO_3$ (500 mg) and 1-bromo-3-chloropropane (40 mg, 0.258 mmol). The mixture was heated to reflux with stirring for 4 hours during which time the reaction was monitored by TLC analysis to confirm the formation of the non-polar compound. The reaction mixture was then allowed to cool to room temperature and most of the acetone was removed under reduced pressure to afford a leftover residue that was dissolved in water and ethyl acetate (1:2 ratio, 30 mL total). The mixture was carefully acidified to pH 6.0 with dil. HCl (0.1M) following which the ethyl acetate layer was separated, washed with brine, and concentrated under reduced pressure. The crude material thus obtained was purified with silica gel column chromatography, and the compound was eluted with chloroform: methanol (9:1). The TLC-analyzed homogenous fractions were combined and concentrated under reduced pressure. The oily residue was triturated with ether to yield a white solid (95 mg, 69% yield). This chloropropyl ether was characterized by $^1$H-NMR and mass spectroscopy. MS (ESI): m/z 710.8 for $[M+Na]^+$.

The chloropropyl ether-novobiocin (69.0mg, 0.1 mmol) was dissolved in DMF:Water (9:1, 3mL). $NaN_3$ (26 mg, 4 mmol, 4.0eq.) was added to the solution and the mixture was heated at 70 °C for 16 hrs. The next day, TLC analysis showed a slightly more non-polar spot formed and mass analysis of the reaction mixture revealed complete conversion to azide. The reaction mixture was cooled to RT, diluted with ethyl acetate and water (1:1, 20 mL) and extracted. The organic layer was washed with water, brine and dried over $Na_2SO_4$. Concentration under reduced pressure afforded the crude material, which was purified with silica gel column chromatography using chloroform:methanol (98:2), to yield the propyl-azide derivative of novobiocin as an off white solid (42 mg, 60%). NMR $^1H$ (600 MHz, $CD_3OD$): δ 7.77 (m, 2H, ArH), 7.71 (d, J = 12Hz, 1H, ArH), 7.25 (d, J = 12Hz, 1H, ArH), 6.86 (d, J = 6.0 Hz, 1H, ArH), 5.60 (s, 1H, OCHO), 5.33 (m, 2H), 4.53 (t, J = 6.0 Hz, 2H, O-$CH_2$) 4.25 (brs, 1H), 3.59 (d, J = 6 Hz, 1H), 3.56 (s, 3H, $OCH_3$), 3.51 (t, J = 6 Hz, 1H), 3.36 (d, J = 12 Hz, 1H), 2.32 (s, 3H, $ArCH_3$), 2.04 (m, 2H, $CH_2$), 1.75 (d, J = 6Hz, 6H, $CH_3$ x 2), 1.35 (s, 3H, $CH_3$),1.15 (s, 3H, $CH_3$). MS (ESI): m/z 696.2 and 718.2 and for $C_{16}H_{22}N_7O_3$ $[M + H]^+$ and $[M + Na]^+$ respectively ; observed 718.2701.

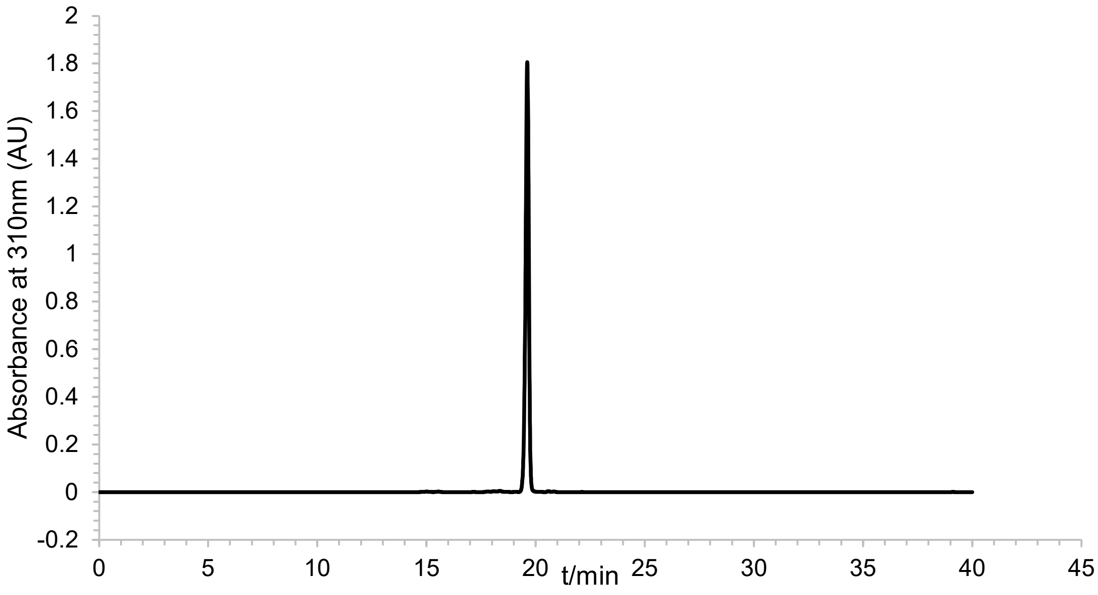


Analytical HPLC chromatogram of **novAZ1**.

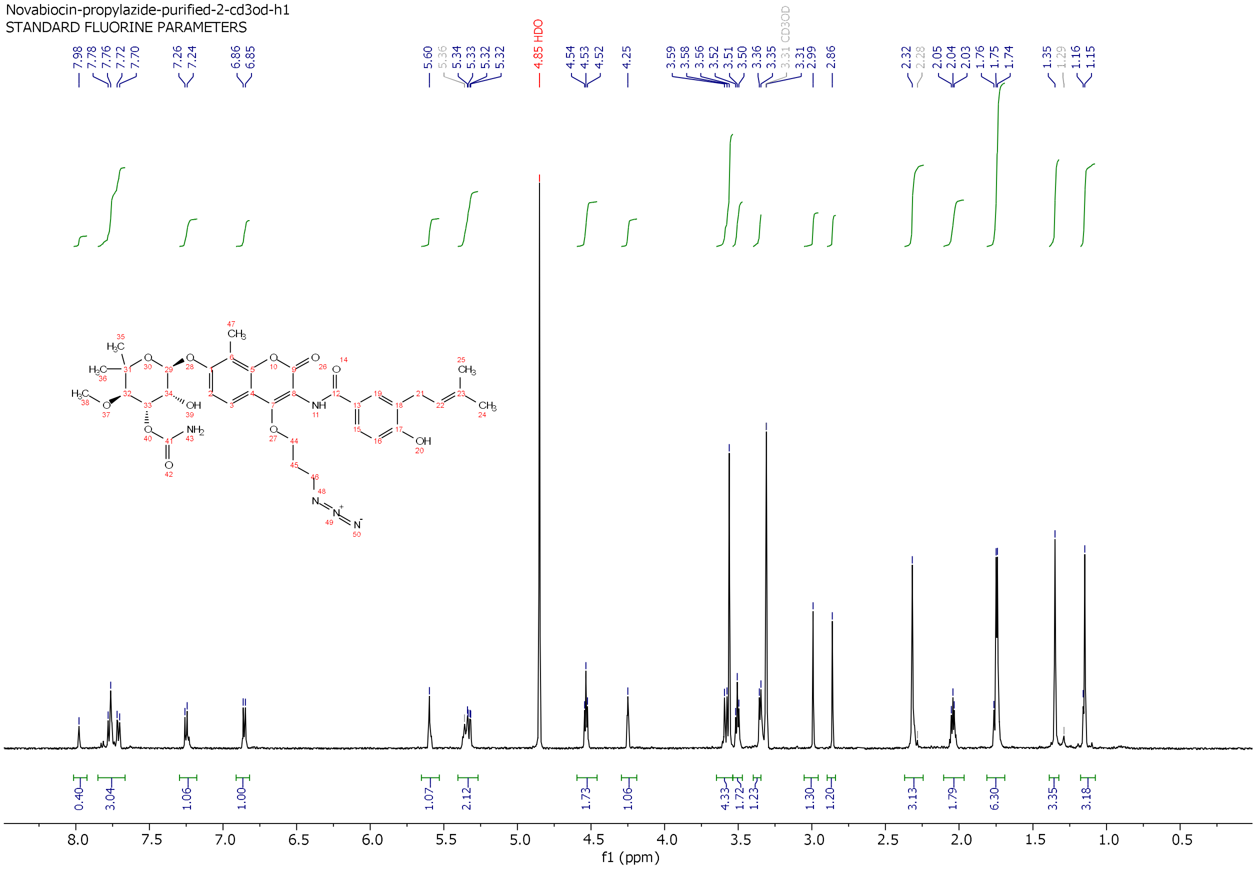


[1]H-NMR of **novAZ1**.

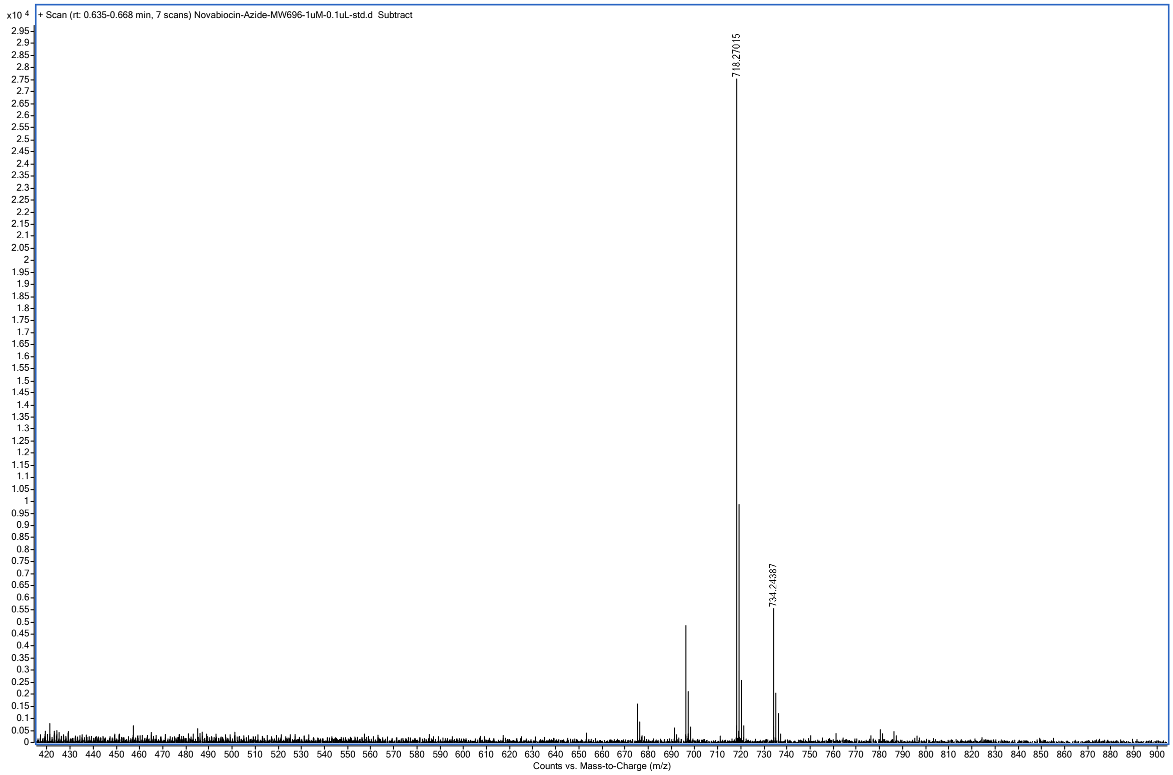


High resolution mass spectrum (ESI) of **novAZ1**.

**Synthesis of purAZ1**.

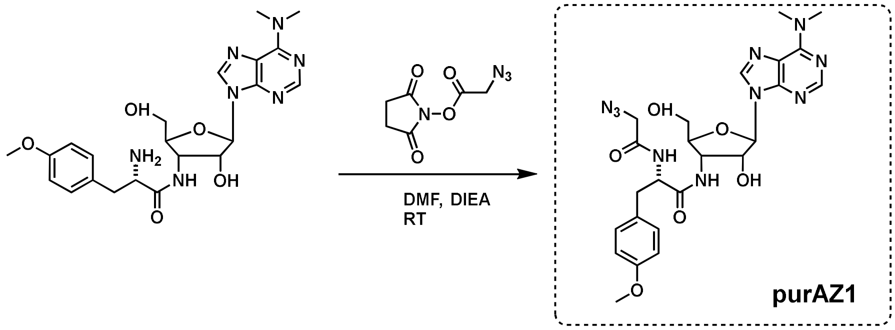


Puromycin (47.1mg, 0.1mmol) dissolved in dry DMF (1 mL), DIEA (50 µL, 37.1 mg, 0.29 mmol) was first added followed by NHS-ester of azido-acetic acid (24.5 mg, 0.12 mmol). The mixture was stirred under nitrogen at room temperature, and the reaction progress was followed by ESI MS analysis, observed m/z 555 for expected product. At the end of 12 h stirring starting material appears to be completely consumed and the new product was formed. The mixture was then diluted with ether (10 mL) to allow oily residue separated out at room temperature supernatant ethereal layer discarded. The oily residue was further triturated with 1 mL ether to remove most of the DMF, and the left-over oily residue was used for further purification. The residue was dissolved first dissolved in HPLC grade acetonitrile (1 mL), followed by dilution with 90:10 ($H_2O$:$CH_3CN$ containing 0.1%TFA, 8 mL). The solution was then subjected to preparative HPLC purification using starting gradient of 10% B ($CH_3CN$ containing 0.1%TFA), and reaching to 100% B in min. The fractions containing desired product were combined, concentrated under reduced pressure using rotary evaporator, the remaining aqueous solution was frozen to -80 $^oC$ and lyophilized to obtain the dry residue. The compound was analyzed by analytical HPLC, characterized by High resolution mass and $^1$H-NMR as follows: HRMS observed m/z 555.2419 $[M+H]^+$ and 577.2243 $[M+Na]^+$; expected m/z 555.2423 and 577.2242 for $C_{24}H_{30}N_{10}O_6$ $[M+H]^+$ and $[M+Na]^+$ respectively. $^1$H NMR (600 MHz, $CD_3OD$) δ 8.55 (brs, 1H), 8.40 (d, J=6 Hz, 1H), 8.34 (s, 1H), 8.23 (d, J= 6 Hz, 1H), 7.79 (d, J= 6 Hz, 1H), 7.79 (d, J= 6 Hz, 1H), 7.39 (t, J= 6 Hz, 1H), 7.31 (t, J= 6 Hz, 1H), 7.16 (d, J= 12 Hz, 2H), 6.88 (d, J= 12 Hz, 2H), 6.27 (brd, J= 3Hz, 1H), 5.74 (dd, J= 6, 3 Hz, 1H), 4.52 (q, J= 6Hz, 1H), 4.37 (d, J= 6 Hz, 1H),

4.24 (brs, 1H), 4.21 (m, 1H), 3.98-4.06 (m, 2H), 3.84 (s, 3H), 3.78 (brs, 6H), 3.46 (brd, J= 6 Hz, 2H), 3.00 (dd, J= 6 Hz, 1H), 2.91 (dd, J= 6 Hz, 1H).

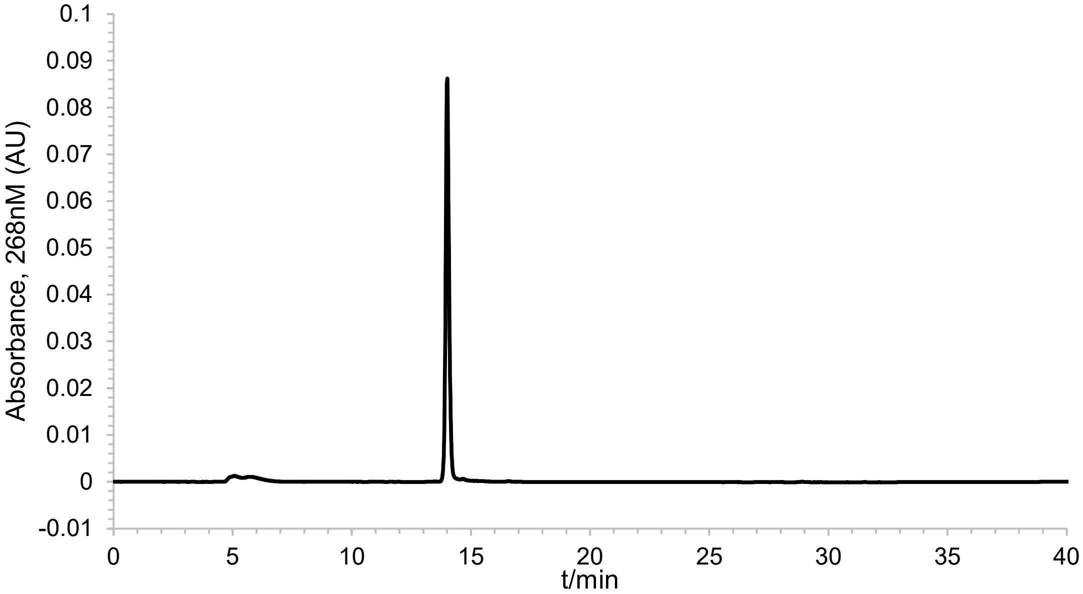


Analytical HPLC chromatogram of **purAZ1**.

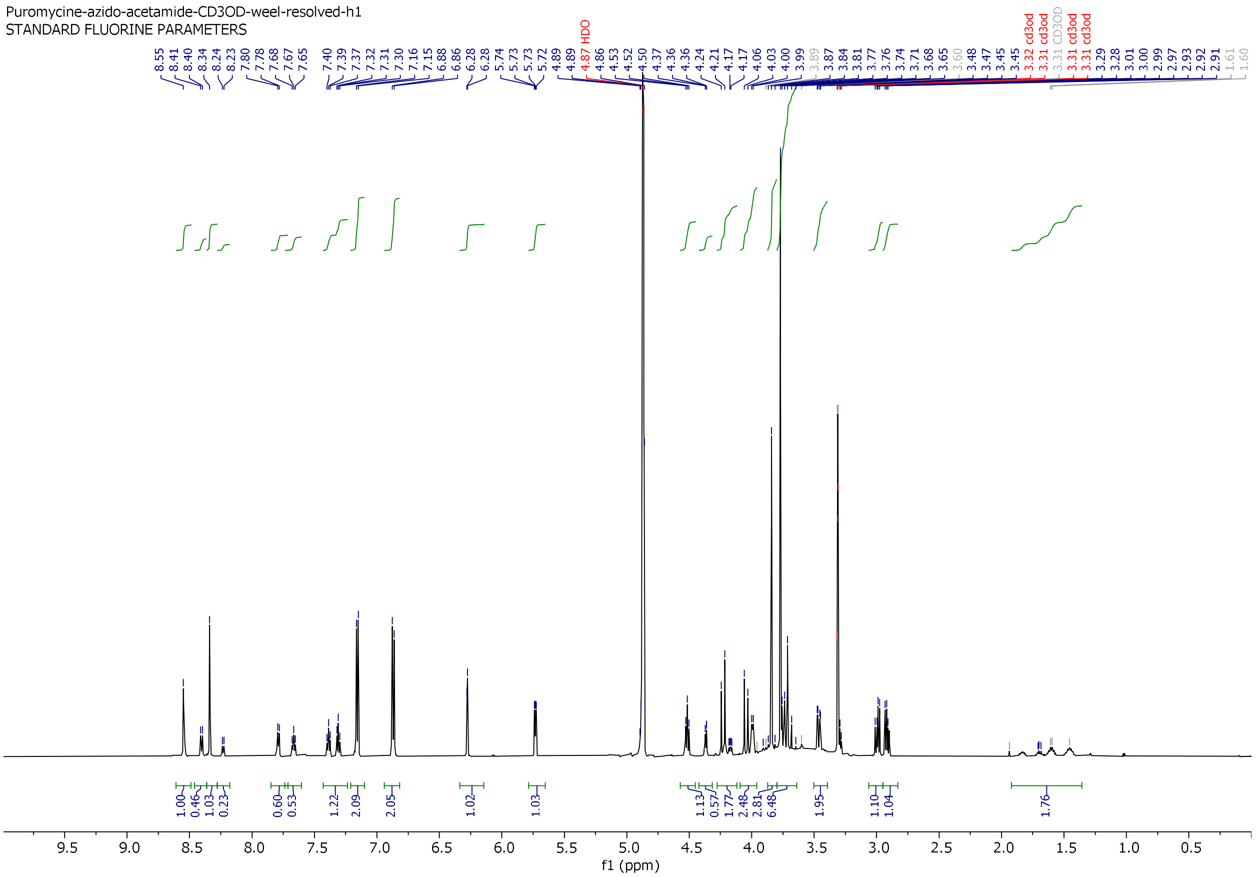


[1]H-NMR of **purAZ1**.

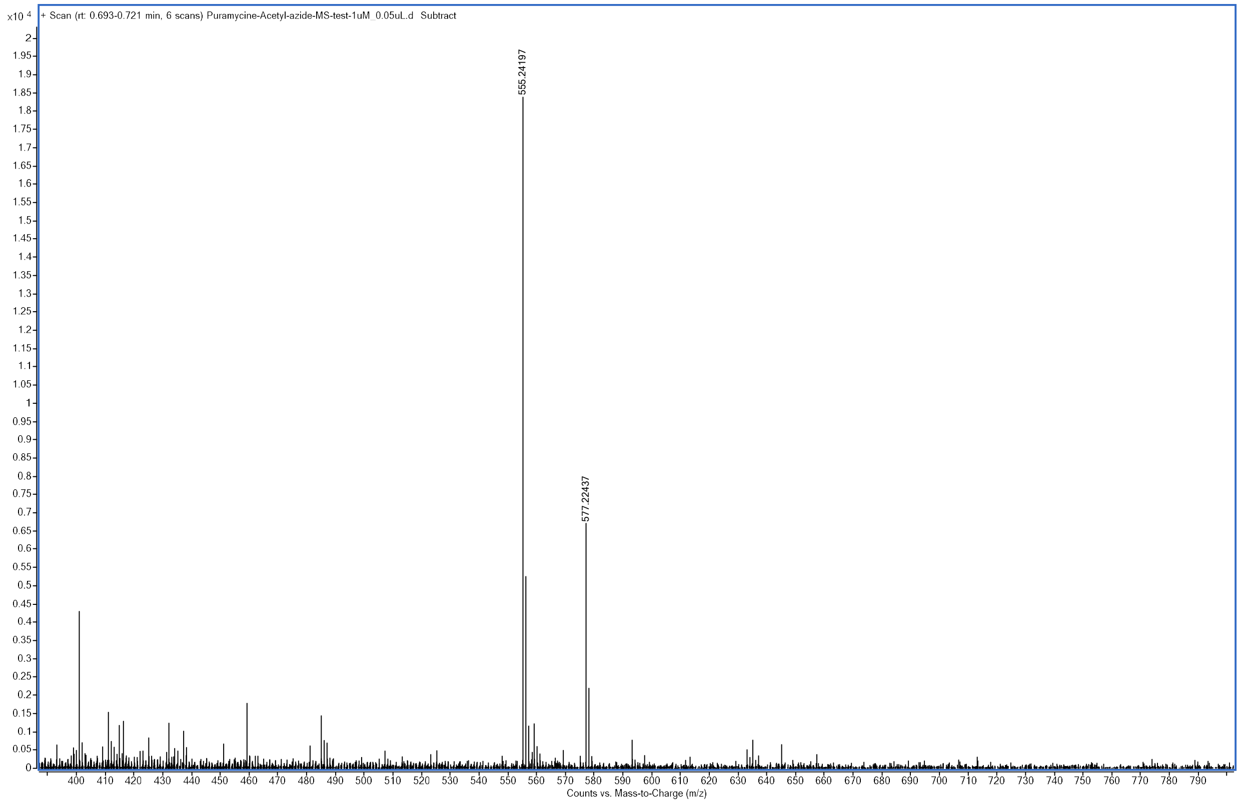


High resolution mass spectrum (ESI) of **purAZ1**.

**Synthesis of zolAZ1.**

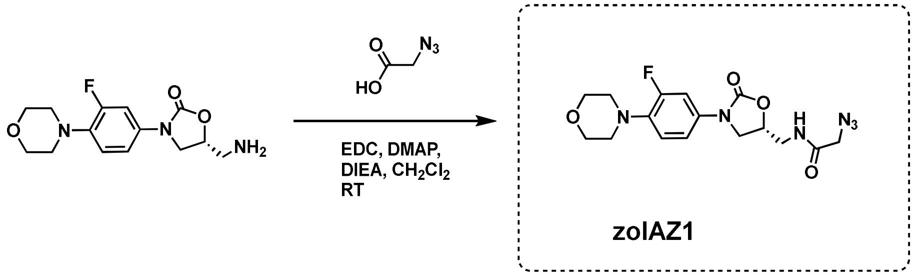


Following a published protocol, **zolAZ1** was synthesized and characterized as follows:[3]

To a solution of amine in $CH_2Cl_2$ (10 mL) were successively added EDCI.HCl (107 mg, 0.56 mmol) and 2-azidoacetic acid 24 (42 µl, 57 mg, 0.56 mmol) were added. The reaction mixture was then stirred for 3 h at 20 °C. The reaction mixture was adsorbed on silica gel and purified on a silica gel column ($CH_2Cl_2$ then $CH_2Cl_2$/EtOH: 96/4) to afford the compound 5 (134 mg, 0.34 mmol, yield: 76%) isolated as a white solid. NMR $^1$H (400 MHz, $CDCl_3$): δ 7.38 (dd, J = 2.6, 14.3 Hz, 1H), 7.05–7.01 (m, 2H), 6.88 (dd, J = 8.8, 9.4 Hz, 1H), 4.78–4.71 (m, 1H), 4.01 (t, J = 8.9 Hz, 1H), 3.96 (s, 2H), 3.84–3.81 (m, 4H), 3.75–3.56 (m, 3H), 3.02–3.00 (m, 4H). NMR $^{13}$C (100 MHz, $CDCl_3$): δ 168.0, 156.6, 156.8, 136.8, 132.9, 119.0, 114.1, 107.7, 71.7, 67.1, 52.5, 51.1, 47.9, 42.2. MS (ESI): m/z 379.2 $[M + H]^+$.

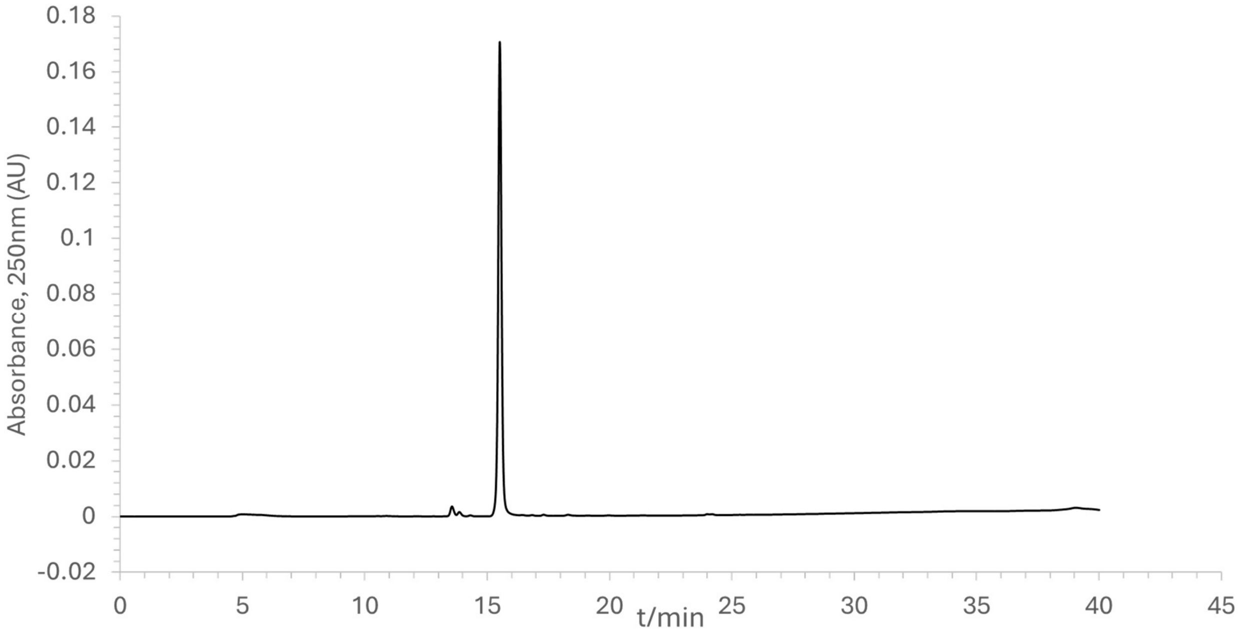


Analytical HPLC chromatogram of **zolAZ1**.

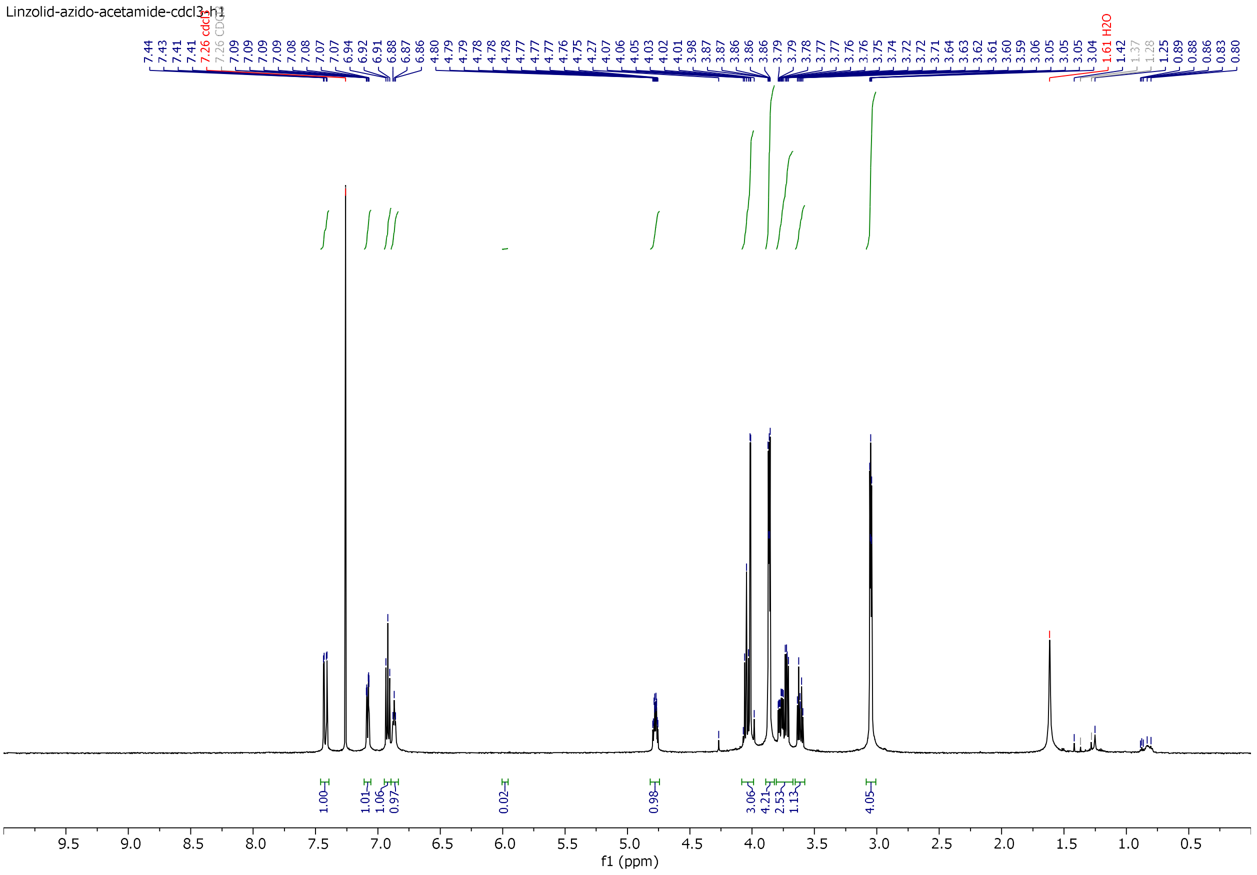

[1]H-NMR of **zolAZ1**.

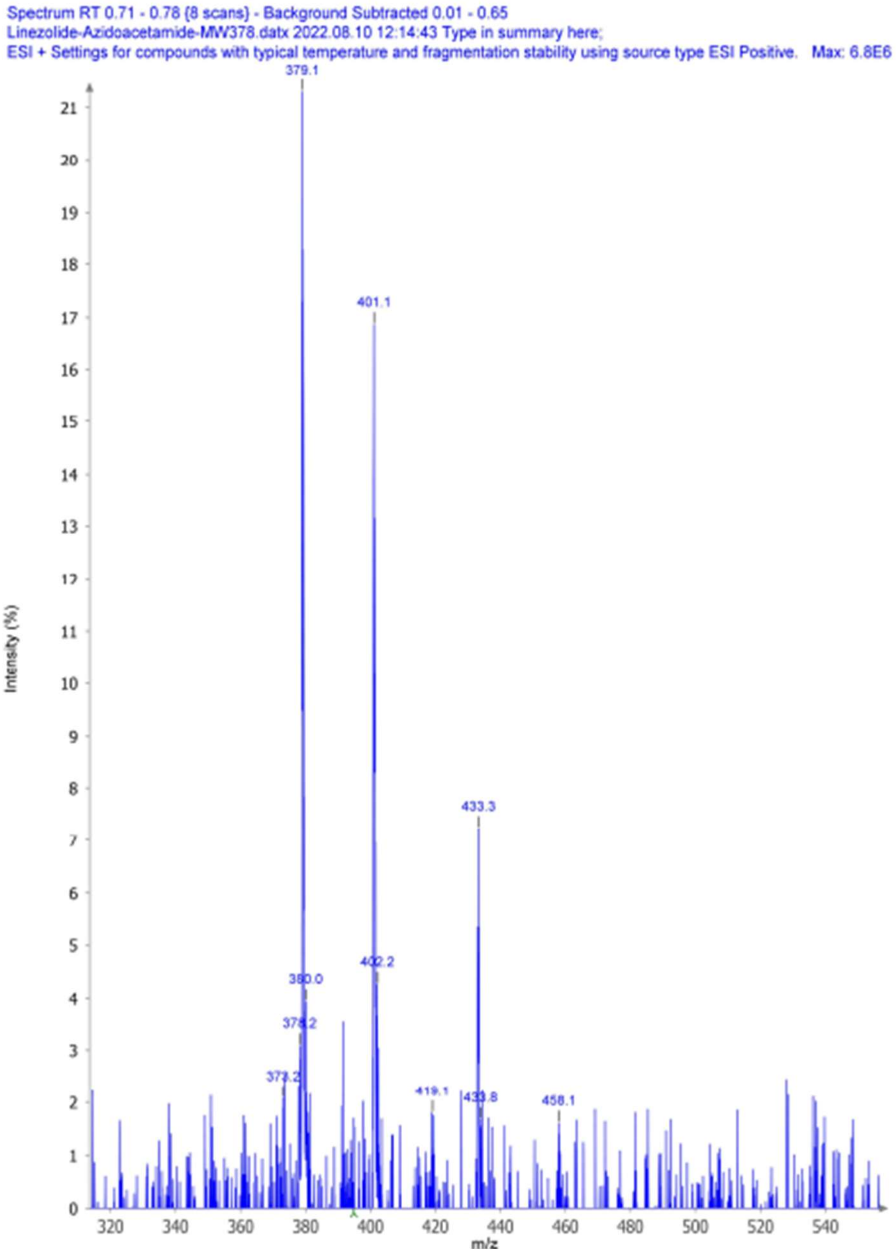


Mass spectrum (ESI) of **zolAZ1**.

**Synthesis of zolAZ2.**

Following a published protocol, **zolAZ2** was synthesized and characterized as follows:[4]

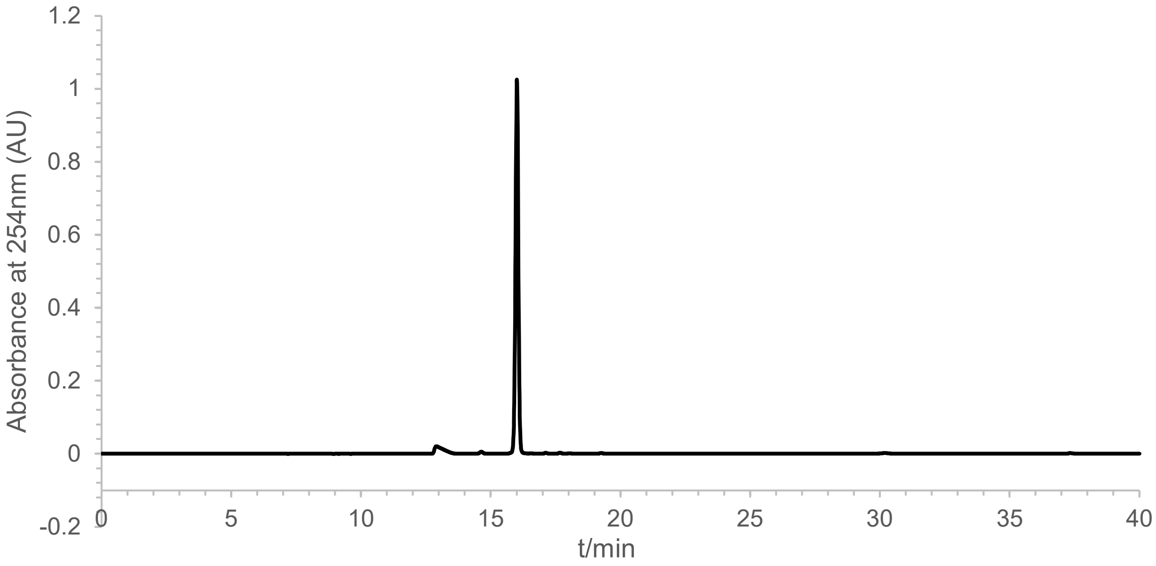


Analytical HPLC chromatogram of **zolAZ2**.

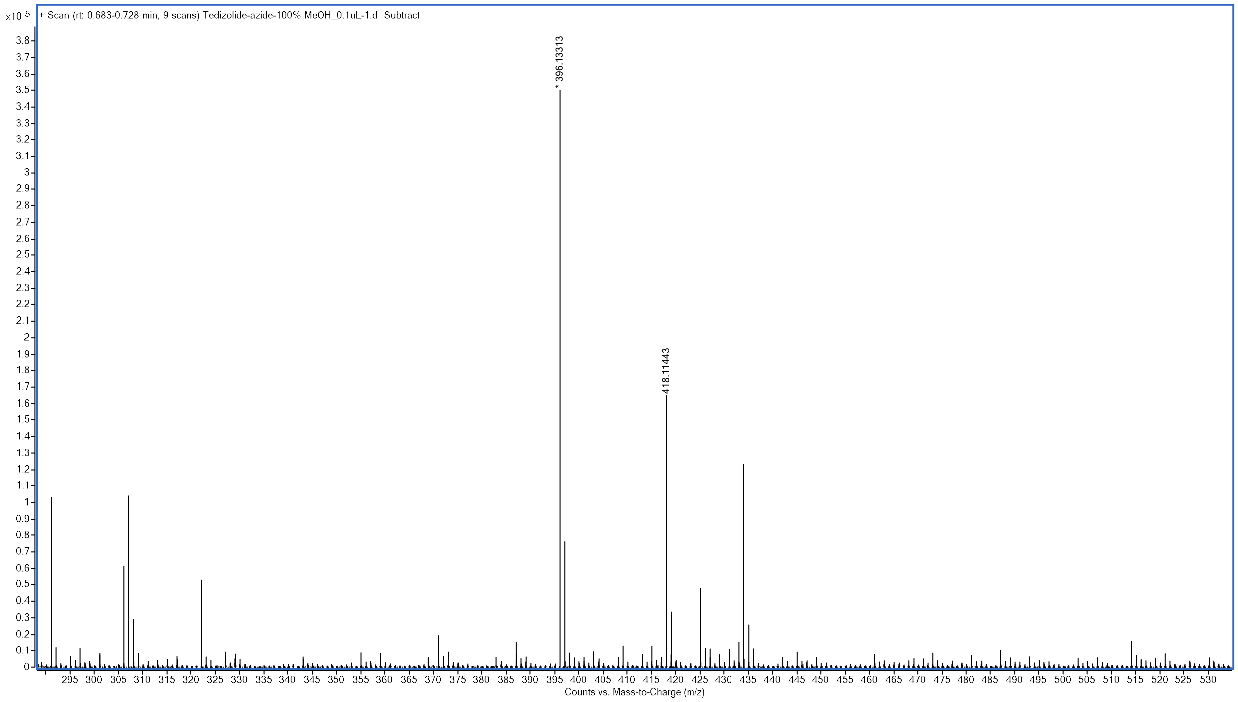


High resolution mass spectrum (ESI) of **zolAZ2**.

**Synthesis of zolAZ3.**

Following a published protocol, **zolAZ3** was synthesized and characterized as follows:[4]

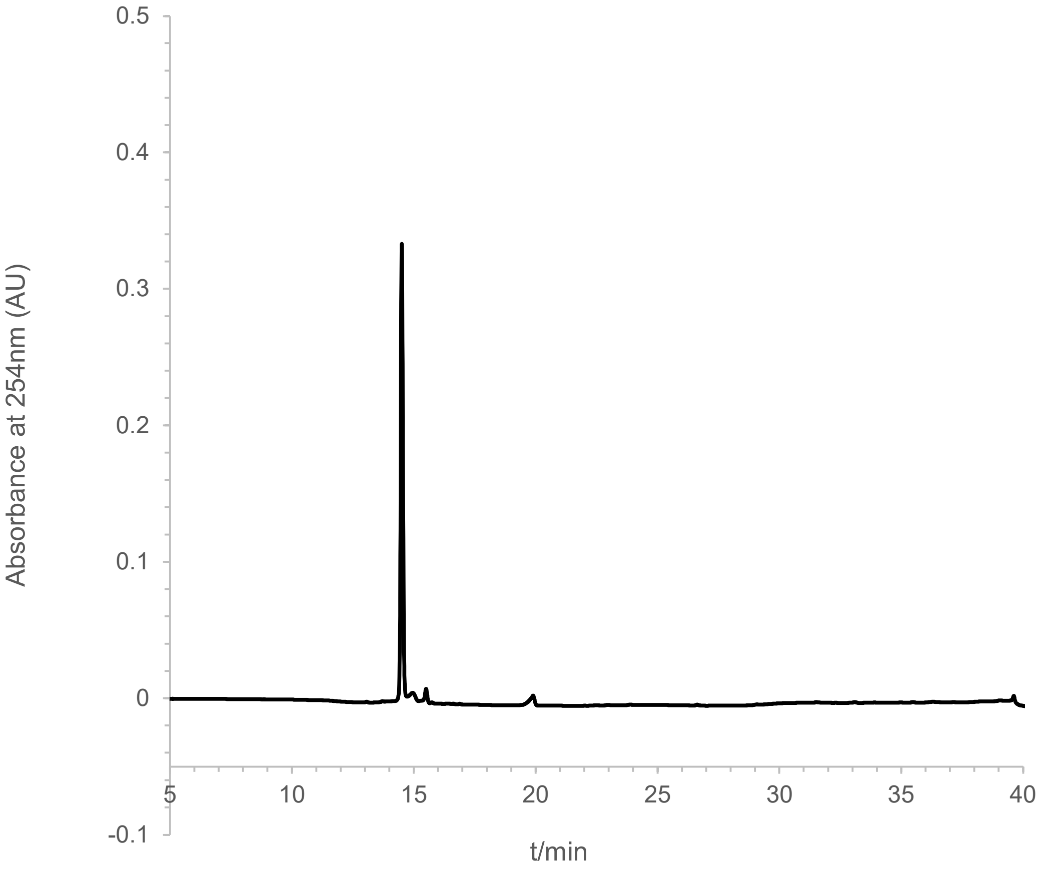


Analytical HPLC chromatogram of **zolAZ3**.

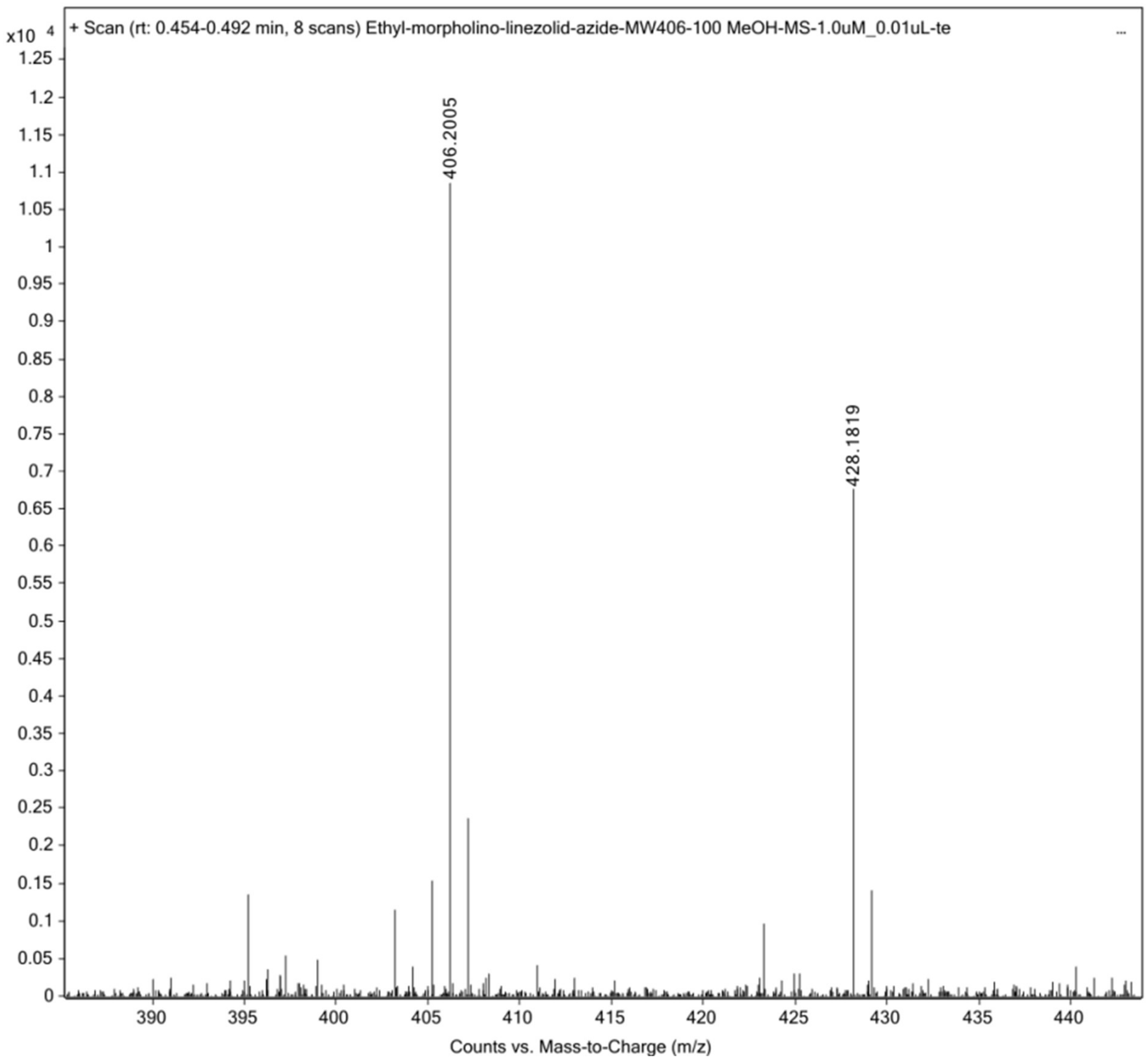


High resolution mass spectrum (ESI) of **zolAZ3**.

**Synthesis of metAZ1.**

Following a published protocol, **metAZ1** was synthesized and characterized as follows:[5]

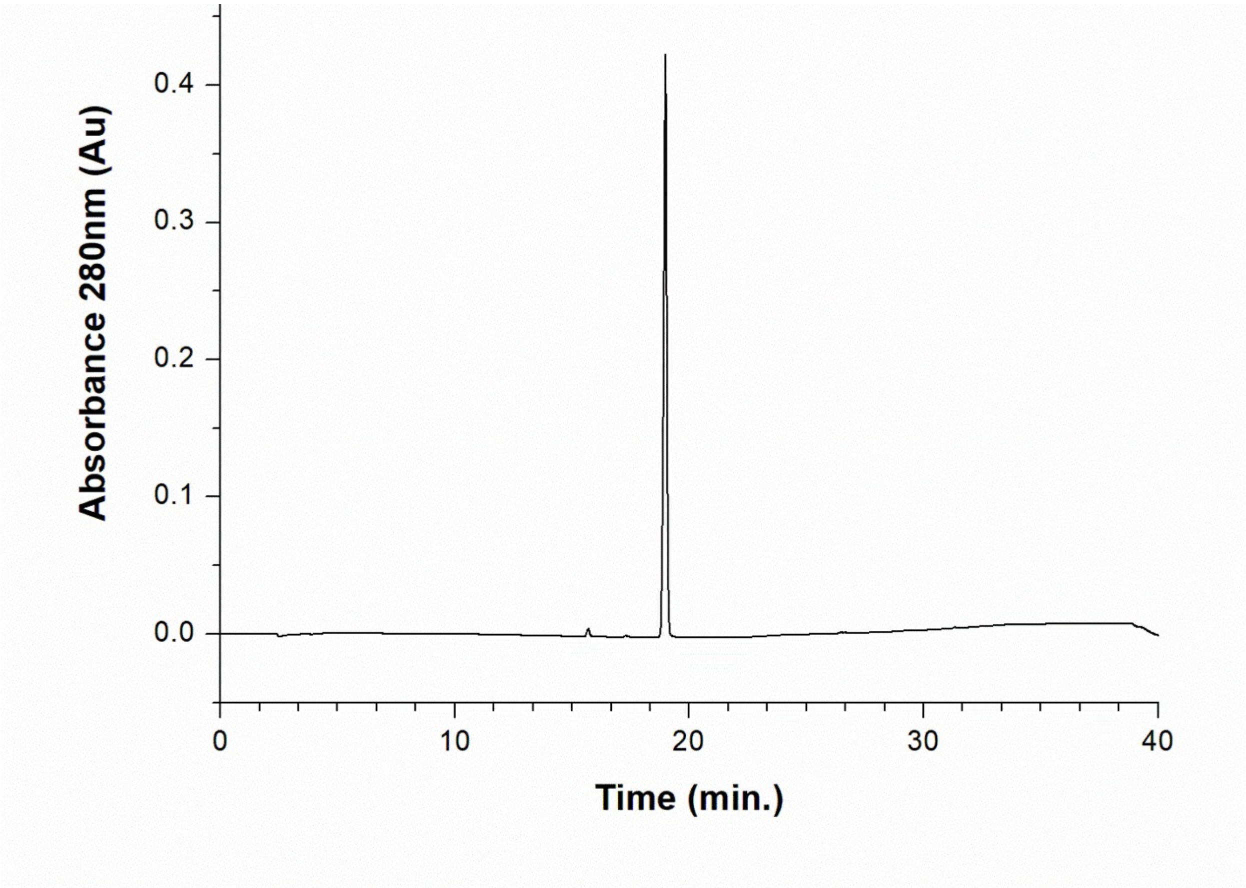


Analytical HPLC chromatogram of **metAZ1**.

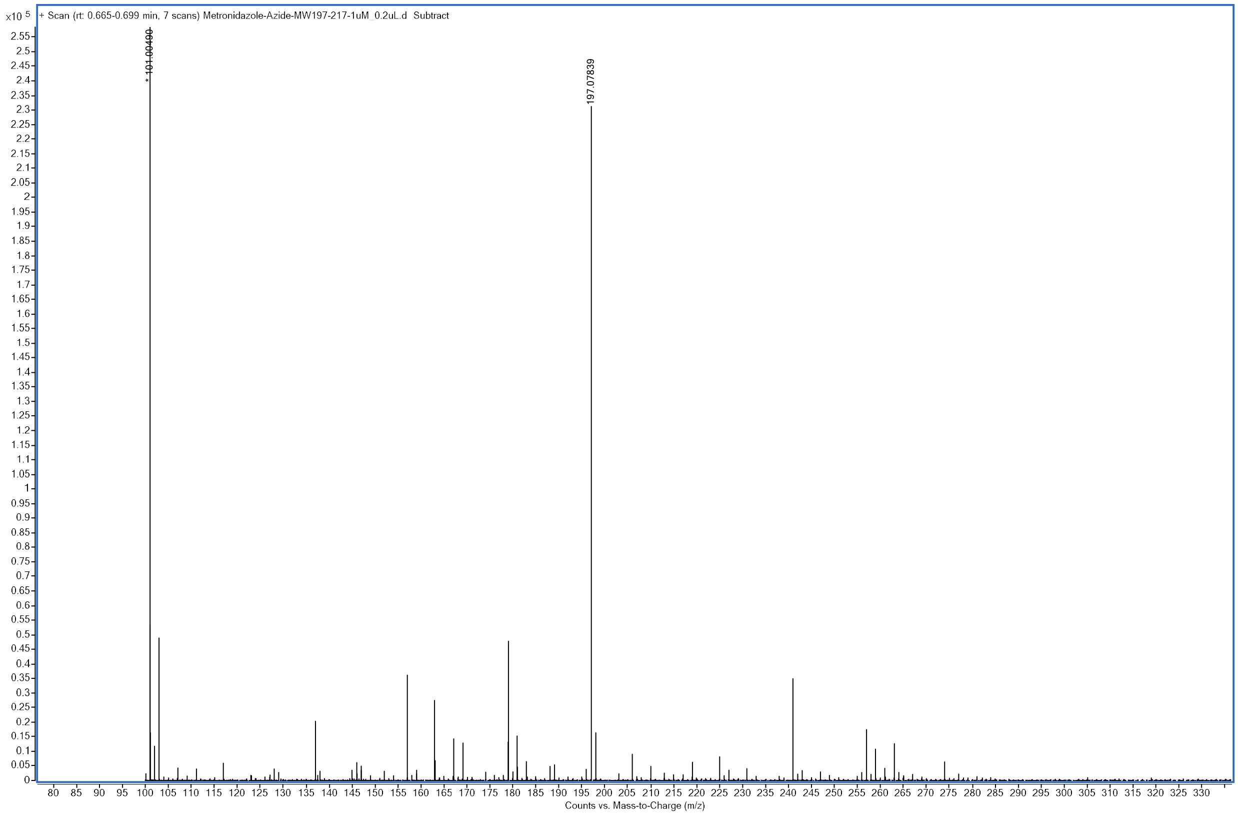


High resolution mass spectrum (ESI) of **metAZ1.**

**Synthesis of sulAZ1.**

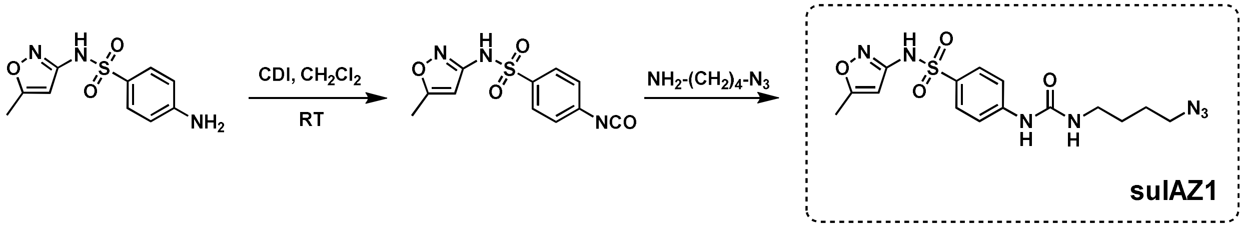


Following a published protocol, **sulAZ1** was synthesized and characterized as follows:[6]

A solution of sulfamethoxazole (51.0 mg, 0.2 mmol) in anhydrous $CH_2Cl_2$ (5 mL) was added CDI (40 mg, 0.25 mmol) at room temperature. The homogenous mixture was stirred overnight (~16 h). Next day 4-azidobutyl amine was added and the mixture was further stirred for 4hrs, TLC and ESI mass analysis indicated the new product formation (non-polar compound compared with starting material). The reaction was quenched with aq. citric acid solution, diluted with 10 mL $CH_2Cl_2$ and organic layer separated. Organic washed with water, brine and dried over $Na_2SO_4$. Concentration under reduced pressure using rotary evaporator followed by silica gel chromatography hexane:ethyl acetate (70:30) yielded the desired compound as solid (61.0mg, 62%). The compound was analyzed by analytical HPLC, characterized only by High resolution mass and used as is: HRMS observed m/z 394.1296 $[M+H]^+$ and 416.1114 $[M+Na]^+$; expected m/z 394.1292 and 416.1111 for : $C_{15}H_{20}N_7O_4S$ $[M+H]^+$ and $[M+Na]^+$ respectively; observed 394.1296. $^1H$ NMR (600 MHz, $CD_3CN$) δ 1.53-1.62 (m, 4H), 2.30 (s, 3H, $CH_3$), 3.18 (q, J= 6Hz, 2H), 3.32 (t, J= 6 Hz, 2H), 6.11 (s, 1H), 7.53 (s, 1H), 7.56 (d, J= 6 Hz, 2H), 7.72 (d, J= 6 Hz, 2H).

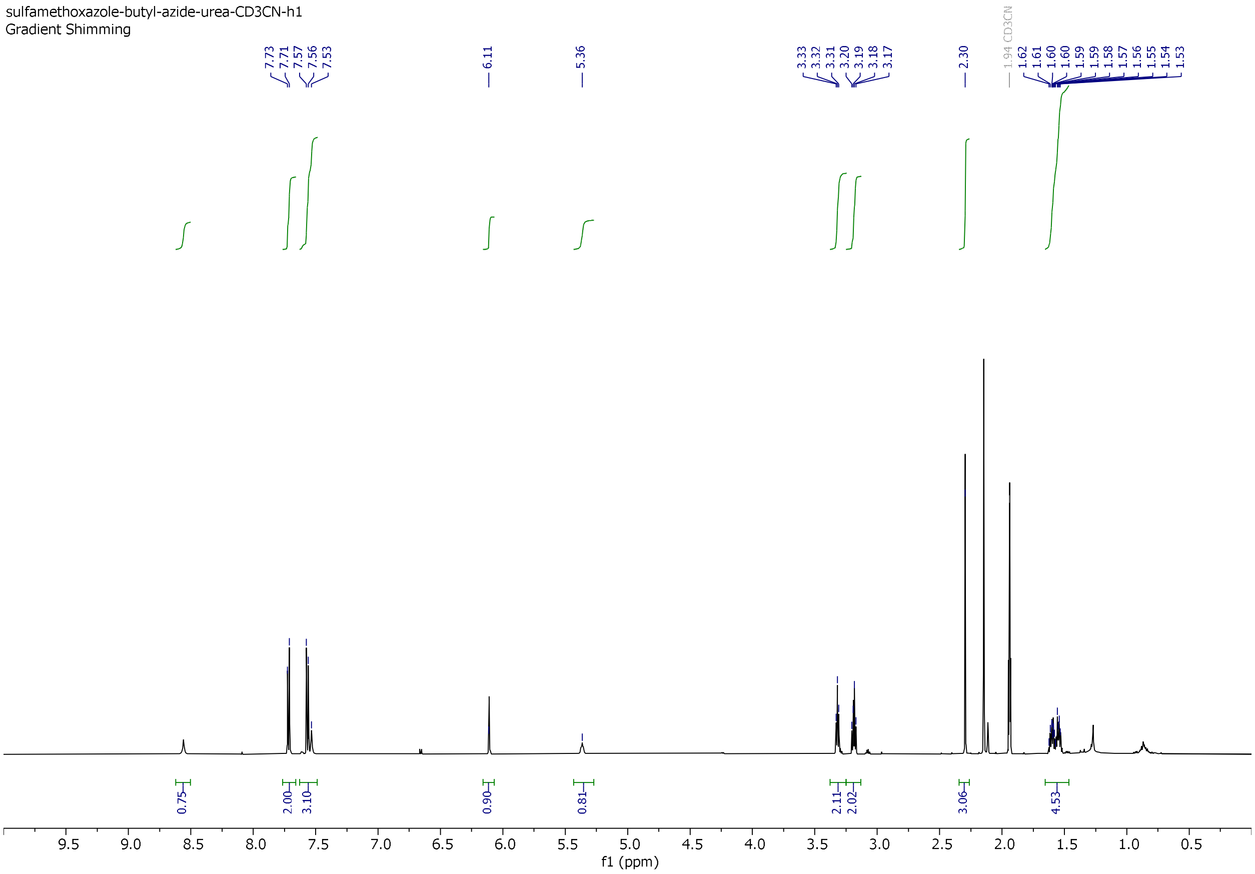


$^1$H-NMR of **sulAZ1**.

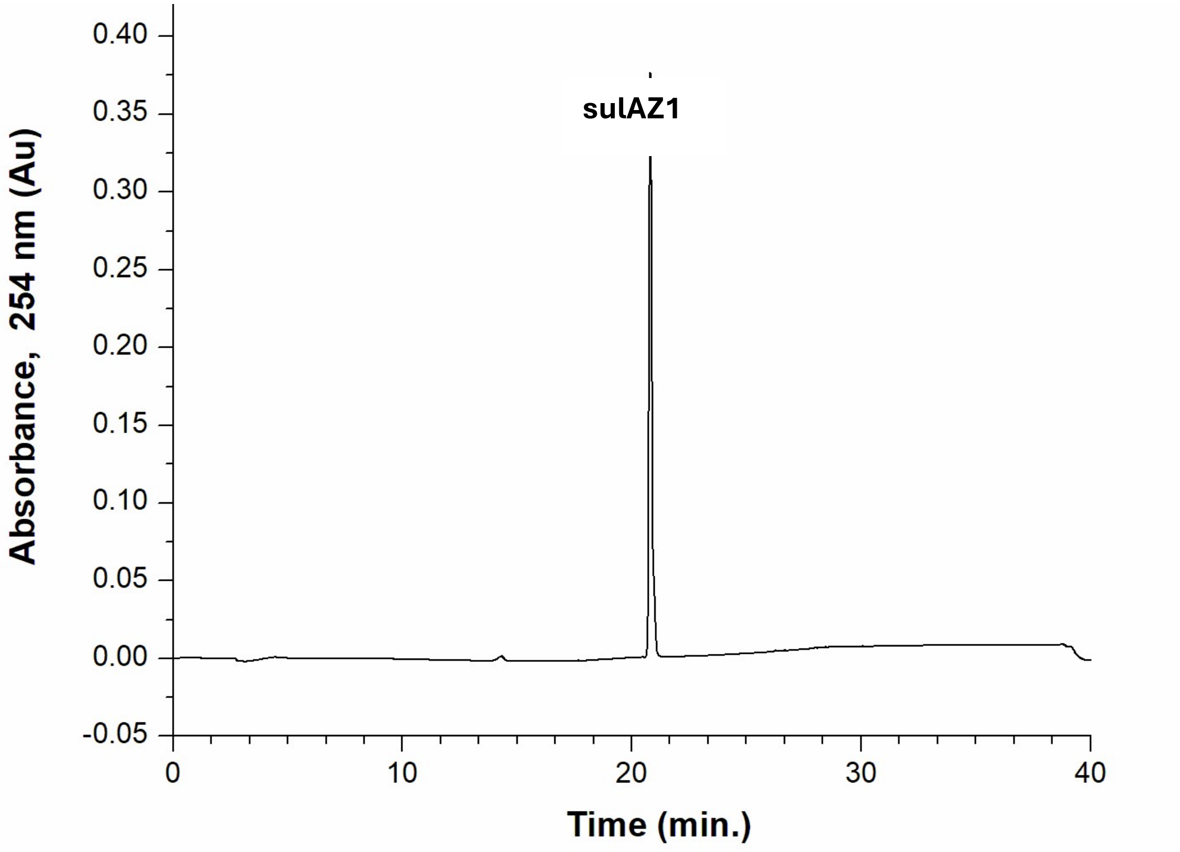


Analytical HPLC chromatogram of **sulAZ1**.

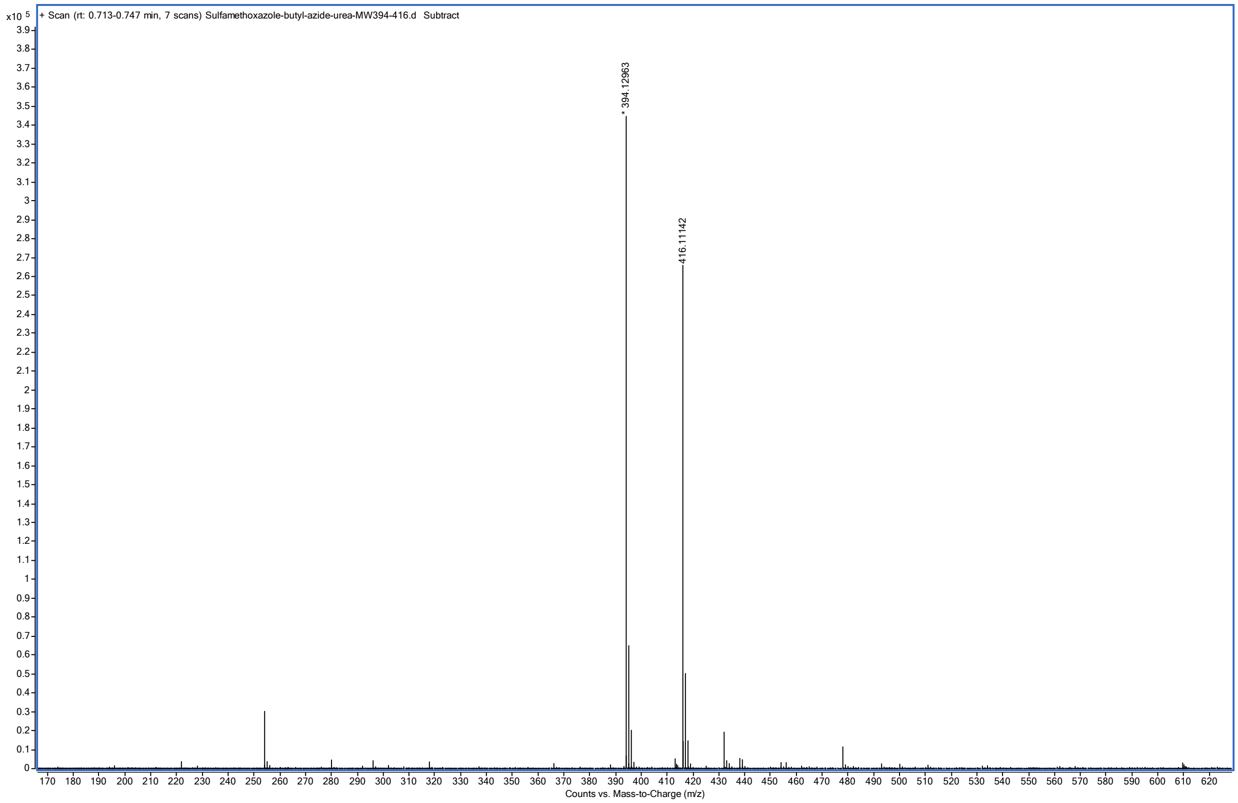


High resolution mass spectrum (ESI) of **sulAZ1**.

### 404 Library of azido-tagged molecules (Enamine/Sigma)

1
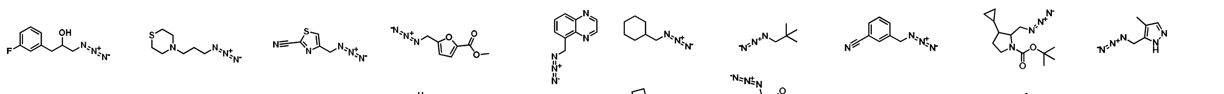

11
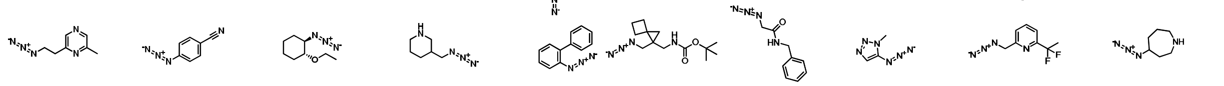

21
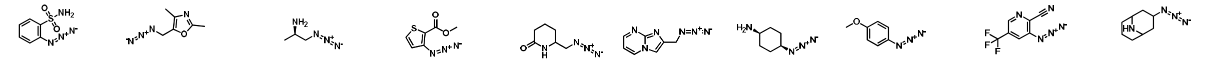

31
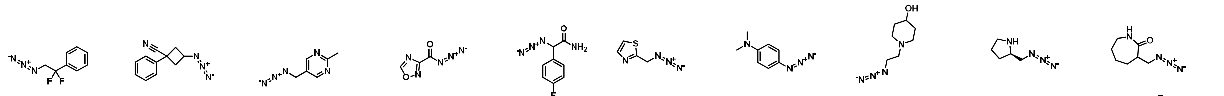

41
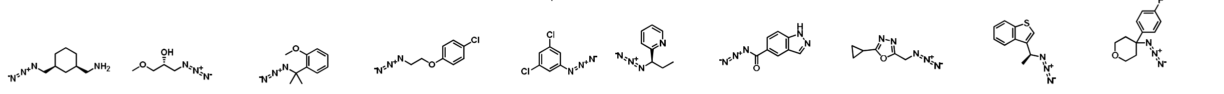

51
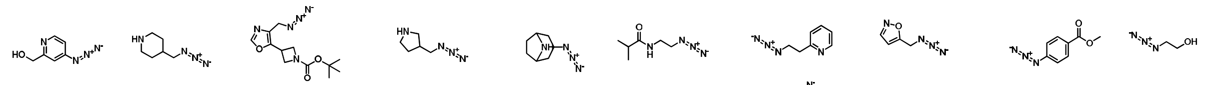

61
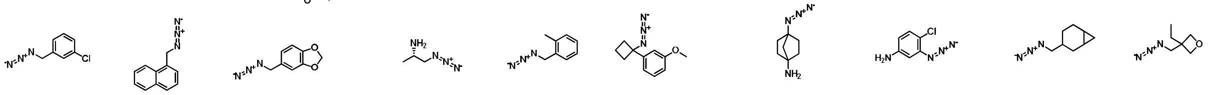

71
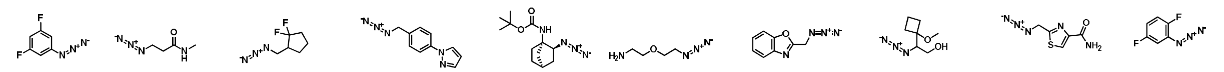

81
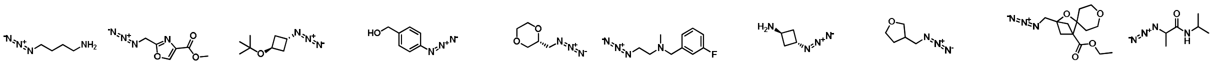

91
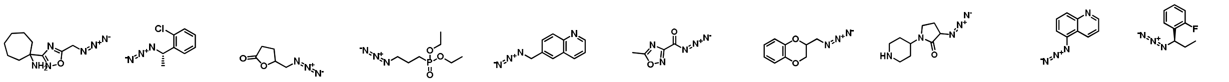

101
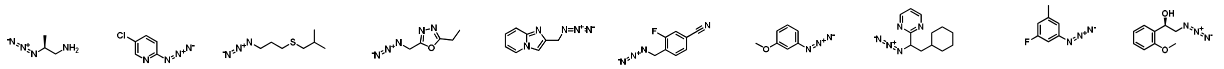

111
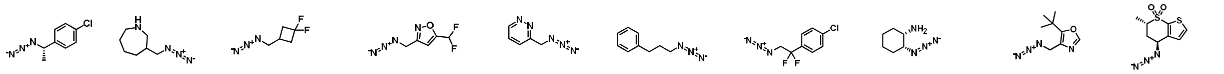

121
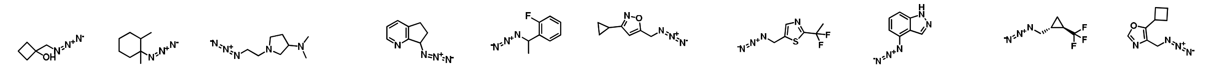

131
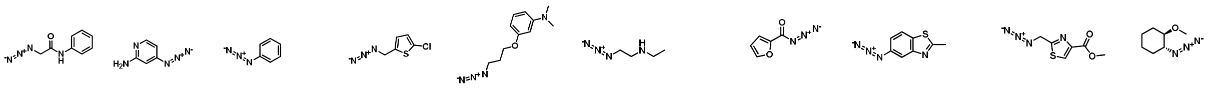

141
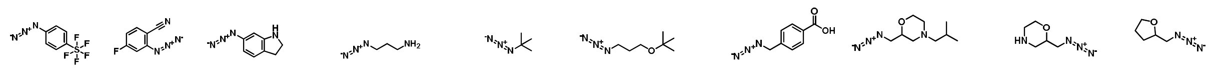

151
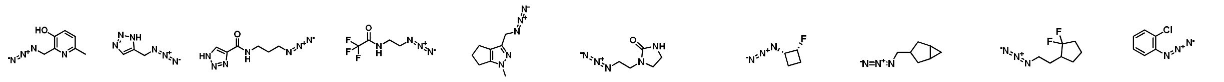

161
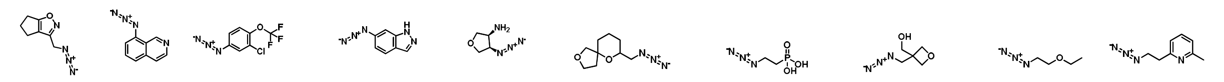

171
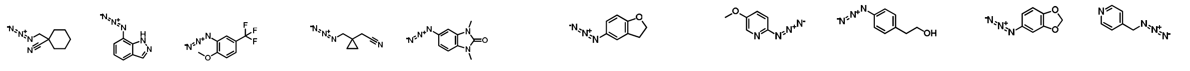

181
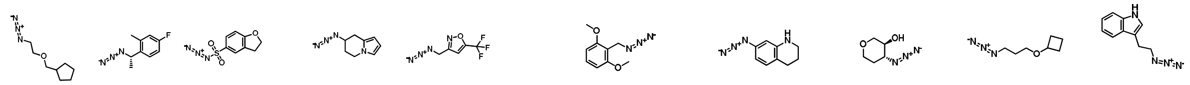

191
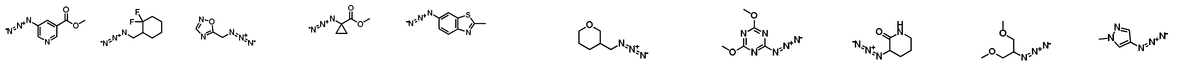

201
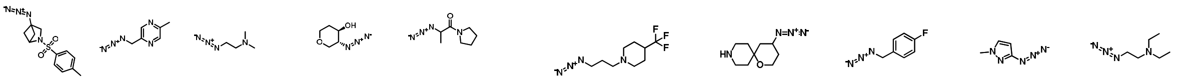

211
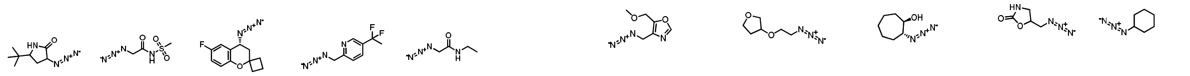

221
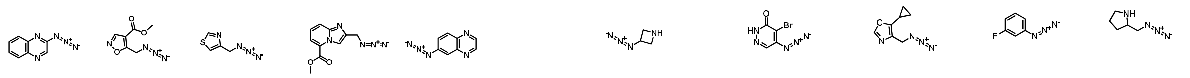

231
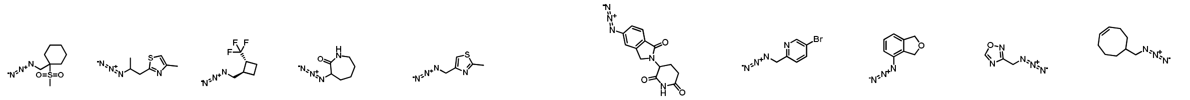

241
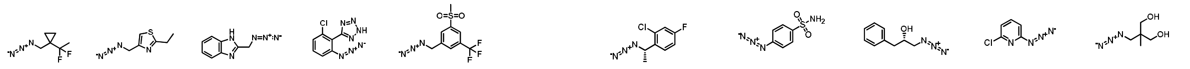

251
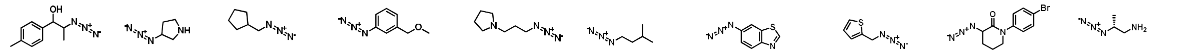

261
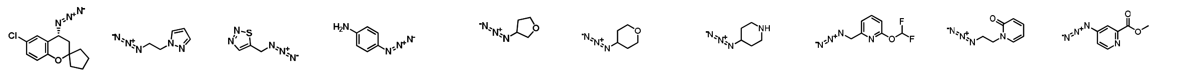

271
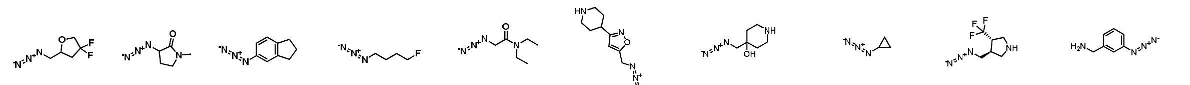

281
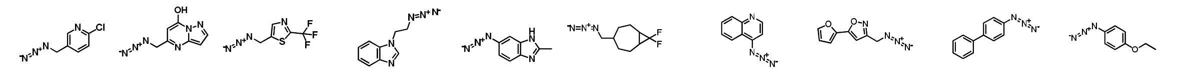

291
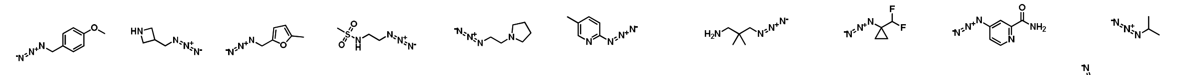

301
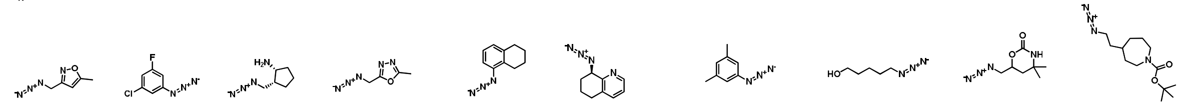

311
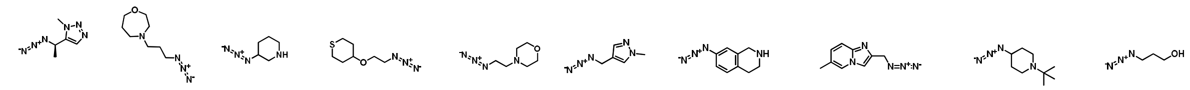

321
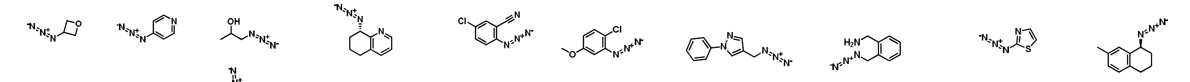

331
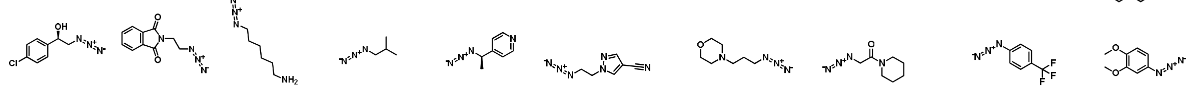

341 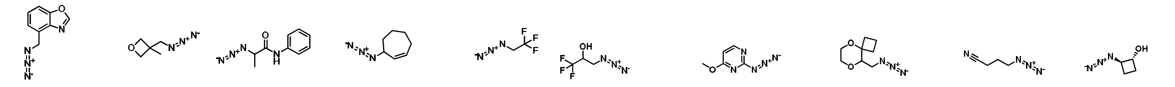

351 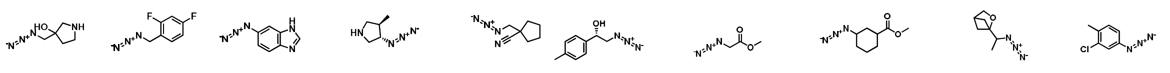

361 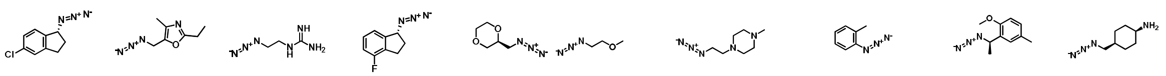

371 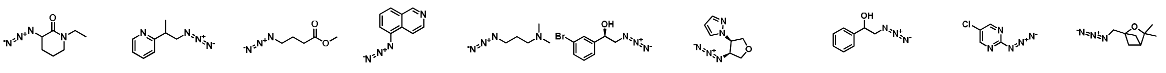

381 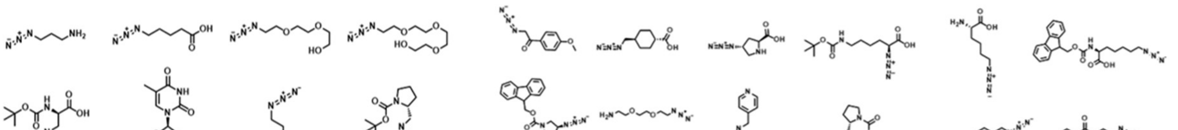

391 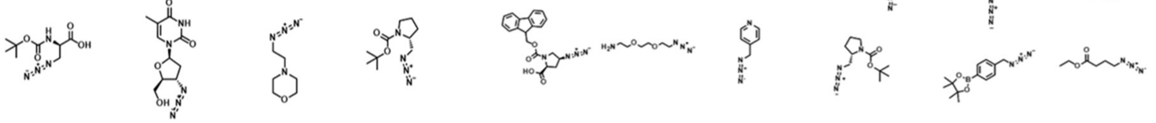

401 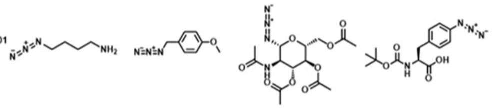
